# Structural basis for catalytic and inhibitory divergence between archaeal and bacterial ammonia monooxygenases

**DOI:** 10.64898/2026.08.31.748207

**Authors:** Xiaoyun Yang, Tie-Qiang Mao, Zhi-Cong He, Yanwei Chen, Guoping Zhao, Peng Jin, Shengying Li, Hong-Po Dong, Wei Peng, Chuanlun Zhang, Zongqiang Li

## Abstract

Ammonia oxidation initiates nitrification and is closely linked to microbial N_2_O production. Ammonia monooxygenase (AMO) catalyzes the first and rate-limiting step of nitrification and is widespread across evolutionarily distinct ammonia-oxidizing archaea (AOA) and bacteria (AOB). The ocean is the largest biome for AOA and AOB, which have distinct ecological niches and markedly different sensitivities to nitrification inhibitors. However, the lack of archaeal AMO structures and inhibitor-bound AMO complexes has hindered mechanistic understanding of the architectural, catalytic, and inhibitory divergence between these two enzyme systems. Here, we report high-resolution cryo-electron microscopy (cryo-EM) structures of marine archaeal AMO captured in active and inactivated states within its native membrane environment, together with inhibitor-bound structures of estuarine bacterial AMO. Archaeal AMO forms an unexpected cup-shaped homotrimer composed of eight subunits per protomer and exhibits substantial architectural divergence from bacterial AMO. Integrated structural, biochemical, kinetic, and computational analyses reveal distinct periplasmic architectures, copper-center organization, and hydrophobic channels between archaeal and bacterial AMOs for ammonium acquisition, catalysis and inhibitor response. These findings provide a structural and mechanistic framework for understanding how archaeal and bacterial AMOs have diverged to distinct ammonia-oxidizing strategies and inhibitor susceptibilities across environmentally important ammonia oxidizers.

## Introduction

The ocean is Earth’s largest biome and plays a central role in the global nitrogen cycle, governing primary productivity, carbon sequestration, and climate regulation ^1,2^. A critical step in this cycle is ammonia oxidation, the first and rate-limiting step of nitrification ^3^. Ammonia oxidation is catalyzed by the integral membrane-bound ammonia monooxygenase (AMO) ^4^, which converts ammonia to hydroxylamine and provides the primary energy source for ammonia-oxidizing microorganisms, including ammonia-oxidizing archaea (AOA), ammonia-oxidizing bacteria (AOB), and complete ammonia oxidizers (comammox) ^5–10^. Beyond its biogeochemical significance, ammonia oxidation is a major biological source of nitrous oxide (N_2_O), a potent greenhouse gas and ozone-depleting substance with a global warming potential approximately 300 times that of CO_2_ over a 100-year timeframe ^11,12^.

AOA predominate across vast oligotrophic ocean regions, accounting for up to 40% of total microbial abundance in marine water columns. Relevant studies indicate that ammonia oxidation accounts for a large fraction of marine N_2_O production, and source-resolved analyses identify AOA as a major and widespread microbial contributor to oceanic N_2_O fluxes ^13,14^. In contrast, AOB are typically enriched in ammonia-rich habitats such as estuaries and coastal waters, where they also contribute significantly to ammonia oxidation and, consequently, substantial N_2_O emissions ^15^. Given the increasing necessity for human intervention in the management of nitrogen levels and greenhouse gas emissions, it is critical to understand how different ammonia oxidizers perform ammonia oxidation across diverse environments. Marine and estuarine environments provide a natural context in which AOA and AOB coexist but often respond differently to ammonium availability and chemical inhibition, making them useful systems for dissecting AMO divergence.

AOA and AOB exhibit distinct physiological characteristics, including differences in substrate affinity and ecological distribution ^16–20^, suggesting potential differences in their underlying enzymatic machinery. Recent studies have determined the structures of bacterial AMOs ^21–23^, whereas complementary biochemical evidence suggests a more elaborate subunit composition for archaeal AMO ^24^. However, understanding of the catalytic mechanism of archaeal AMO has been hindered by the unavailability of an archaeal AMO structure, which limited a unified mechanistic understanding of ammonia oxidation across distinct domains of life.

Another critical knowledge gap is the differential inhibition of AOA and AOB by nitrification inhibitors (NIs). Chemical NIs have long been used as experimental probes to distinguish ammonia-oxidizer activities and, in some settings, to suppress ammonia oxidation ^25–28^. In marine and estuarine studies, allylthiourea (ATU) and acetylene are frequently employed to interrogate ammonia-oxidizer contributions ^29,30^. However, inhibitor responses are strongly compound-, taxon-, and environment-dependent. AOB are often more sensitive to current inhibitors, whereas AOA frequently show weaker or more variable inhibition ^27,31^. This differential sensitivity limits inhibitor-based dissection of ammonia-oxidizer activity and raises a central mechanistic question: how does AMO architecture determine inhibitor susceptibility in archaeal and bacterial ammonia oxidizers?

Here, leveraging cryo-electron microscopy (cryo-EM), we determined structures of marine archaeal AMO captured in active and inactivated states within its native membrane environment. The archaeal enzyme displays an unexpectedly elaborate, cup-shaped architecture and a membrane-embedded organization distinct from bacterial AMO. Combined with inhibitor-bound structures of estuarine bacterial AMO, biochemical data, kinetic analysis, MD simulations, and QM/MM calculations, our work reveals distinct catalytic and inactivation mechanisms in archaeal and bacterial AMOs. These findings provide a molecular basis for understanding AMO diversification across domains of life and offer structural principles for interpreting differential inhibitor response among environmentally important ammonia oxidizers.

### Overall structure of archaeal AMO

To elucidate the long-elusive molecular architecture of archaeal AMO, we cultivated *Nitrosopumilus maritimus* SCM1 (*N. maritimus* SCM1), the first axenically cultured AOA ^6^. Membrane fractions were freshly isolated and subjected to cryo-EM single-particle analysis, culminating in the three-dimensional (3D) reconstruction of *N. maritimus* SCM1 AMO (*Nm*AMO) at a nominal resolution of 2.88 Å (Extended Data Fig. 1a-d and Extended Data Table 1). This map enabled us to unambiguously build an unexpected cup-shaped homotrimer, with each protomer comprising eight distinct subunits that are considerably more intricate than its bacterial counterpart (Fig. 1a-d and Extended Data Figs. 1e and 2). Consistent with recent structural observations of bacterial AMO and pMMO ^22^, *Nm*AMO trimers in isolated membranes assemble into hexagonal arrays (Fig. 1c), suggesting that this evolutionarily conserved supramolecular architecture may be functionally optimized for enzymatic activity ^32^. In addition to the canonical subunits AmoA, AmoB and AmoC, each protomer encompasses three recently predicted components, AmoD (AmoX), AmoF (AmoY), and AmoG (AmoZ) ^24^, a newly identified transmembrane subunit AmoE, and an unknown subunit AmoH. With 51 transmembrane (TM) helices, the archaeal AMO constitutes a slightly larger assembly than bacterial AMO (Fig. 1a-d and Extended Data Fig. 2a-b) ^21–23^. Notably, the single-transmembrane subunit *Nm*AmoG appears to compensate for the absence of a corresponding helix in bacterial AmoB. The first two helices of *Nm*AmoD and *Nm*AmoC closely resemble the six-helix topology of bacterial AmoC, whereas the third helix of *Nm*AmoD shares structural homology with bacterial AmoD. *Nm*AmoB retains only the N-terminal cupredoxin domain, resulting in a flattened extracellular surface relative to bacterial AMO (Extended Data Fig. 2a-c) ^21–23^.

**Figure 1.**
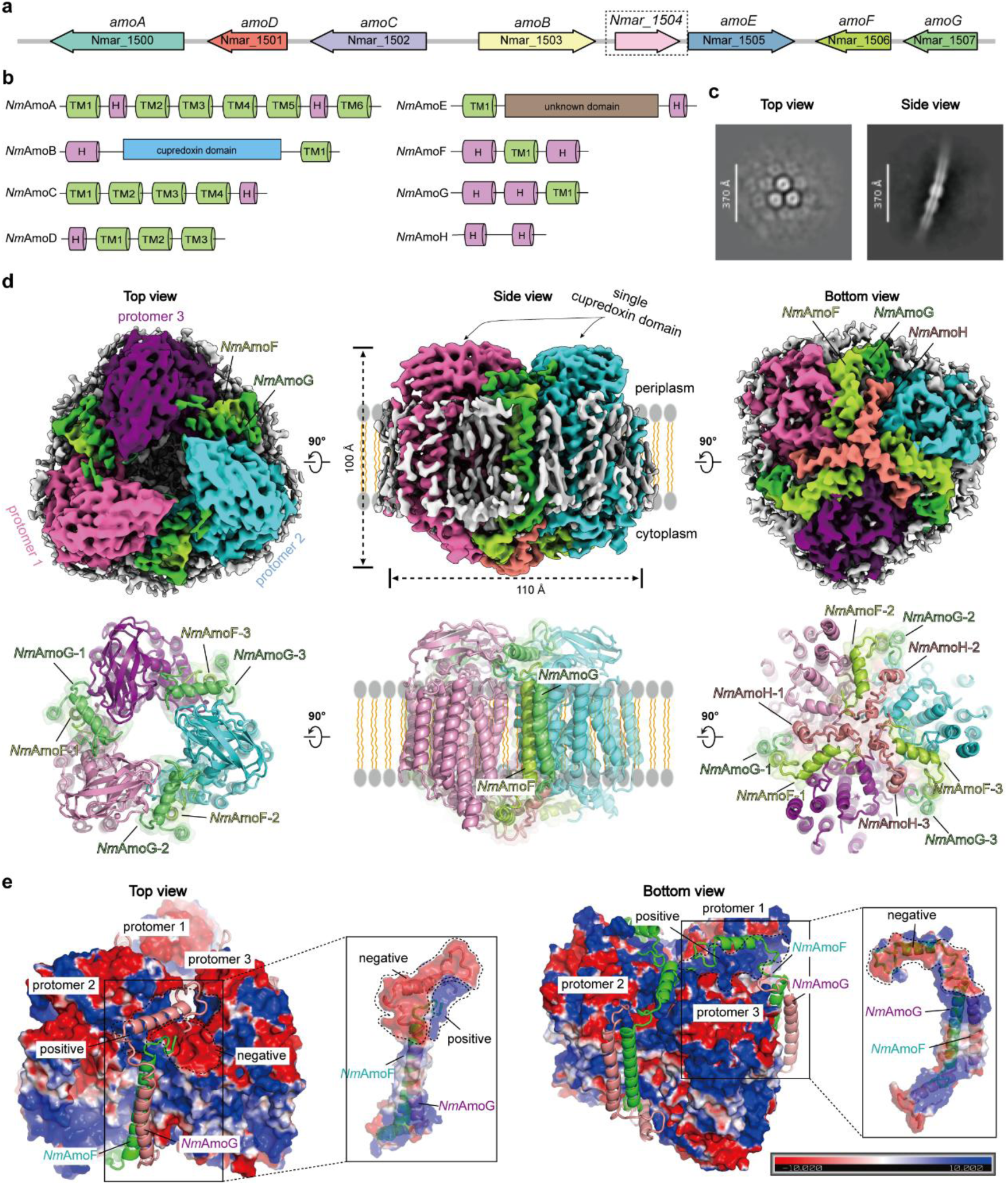
Composition and structure of the archaeal AMO holoenzyme. **a.** Genetic organization of *amo* gene cluster in *N. maritimus* SCM1. Genes are depicted as colored arrows indicating the direction of transcription, with corresponding protein names (*Nm*AmoA-*Nm*AmoG) labeled above. *Nmar*_1504, which is not incorporated into the *Nm*AMO complex, is represented by a dashed box. **b.** Schematic representations of the *Nm*AMO subunit structures. “H” and “TM” denote helical regions and transmembrane helices, respectively. **c.** Representative 2D class averages of *Nm*AMO in native membranes showing the hexagonal arrays of *Nm*AMO trimers. Top (Left) and side (Right) views are shown. **d.** Cryo-EM density map (upper) and corresponding atomic model (lower) of the *Nm*AMO complex. Three views are presented, with individual protomers and subunits color-coded. **e.** The *Nm*AmoFG heterodimer primarily interacts with the *Nm*AMO complex via strong electrostatic interactions. Electrostatic surface potential maps of the *Nm*AMO complex (in the absence of *Nm*AmoFG heterodimers) and the *Nm*AmoFG heterodimer are displayed. Positively (blue) and negatively (red) charged regions involved in the interaction are outlined by dotted lines.

Structural divergence is also evident in the newly identified subunit *Nm*AmoE, which features a single TM helix and a C-terminal domain of unknown function that are absent in the soluble bacterial counterpart (Extended Data Fig. 2d-f). Furthermore, *Nm*AmoE exhibits low evolutionary conservation and appears specific to marine AOA lineages (Extended Data Fig. 3). The stabilization of this massive assembly of *Nm*AMO is mediated by accessory subunits, namely *Nm*AmoF, *Nm*AmoG and *Nm*AmoH. *Nm*AmoF and *Nm*AmoG form a clamp-like heterodimer positioned between adjacent *Nm*AMO protomers, serving as a molecular scaffold (Extended Data Fig. 4). Charged residues at their termini establish a network of electrostatic interactions that reinforce *Nm*AMO trimer integrity (Extended Data Figs. 1e and 4). *Nm*AmoH, enriched in flexible loops, participates in intra- and inter-protomer contacts and assembles into a central propeller-shaped trimer, closing the cytoplasmic base of the cup-shaped architecture. This contrasts sharply with the open conformation observed in bacterial AMO (Extended Data Figs. 1d and 4b). Moreover, the trimeric interface of *Nm*AmoH appears to form a layered, hierarchical channel (Extended Data Fig. 5), suggesting a potential role in regulating material transport or enzymatic dynamics.

Together, these structural analyses reveal fundamental compositional and architectural differences between archaeal and bacterial AMO, providing a framework for understanding their distinct catalytic mechanisms.

### Divergent ammonium capture in archaeal and bacterial AMOs

Previous data have indicated that AOB are typically dominant in environments with high-ammonium concentrations, whereas AOA prevail in oligotrophic, low-ammonium habitats such as the deep ocean, with a significantly higher substrate affinity than AOB ^16^. Given that AMO is essential for both the survival and metabolic activity of ammonia-oxidizing microorganisms, it represents a compelling molecular target for elucidating the mechanistic basis of their divergent ammonium acquisition strategies. Accordingly, we investigated whether structural differences between archaeal and bacterial AMO isoforms underlie their distinct kinetic and physiological behaviors in ammonium capture.

A major difference lies in the extracellular cupredoxin architecture. Genomic analysis shows that *Nmar*_1504, located within the archaeal *amo* gene cluster, is not incorporated into the AMO complex (Fig. 1). Instead, it appears to have been evolutionarily repurposed into a separate, soluble β-barrel protein. Structurally, it closely resembles the C-terminal cupredoxin domain of bacterial AmoB and shares a similar electronegative surface (Fig. 2a). In bacterial AMO, this domain considerably expands the electronegative extracellular surface, likely facilitating the recruitment of cationic ammonium ^23^. Its absence in archaeal AMO reduces the electronegative periplasmic surface area by ∼50% compared to that of bacterial AMO (Fig. 2b). Consistent with this observation, the purified cupredoxin domain of *Nm*AMO exhibits a comparable ∼30% reduction in ammonium-binding capacity relative to that of *N. halophila* AMO (*Nh*AMO) (Fig. 2c), suggesting that bacterial AMO may possess a greater ammonium-capture capacity. Conversely, the absence of *Nmar*_1504 enhances local periplasmic electronegativity of *Nm*AMO, a change that is consistent with the higher ammonium affinity of cupredoxin domain of *Nm*AMO, as measured by ITC (Fig. 2d). *Nmar*_1504 is phylogenetically conserved in marine AOA, but absent in terrestrial AOA lineages (Extended Data Fig. 3). Pull-down assays showed that the *Nmar*_1504-encoded protein interacts with the cupredoxin domains of *N. maritimus* SCM1 and *Nitrososphaera viennensis* (a representative terrestrial AOA) (Fig. 2e), suggesting that absence of *Nmar*_1504 homologs in archaeal AMOs may represent an evolutionary adaptation enabling AOA to maintain efficient ammonia oxidation under nutrient-limited habitats. Thus, archaeal and bacterial AMOs appear to emphasize different ammonium-recruitment strategies: high-affinity binding in *Nm*AMO versus higher-capacity ammonium capture in *Nh*AMO.

**Figure 2.**
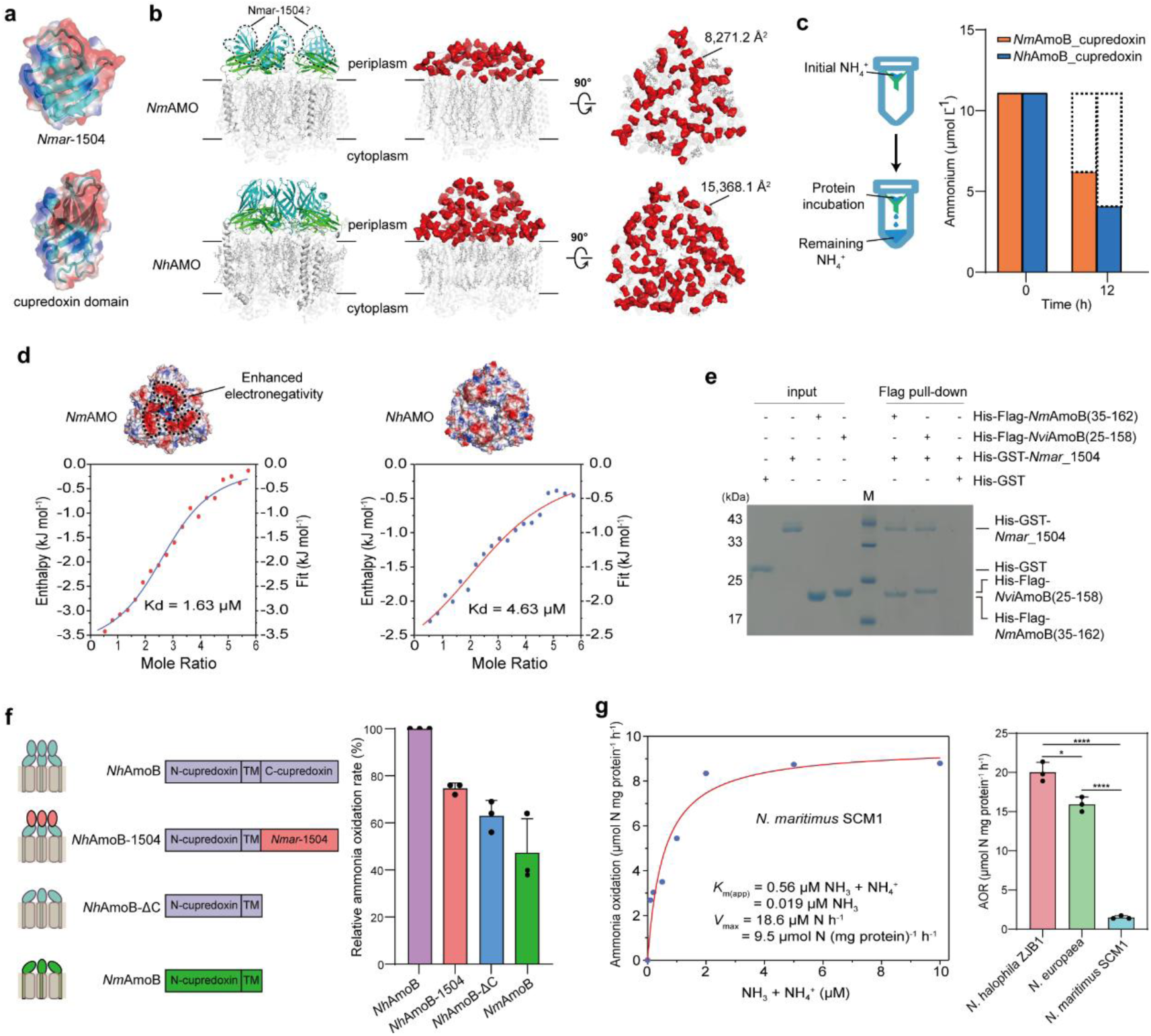
Structural insights into ammonium-capture differentiation of AOA and AOB. a. Structural comparison and electrostatic surface potentials of the C-terminal cupredoxin domain of *Nh*AmoB and *Nmar*_1504-encoded protein. **b.** Electrostatic surface potentials of *Nm*AMO and *Nh*AMO. The lack of *Nmar*_1504-encoded protein reduces negatively charged surface on the periplasmic surface of *Nm*AMO. **c.** NH_4_^+^ binding-capacity assay for purified *Nm*AmoB and *Nh*AmoB cupredoxin domains. **d.** Comparison of the periplasmic negative charge intensities and ITC analysis of NH_4_^+^ binding of *Nm*AMO and *Nh*AMO. **e.** Flag pull-down assays examining interactions between *Nmar*_1504-encoded protein and cupredoxin domains from archaeal AMOs. The pull-down assays employed Flag resin, together with His-Flag-tagged *Nm*AMO (35-162) and *Nvi*AMO (25-158) as the bait proteins. **f.** Functional effects of cupredoxin-domain replacement or deletion in *Nh*AMO. Schematic representation of wild-type *Nh*AmoB and engineered cupredoxin-domain variants (left), together with their relative ammonia oxidation rates (right). **g.** Ammonia oxidation kinetics of representative AOA and AOB. Michaelis–Menten kinetic analysis of ammonia oxidation by *N. maritimus* SCM1 (left) and AMO abundance-normalized ammonia oxidation rates (AOR) of *N. halophila* ZJB1 (co-occupied Cu_C_-Cu_D_ center), *N. europaea* (Cu_C_ center), and *N. maritimus* SCM1 (Cu_D_ center) (right). Data represent mean ± SD (n = 3). **p < 0.01, ****p < 0.0001 (one-way ANOVA with Tukey’s test).

We further examined whether this extracellular architecture contributes to ammonia oxidation activity. Using the genetic manipulation system established for *N. halophila* ZJB1 ^23^, we engineered AOB variants in which the periplasmic domains of *Nh*AMO were replaced, deleted, or swapped with the corresponding archaeal domain. These modifications progressively reduced ammonia oxidation activity compared with wild-type *Nh*AMO (Fig. 2f), suggesting a functional role for the extracellular cupredoxin module in efficient ammonium acquisition and catalysis. Supporting this inference, kinetic measurements of *N. maritimus* SCM1 exhibit a significantly lower apparent Michaelis constant (*K*_m(app)_) for total ammonium compared with those of the examined AOB strains, indicating its higher substrate affinity. In contrast, the AOB strains displayed higher maximal reaction velocities (*V*_max_) and greater ammonia oxidation rates (AOR) (Fig. 2g and Extended Data Fig. 6). In addition, microcosm enrichment experiments demonstrate that low-ammonium conditions selectively favor AOA proliferation, whereas high-ammonium conditions markedly enhance AOB growth (Extended Data Fig. 7)—thereby corroborating the well-documented niche-associated differences between AOA and AOB.

Collectively, the structural and functional divergence in extracellular domains between archaeal and bacterial AMOs resolves a classical biochemical trade-off for ammonium acquisition: bacterial AMO prioritizes capture capacity, making it ecologically suited for high-ammonium environments; whereas archaeal AMO prioritizes binding affinity, conferring a competitive advantage in oligotrophic habitats.

### Active center in archaeal AMO

To quantify the copper content in the *Nm*AMO sample, inductively coupled plasma (ICP) spectrometry was performed. Quantitative analysis revealed an average of approximately 1.1 copper per protomer (Fig. 3a and Extended Data Table 3), comparable to the copper content previously reported for particulate methane monooxygenase (pMMO) (*33*). This finding is corroborated by structural analysis, which identified two well-resolved copper-binding sites in *Nm*AMO, designated Cu_B_ and Cu_D_ (Fig. 3b). The Cu_B_ site is coordinated by three conserved histidine residues, as in related bacterial enzymes ^21–23^. However, the imidazole ring of His35 is rotated by >90° relative to its orientation in bacterial AMO, likely reflecting adaptation to a distinct local hydrophobic environment (Extended Data Fig. 8). Unlike bacterial AMO, where either a mononuclear Cu_C_ site or a dicopper Cu_C_-Cu_D_ center can be occupied ^21–23^, only the Cu_D_ site is occupied in active *Nm*AMO, whereas the Cu_C_ site remains vacant (Fig. 3b). Consistent with the recently identified active site in pMMO homologue ^33,34^, Cu_D_ in *Nm*AMO may constitute the functional catalytic center.

**Figure 3.**
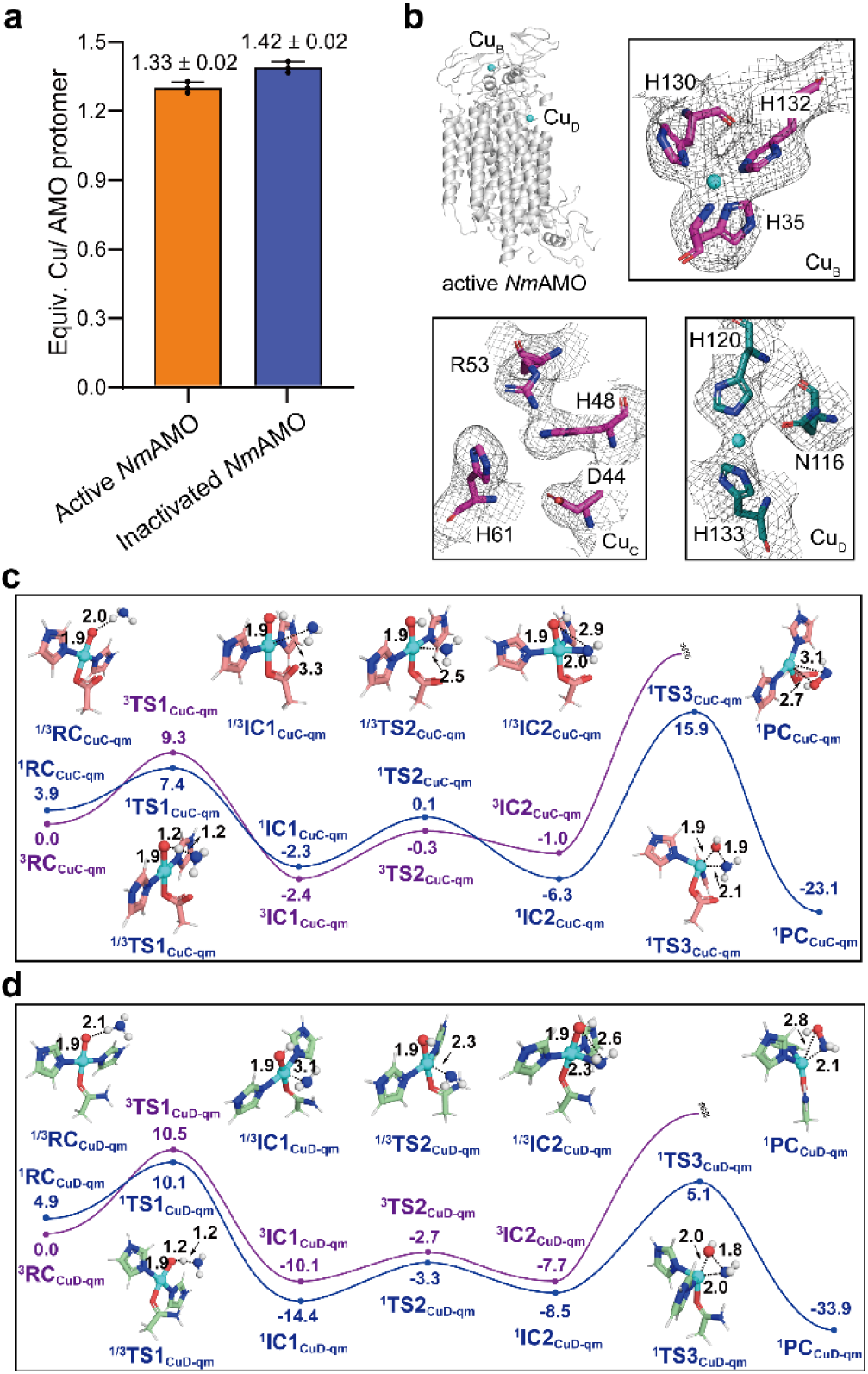
Cu_D_ is the potential active site of *Nm*AMO. **a.** Copper contents of active *Nm*AMO and inactivated *Nm*AMO in native membrane. Error bars represent standard deviation of n ≥ 3 biological replicates, each measured in triplicate. **b.** Copper sites in the active *Nm*AMO. A protomer of *Nm*AMO is depicted with the copper-binding sites indicated. Detailed coordination geometries of the copper ligands are displayed in the corresponding enlarged panels on the right. Omit maps (mesh) are contoured at 5σ. **c-d.** QM (UMN15/def2-TZVP//def2-SVP) calculated potential energy profile (in kcal/mol) for Cu_C_(II)−O^•–^ (**c**) and Cu_D_(II)−O^•–^ (**d**) mediated ammonia hydroxylation to hydroxylamine via the HAT, NH_2_^•^ rebound and NH_2_…OH coupling mechanism in both open-shell singlet and triplet states. Key distances are given in Å.

QM simulations of ammonia oxidation further support this Cu_D_-centered assignment. Although hydrogen abstraction from NH_3_ proceeds with comparable energy barriers from Cu_C_(II)–O^•^⁻ (7.4 kcal·mol⁻¹, from ^3^RC_CuC-qm_ to ^3^TS1_CuC-qm_) and Cu_D_(II)–O^•^⁻ species (10.1 kcal·mol⁻¹, from ^3^RC_CuD-qm_ to ^1^TS1_CuD-qm_), the subsequent hydroxyl–amino coupling step requires a substantially higher barrier via the Cu_C_-initiated pathway (22.2 kcal mol⁻¹, from ^1^IC2_CuC-qm_ to ^1^TS3_CuC-qm_ vs. 13.6 kcal mol⁻¹, from ^1^IC2_CuD-qm_ to ^1^TS3_CuD-qm_) (Fig. 3c-d and Extended Data Figs. 9 and 10). In addition, the overall reaction is more exothermic when initiated at the Cu_D_ site (ΔE = −33.9 kcal·mol⁻¹ vs. −23.1 kcal·mol⁻¹) (Fig. 3c-d). This mechanistic distinction explains the preferential utilization of Cu_D_ as the catalytic center in *Nm*AMO, consistent with the experimentally observed copper occupancy.

### Channel architecture and catalytic advantage of dicopper center

Building on the divergent ammonium-capture strategies observed at the extracellular surface, we next investigated whether internal substrate-access pathways and copper-center organization further differentiate archaeal and bacterial AMOs. MD simulations revealed a significantly narrower hydrophobic channel in *Nm*AMO than in *Nh*AMO, with minimum diameters of ∼5.4 Å and ∼7.1 Å, respectively (Fig. 4a-f and Extended Data Figs. 11 and 12). This constriction correlated with a longer substrate-dissociation time in *Nm*AMO (Fig. 4g). Furthermore, umbrella sampling calculations demonstrated substrate egress from *Nm*AMO encounters a substantially higher free-energy barrier (34.9 kcal mol⁻¹) than from *Nh*AMO (22.7 kcal mol^-1^), with the barrier maximum coinciding with the channel’s narrowest point (Extended Data Fig. 13). Collectively, these data indicate that archaeal AMO enforces a more restrictive internal pathway, whereas bacterial AMO facilitates a more accessible route for substrate and product exchange, consistent with the higher catalytic throughput observed in AOB under ammonium-replete conditions.

**Figure 4.**
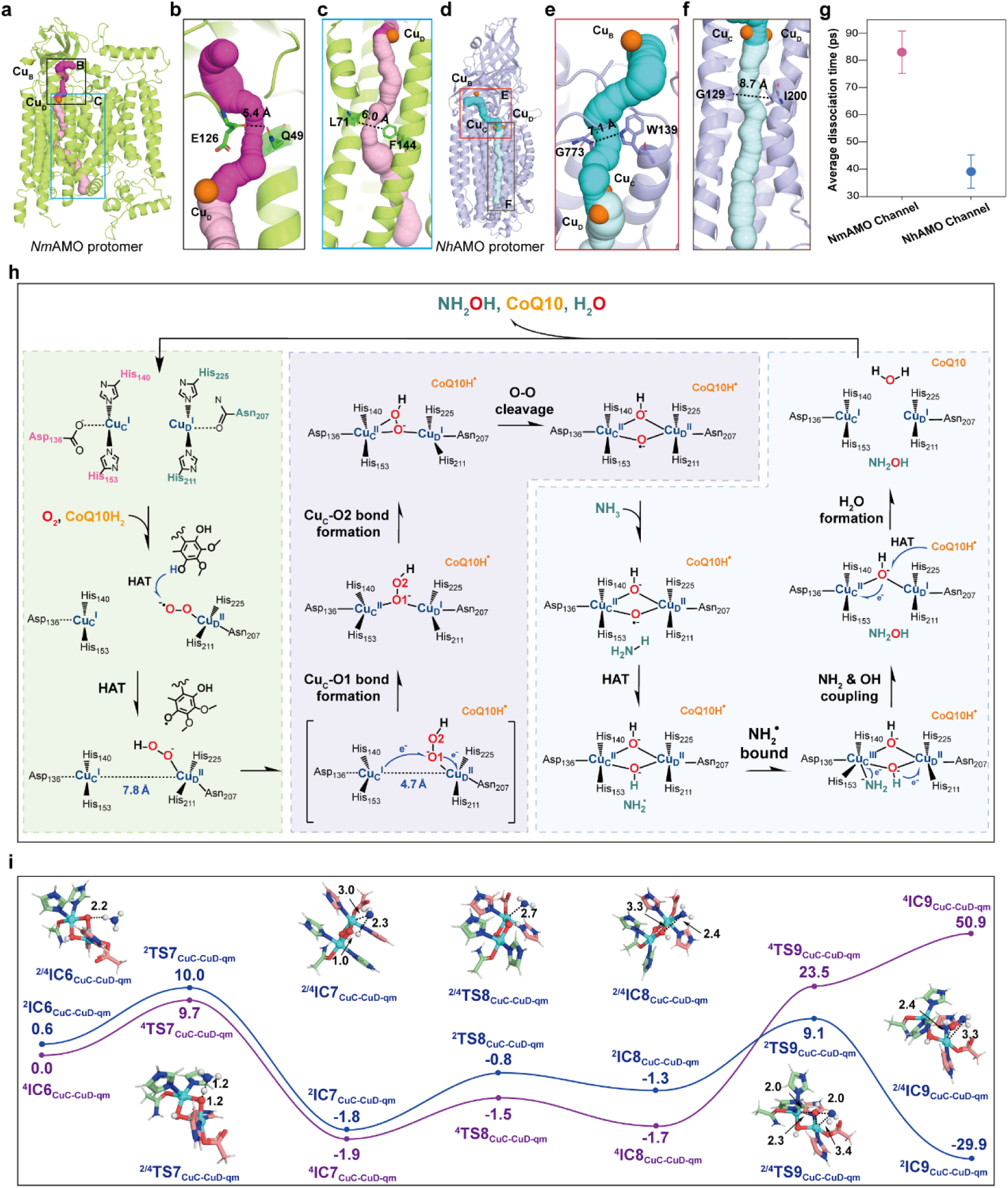
Channel architecture and reaction mechanism of the dinuclear copper center. **a-f.** Overview diagram illustrating the most favorable entry pathways in *Nm*AMO (**a**) and *Nh*AMO (**d**). The upstream entry pathways of the Cu_D_ site in *Nm*AMO (**b**) and Cu_C_-Cu_D_ site in *Nh*AMO (**e**) are highlighted in pink and cyan respectively, whereas the downstream routes from the Cu_D_ site in *Nm*AMO (**c**) and Cu_C_-Cu_D_ site in *Nh*AMO (**f**) are highlighted in light pink and cyan respectively. The amino acid residues at the narrowest regions of the *Nm*AMO and *Nh*AMO channels are shown as green and purple stick models, respectively. **g.** Times required for substrate to traverse from entry into the Cu_D_ centers of *Nm*AMO and the Cu_C_-Cu_D_ center of *Nh*AMO to dissociation into the external environment. **h.** Overview of the proposed catalytic cycle of the co-occupied Cu_C_-Cu_D_ center. The reaction is initiated by O_2_ activation at the dinuclear copper center (green background), followed by the bimetallic Cu-based reactive oxygen species (*μ*-oxo)(*μ*-hydro)Cu_C_(II)Cu_D_(II) generation (purple background), substrate oxidation steps, ultimately leading to product formation (blue background). **i.** QM (UMN15/def2-TZVP//def2-SVP) calculated potential energy profile (in kcal/mol) for (*μ*-oxo)(*μ*-hydro)Cu_C_(II)Cu_D_(II) mediated ammonia hydroxylation to hydroxylamine via the HAT, NH_2_^•^ rebound and NH_2_…OH coupling mechanism in both doublet and quartet states. Key distances are given in Å.

While the channel architecture dictates substrate traffic, the copper-center configuration governs the chemistry of ammonia oxidation. In contrast to archaeal AMO, which harbors a mononuclear Cu_D_ site, recent structures of bacterial AMO reveal that co-occupied Cu_C_ and Cu_D_ sites can assemble into a dinuclear copper center (Extended Data Fig. 14). Given that endogenous CoQ10H_2_ has been proposed as a physiological reductant ^21,23^, we assessed its interaction with this site. MD simulations indicated that CoQ10H_2_ access the substrate-accessible pocket of membrane-bound *Nh*AMO, positioning a hydroxyl group proximal to the Cu_D_-bound superoxo species (Extended Data Figs. 15-19). QM/MM calculations further showed that hydrogen atom transfer (HAT) from CoQ10H_2_ to this superoxo intermediate proceeds with a modest energy barrier of 14.7 kcal mol⁻¹ (from ^3^IC1_CuD_ to ^3^TS2_CuD_), yielding a Cu_D_(II)–OOH⁻ species (Extended Data Figs. 20 and 21). This barrier is comparable to that reported for pMMO ^35^, underscoring the intrinsic reactivity of the Cu_D_-bound superoxo species.

Following proton-coupled reduction, the hydroperoxo intermediate bridges to the adjacent Cu_C_ site, forming a hydroperoxo dicopper complex with a low energetic barrier (5.6 kcal mol⁻¹) (Extended Data Figs. 22 and 23). Direct coupling of a Cu_D_-bound superoxo to Cu_C_ was found to be energetically unfavorable (Extended Data Figs. 24 and 25), indicating that binuclear oxygen species formation is energetically favored via the Cu(II)–OOH⁻ intermediate ^36^. Subsequent O–O bond cleavage (11.1 kcal mol⁻¹) generates a catalytically competent *μ*-oxo-*μ*-hydroxo-Cu_C_(II)Cu_D_(II) core (Extended Data Fig. 26). Notably, this step is strongly favored when the hydroxyl group migrates to Cu_C_, revealing an asymmetric division of labor: Cu_D_ specializes in initial oxygen activation and reduction, while Cu_C_ accommodates the second oxygen unit during O– O bond cleavage (Fig. 4h and Extended Data Fig. 27).

To evaluate the catalytic consequence of this binuclear center, we computed the energy landscape for ammonia hydroxylation starting from the *μ*-oxo-*μ*-hydroxo dicopper configuration (IC6_CuC–CuD_, Fig. 4i and Extended Data Fig. 28). This pathway proceeds with significantly lower energy barriers than those calculated for mononuclear Cu_C_ or Cu_D_ centers. HAT from NH_3_ requires only 9.7 kcal·mol^-1^, followed by a nearly barrierless NH_2_^•^ rebound and hydroxyl-amino coupling (11.0 kcal mol⁻¹) to produce NH_2_OH (IC9_CuC–CuD_) (Fig. 4i and Extended Data Figs. 10 and 29). These calculations establish that the bacterial dicopper center provides a kinetically favorable, low-barrier route for both oxygen activation and ammonia hydroxylation, mechanistically explaining the higher ammonia oxidation rates (AORs) observed in bacteria harboring a co-occupied Cu_C_-Cu_D_ center (Fig. 2g).

In summary, the integration of channel architecture and active-site analyses reveals two synergistic axes of archaeal–bacterial AMO divergence. Bacterial AMO pairs a capacious hydrophobic channel with a highly efficient dicopper catalyst, a combination that supports rapid ammonia oxidation in high-ammonium environments. Conversely, archaeal AMO employs a narrower internal pathway and a mononuclear Cu_D_ site, architectural choices that prioritize substrate retention over catalytic speed—aligning with its ecological dominance in oligotrophic habitats. Together with the extracellular capture strategies described above, these findings provide a comprehensive molecular framework linking AMO structural specialization to the distinct physiological behaviors of AOA and AOB.

### Inactivated structure of archaeal AMO

Although NIs have been widely employed to suppress ammonia oxidation activity of ammonia-oxidizing microorganisms, their efficacy exhibits considerable variation across different lineages ^27^. Inhibitors strongly inhibit AOB but are far less effective against AOA, with significant suppression only at high concentrations (Fig. 5a and Extended Data Fig. 30). To elucidate the structural basis of AOA inhibition, we determined the cryo-EM structure of the membrane-bound *Nm*AMO complex isolated from SCM1 cells treated with a high concentration of ATU, at a resolution of 2.75 Å (Extended Data Fig. 31 and Extended Data Table 1). The overall architecture of the inactivated *Nm*AMO closely resembles the active enzyme (Fig. 5b). Crucially, no electron density corresponding to ATU was observed in the active site, suggesting that inhibition of archaeal AMO activity does not involve direct binding of NIs. Instead, the copper center undergoes a major rearrangement: the copper density was absent from the Cu_D_ site and uniquely coordinated at the Cu_C_ site (Fig. 5c). Supporting this structural observation, ICP spectrometry indicated identical copper content in both inactivated and active *Nm*AMO (Fig. 3a), indicating the presence of two copper ions per protomer. Collectively, these data suggest that the tested inhibitors indirectly inactivate *Nm*AMO by triggering copper relocation.

**Figure 5.**
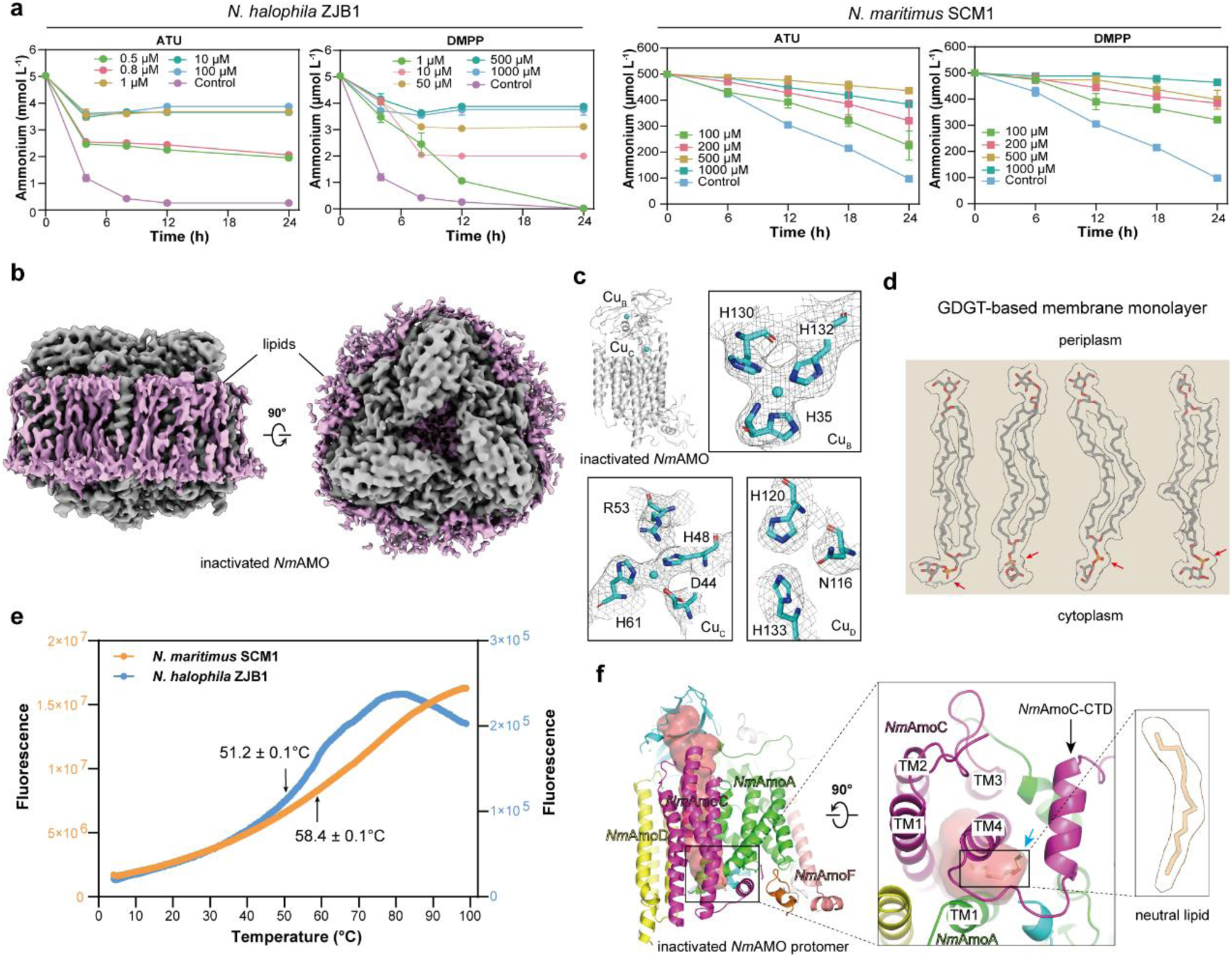
Inactivation of archaeal AMO. **a.** Average inhibition of ammonia oxidation in AOB (*N. halophila* ZJB1, left two panels) and AOA (*N. maritimus* SCM1, right two panels) under varying concentrations of ATU and DMPP. Error bars represent standard deviations from ≥3 biological replicates, each measured in triplicate. **b.** 3D structure of the inactivated *Nm*AMO complex. **c.** Copper sites in the inactivated *Nm*AMO. A protomer of *Nm*AMO is depicted with the copper-binding sites indicated. The detailed coordination geometries of the copper ligands are displayed in the corresponding enlarged panels on the right. Omit maps (mesh) are contoured at 5σ. **d.** Representative archaeal GDGT lipids modeled in the *Nm*AMO structure. The cytoplasmic negatively charged phosphate groups are indicated by red arrows. Electron densities are shown as transparent surfaces. **e.** Thermal denaturation profiles of purified *Nm*AMO and *Nh*AMO monitored by fluorescence. The corresponding thermal melting temperatures (TM) are indicated by black arrows. **f.** Surface representation of the hydrophobic channel within the *Nm*AMO protomer, showing obstruction by a neutral lipid. The lipid’s position is boxed and magnified in the right panel, with a blue arrow indicating its location. Corresponding AMO subunits are color-coded.

Additionally, the inactivated *Nm*AMO exhibits improved definition of surrounding membrane lipids compared to the wild-type enzyme (Fig. 5b), likely reflecting lower conformational flexibility upon inactivation. Guided by comprehensive lipidomic analysis, we modeled numerous archaeal-specific glycerol dialkyl glycerol tetraether (GDGT) lipids bound to the *Nm*AMO surface (Fig. 5d and Extended Data Fig. 32). These lipids form a characteristic monolayer that spans the entire membrane (Fig. 5b and 5d), a feature proposed to enhance membrane stability ^37–39^. Structural analysis shows extensive interactions between AMO and the polar headgroups of these lipids. These interactions are predominantly mediated by positively charged residues from *Nm*AMO (Extended Data Fig. 33a). Thermal denaturation assays revealed that purified *Nm*AMO exhibits a higher thermal melting temperature (Tm) than *Nh*AMO (Fig. 5e), suggesting that GDGT-mediated interactions contribute to the enhanced structural stability of *Nm*AMO. Moreover, the cryo-EM density map showed pronounced enrichment of phosphate-containing GDGT headgroups at the cytoplasmic interface of *Nm*AMO (Fig. 5d and Extended Data Fig. 33b-c), suggesting that archaeal GDGTs confer asymmetric stabilization to the cytoplasmic and periplasmic domains of AMO.

We also observed an unassigned lipid-like density inserted into the lateral opening of the hydrophobic channel formed by a five-TM-helix bundle (Fig. 5f)—a feature absent in bacterial AMO ^21–23^. Given its location, the lipid-like density may influence the local conformation or product-egress dynamics of the hydrophobic channel, analogous to pore lipids in TRAAK channels ^40^. This hypothesis was supported by our active-state *Nm*AMO structure, in which the corresponding density exhibits increased flexibility, suggesting possible mobility of this lipid-associated region. This finding redefines the architecture of AMO: the hydrophobic channel does not constitute a continuous conduit; instead, its cytoplasmic opening faces a hydrophobic interface formed by TM4 and the C-terminal domain of AmoC (Fig. 5f), delineating a product-exit route distinct from that of bacterial pMMO and AMO (Extended Data Fig. 34). Further functional studies may help establish the physiological role of this lipid-like density.

### Inhibition of bacterial AMO

The pronounced difference in sensitivity to NIs between archaeal and bacterial AMOs suggests distinct mechanisms of action. In AOB, inhibition followed apparent biphasic kinetics, with continued ammonia consumption during the initial phase followed by a rapid plateau, suggesting delayed inhibitor access before efficient active-site inhibition. By contrast, the more gradual response of AOA is consistent with weaker or indirect AMO inactivation (Fig. 5a and Extended Data Fig. 30). Given the presence of a dinuclear Cu_C_-Cu_D_ copper center in *Nh*AMO and the potent inhibition of AOB by NIs, we hypothesized that these inhibitors act by directly targeting this dicopper site ^23^. To test this, we examined ATU and DMP, the dephosphorylated active form of DMPP, and determined cryo-EM structures of *Nh*AMO bound to DMP at 2.15 Å and ATU at 2.47 Å (Extended Data Figs. S35-36). These high-resolution reconstructions enabled precise atomic modeling at the side-chain level to accurately analyze the coordination environments of these small molecule inhibitors (Fig. 6a-b and Extended Data Figs. 35-36). Overall, the two inhibitor-bound structures are highly similar, with the periplasmic subunit *Nh*AmoE showing reduced or absent density—particularly in the DMP-bound complex (Fig. 6a-b)—suggesting destabilization or dissociation of this putative regulatory subunit ^23^.

**Figure 6.**
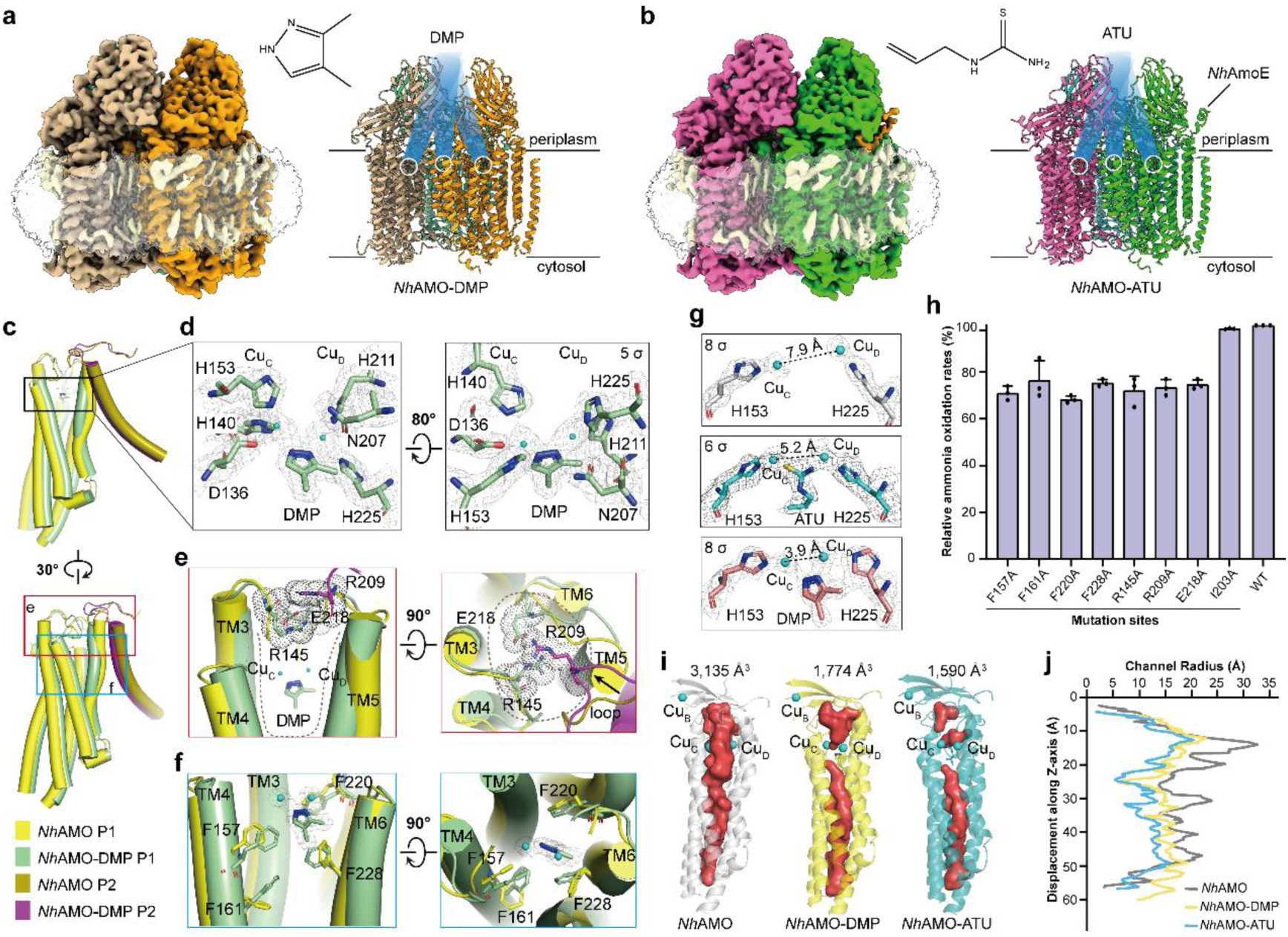
Structures of inhibited bacterial AMO. **a-b.** Overall structures of *Nh*AMO in complex with DMP (**a**) and ATU (**b**). Cryo-EM maps (left) and corresponding atomic models (right) are shown. The positions of inhibitor molecules within each AMO protomer are indicated by white circles, with their respective chemical structures depicted. **c.** Structural superposition of wild-type *Nh*AMO and *Nh*AMO-DMP revealing conformational changes in the hydrophobic channel. Models from adjacent AMO protomers are denoted as P1 and P2 and color-coded accordingly. **d.** DMP occludes the active site by simultaneously chelating the Cu_C_ and Cu_D_ copper ions. **e.** A sandwich-like lid structure, formed by R209 from the neighboring loop and E218 and R145 from the *Nh*AmoC subunit, closes the entrance of the hydrophobic channel upon inhibitor binding through conformational rearrangement. The three key residues are represented as meshed surfaces. **f.** A conserved cluster of hydrophobic residues (F157, F161, F220, F228) on the inner surface of the hydrophobic channel undergoes inward contraction upon inhibitor binding. **g.** Distances between Cu_C_ and Cu_D_ metal centers in *Nh*AMO (top, 7.9 Å), *Nh*AMO-ATU (middle, 4.7 Å), and *Nh*AMO-DMP (bottom, 3.9 Å). **h.** Ammonia oxidation activity of wild-type (WT) and single-copy mutant *N. halophila* ZJB1 strains. Mutations target residues coordinating the hydrophobic F-cluster or the inhibitor-induced lid region. I203A of AmoA serves as a structural control: this residue resides in a loop adjacent to, but outside, the proposed lid region. Data are represented as the mean ± SD of n = 3 independent biological replicates. \*\**p* < 0.01, \*\*\*\**p* < 0.0001 (one-way ANOVA with Tukey’s test). **i.** Volume renderings of the hydrophobic channels in *Nh*AMO, *Nh*AMO-DMP and *Nh*AMO-ATU. The α-helices forming the hydrophobic channel are displayed as cartoons, while channel volumes are shown as surfaces with values labeled. The central pores (generated using HOLLOW software ^44^) are shown as red surfaces. **j.** Channel radius of the pore channel in *Nh*AMO (gray), *Nh*AMO-DMP (yellow) and *Nh*AMO-ATU (blue). Contour levels of the omit maps (mesh) are indicated for panels (**d)** and (**g**).

In the *Nh*AMO-DMP structure, the inhibitor bridges the dicopper center via its pyrazole nitrogens, while a methyl group forms a hydrogen bond with His225 (Fig. 6c-d). This binding pulls the two coppers 4.0 Å closer together (Fig. 6g), compacts the active site, and induces a major conformational change in a loop of the adjacent *Nh*AmoC located above the hydrophobic channel. Within this loop, R209 projects into the channel center, triggering inward displacement of E218 and rotation of R145 to form a sandwich-like lid that occludes the channel entrance (Fig. 6e). Mutating these residues reduced ammonia oxidation activity by ∼30% (Fig. 6h), which is comparable to that of the deletion of single AMO subunit ^41^, underscoring the functional importance of this lid region. As this loop mediates inter-protomer interactions, lid formation likely couples inhibitor binding to coordinated regulation across the homotrimer. Indeed, structural comparisons show that inhibitor binding induces the global constriction of *Nh*AMO (Extended Data Fig. 37a). In addition, the inhibitor is further stabilized by a conserved aromatic cluster (F157, F161, F220, and F228), with π–π stacking between F220 and F228 effectively sandwiching the DMP molecule (Fig. 6f). Mutation of these residues also significantly impairs ammonia oxidation activity (Fig. 6h). Sequence alignments show that these key residues are relatively conserved across AOA, AOB, and methanotrophic bacteria (Extended Data Fig. 38), suggesting an evolutionarily conserved role in ammonia and methane oxidation.

Concomitant with inhibitor binding, the volume of the hydrophobic channel is substantially reduced—from 3,135 Å^3^ in the active enzyme to 1,774 Å^3^ in the DMP-bound state. Moreover, inhibitor occupancy blocks the hydrophobic channel and divides it into two isolated segments, thereby effectively preventing substrate passage (Fig. 6i and 6j). Another representative NI, ATU, exerts a similar inhibitory effect, though with a distinct coordination geometry that shortens the Cu_C_-Cu_D_ distance to 4.7 Å (Fig. 6g, 6i, 6j, and Extended Data Fig. 37).

Together, these results demonstrate that ATU and DMP inhibit bacterial AMO by directly occupying the dinuclear copper center and inducing channel constriction near the active-site region.

### Two distinct pathways of AMO inactivation

By integrating cryo-EM structures of archaeal AMO in active and inhibitor-treated states with those of bacterial AMO in complex with NIs, together with complementary biochemical, kinetic, omics, and computational data, our work reveals two representative and distinct inactivation pathways for AMO (Fig. 7).

**Figure 7.**
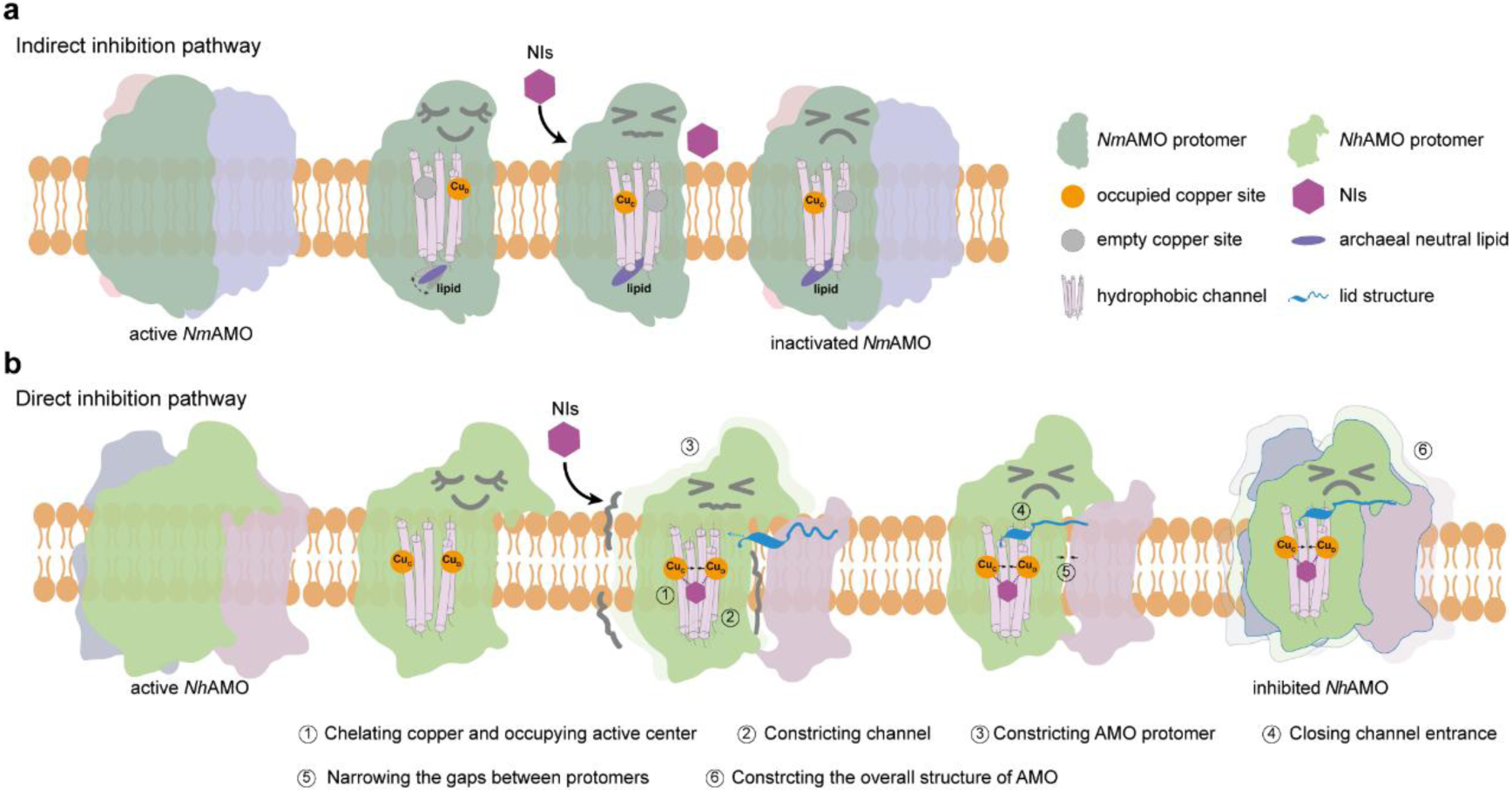
Schematic illustration of the indirect and direct inhibition pathways of NIs on archaeal and bacterial AMO. **a.** Indirect inhibition pathway: archaeal AMO activity may be inhibited by NIs through interference with other key processes involved in ammonia oxidation, rather than by direct binding to the AMO active center. This indirect inhibition results in the repositioning of copper ligands from the active Cu_D_ site to Cu_C_ site, accompanied by obstruction of the hydrophobic channel opening by neutral lipid-like density. **b.** Direct inhibition pathway: bacterial AMO activity is inhibited by direct interaction of NIs with the dicopper ligands, resulting in occupation of the active center, inhibitor-induced constriction of the hydrophobic channel, compaction of the AMO protomer, and overall structural constriction of the AMO complex.

For archaeal AMO, the inhibitors examined here do not act through specific, high-affinity binding. Substantial inhibition requires high concentrations and correlates with copper relocation from the Cu_D_ active site to the Cu_C_ site. This copper displacement, coupled with altered ordering of neutral lipid-like density near the hydrophobic channel, suggests an indirect inhibition mechanism in which NIs perturb the catalytic center without directly binding the enzyme. In contrast, bacterial AMO is potently and specifically inhibited by direct NI binding. Inhibitors coordinate the dinuclear Cu_C_–Cu_D_ center and induce a conformational change that forms a lid structure over the hydrophobic channel. This structural remodeling substantially constricts the hydrophobic channel and effectively abolishes ammonia oxidation activity.

Thus, our findings present two functionally divergent AMO inhibition strategies of ammonia-oxidizing microorganisms: an indirect, copper-displacing mechanism in archaea versus a direct, active site-occluding mechanism in bacteria (Fig. 7), providing a structural foundation for understanding the differential efficacy of NIs and the functional evolution of AMO across domains of life.

## Discussion

AMO catalyzes the initial step of nitrification and has long served as a central enzymatic nexus for understanding how ammonia-oxidizing microorganisms derive energy and regulate nitrogen biogeochemical fluxes. Nevertheless, mechanistic insight into AMO has remained limited. Our study reveals that archaeal and bacterial AMOs have undergone coordinated structural and functional divergence in enzyme architecture, substrate recruitment, transport channel, copper-site organization, membrane-lipid association, and inhibitor response. These findings collectively establish a molecular framework for rationalizing the distinct physiological adaptations and niche-specific chemistries of AOA and AOB.

A major implication of the *Nm*AMO structure is that the evolutionarily conserved AMO catalytic core is embedded within an archaeal-specific scaffold. Within this scaffold, the AmoA, AmoB, and AmoC subunits form a shared catalytic framework, while lineage-specific accessory subunits mediate interprotomer stabilization, forming an unexpected cup-shaped trimeric assembly. Comparative structural analyses suggest that the core AMO architecture and copper-coordination motifs are broadly conserved, whereas accessory subunits are more variable (Extended Data Figs. 39-42) and may fine-tune enzyme stability, membrane organization, or regulatory responses in a lineage-dependent manner. Future genetic and biochemical dissection of individual accessory subunits will be important for defining how this architectural elaboration controls AMO activity across archaeal lineages.

Our data further provide an enzyme-level explanation for the long-observed physiological divergence between AOA and AOB. AOA generally prevail under low-ammonium conditions, whereas AOB often dominate in ammonium-replete environments. The present structures and functional analyses suggest that this ecological pattern reflects a trade-off between substrate affinity, capture capacity, and catalytic throughput. Bacterial AMO presents an expanded extracellular cupredoxin surface, a more accessible hydrophobic pathway, and, in some species or states, a catalytically favorable dicopper center. These features are consistent with high-capacity ammonium capture, rapid substrate/product exchange, and high catalytic output. In contrast, archaeal AMO lacks the second extracellular cupredoxin domain but displays stronger local electronegativity, a narrower internal pathway, and a mononuclear Cu_D_ center with comparatively lower catalytic efficiency. These features synergistically favor high-affinity acquisition and longer substrate residence at the cost of lower turnover. We therefore propose that AMO constitutes an important enzymatic layer contributing to niche differentiation between AOA and AOB: bacterial AMO is functionally optimized for catalytic throughput under ammonium-replete conditions, whereas archaeal AMO is evolutionarily tuned for high-affinity substrate acquisition and retention in ammonium-limited environments.

This model also helps explain the relationship between NH_4_^+^ enrichment and NH_3_ oxidation. NH_3_ is generally considered the substrate oxidized by AMO, whereas NH_4_^+^ predominates under near-neutral environmental conditions. Our results suggest that extracellular and periplasmic architectures enrich ammonium near the membrane enzyme and thereby buffer local NH_3_ availability. In AOA, this process is likely integrated with broader cell-envelope features such as the recently reported archaeal S-layer ^17,42,43^. How NH_4_^+^ enrichment is coupled to NH_3_ delivery remains unresolved. Conserved proton-accepting residues along the hydrophobic pathway may participate in proton transfer or local deprotonation (Extended Data Fig. 43), but direct experimental evidence is still required. Definitive insight will require integrated approaches combining pH-dependent kinetics, isotope tracing, mutagenesis, and time-resolved structural approaches to pinpoint the spatial and temporal dynamics of the NH_4_^+^/NH_3_ equilibrium shift during AMO catalysis.

Together with recent structural insights into bacterial AMOs ^21–23^, copper-site organization suggests that AMO enzymes may operate through a spectrum of catalytic configurations rather than a single invariant active-site state. In *Nm*AMO, copper occupancy is centered at Cu_D_, whereas bacterial AMOs can use a mononuclear Cu_C_ site or a co-occupied Cu_C_–Cu_D_ center (Extended Data Fig. 44). Our calculations indicate that the dicopper configuration provides a lower-barrier route for oxygen activation and ammonia hydroxylation than mononuclear pathways. These observations support a testable hierarchy in which AMOs containing an expanded cupredoxin surface and a dicopper center are expected to exhibit the greatest catalytic capacity, followed by enzymes with expanded cupredoxin surfaces but mononuclear centers, and finally enzymes with a single cupredoxin domain and mononuclear copper usage. This hierarchy should be viewed as a conceptual framework rather than a fixed rule, because copper occupancy may depend on species, redox state, substrate availability, metal homeostasis, and membrane environment. Capturing additional archaeal and bacterial AMO states will be necessary to determine how dynamic copper-site switching contributes to AMO evolution and activity.

The intimate association between *Nm*AMO and archaeal tetraether lipids introduces an additional layer to AMO specialization, notably enhancing thermostability. The asymmetric distribution of GDGTs and the presence of a lipid-like density near the hydrophobic pathway raise the possibility that archaeal membrane composition may influence AMO conformational dynamics, product egress, or inhibitor accessibility. At present, these roles remain inferential. Reconstituting *Nm*AMO in defined lipid environments and testing lipid perturbations in vivo or in native-like membrane systems will be essential to determine whether GDGTs directly regulate catalysis or primarily stabilize the enzyme scaffold.

The inhibitor-bound and inhibitor-treated structures reveal a second major axis of archaeal–bacterial divergence. In bacterial AMO, DMP and ATU directly coordinate the Cu_C_–Cu_D_ center, induce active-site compaction, and trigger closure of the connected hydrophobic channel. These structural changes provide a mechanistic basis for the pronounced sensitivity of AOB to copper-chelating inhibitors. They also account for the biphasic inhibition kinetics observed in bacterial whole-cell assays: the initial phase likely reflects inhibitor partitioning, membrane access, and target engagement, whereas the subsequent plateau reflects a rapid conversion of the active enzyme population into a channel-occluded inhibited state. In contrast, ATU-treated *Nm*AMO lacks detectable inhibitor density at the active site and instead exhibits structural signatures consistent with an indirect, copper-displacing inactivation mechanism. The narrower internal substrate pathway in *Nm*AMO, coupled with extensive association with GDGTs, likely impedes efficient inhibitor access to the catalytic pocket, thereby contributing to the attenuated and kinetically slower inhibition profile characteristic of AOA.

These findings also hold significant practical implications for the development of AMO-targeting inhibitors, as they identify key structural and functional features that can inform next-generation inhibitor design. Specifically, such features include (i) enhanced accessibility to the membrane-embedded AMO channel, (ii) selective engagement with lineage-specific copper centers, and (iii) avoidance of nonspecific metal chelation. Such principles may enable more selective modulation of nitrification in defined ecological settings, including marine, estuarine, and agricultural systems where AOA and AOB contribute differently to nitrogen turnover and N_2_O production. Several limitations should caution interpretation of this work. The experimentally resolved archaeal and bacterial AMO structures from soil-associated lineages will be required to define the full environmental diversity of AMO. AMO architecture constitutes a major enzyme-level determinant of niche differentiation. Integrating these AMO-encoded properties with ammonium transport, respiratory coupling, copper homeostasis, membrane bioenergetics, nitrogen-source regulation, and community-level interactions will ultimately reveal how ammonia oxidizers establish distinct ecological strategies in natural systems. Finally, structures of additional catalytic intermediates, copper-occupancy states, and inhibitor-bound archaeal AMO conformations will be necessary to establish the complete catalytic and inhibitory cycle.

## Acknowledgements

We sincerely thank the staff at the cryo-EM center of Southern University of Science and Technology for their technical support on the Cryo-EM and High-Performance Computation platforms. Z.L. is an investigator of SUSTech Institute for Biological Electron Microscopy. We thank Fengfeng Zheng for valuable discussions regarding archaeal lipids.

## Author Contributions

Z.L. and X.Y. conceived this project and designed the experiments. Z.L., C.Z., W.P., H.D. and G.Z. supervised the project. T.M. and Z.L. cultured the *N. halophila* ZJB1 and *N. maritimus* SCM1. T.M., Z.L. and X.Y. purified the AMO samples and prepared the samples for EM. X.Y. and Z.L. collected the EM data, performed the EM analysis, the model building and the structural analysis. Z.H., W.P. and S.L. did the MD simulations, QM calculations and data analysis. T.M. constructed the mutants and did the enzymatic activity assays. Y.C., C.Z. and G.Z. performed the archaeal lipidomics analysis. T.M. and P.J. conducted the enzyme kinetics assays. X.Y., Z.L., T.M. and W.P. wrote the initial draft; Z.L., X.Y., C.Z. and W.P. edited the manuscript.

## Declaration of generative AI and AI-assisted technologies in the writing process

During the preparation of this manuscript, the author(s) used ChatGPT in order to improve language. After using this tool or service, the authors reviewed and edited the content as needed and take full responsibility for the content of the publication.

## Data and materials availability

The cryo-EM maps of purified *Nh*AMO in complex with DMP (EMD-66948) and ATU (EMD-66954), as well as active (EMD-66926) and inactivated (EMD-66857) membrane-bound *Nm*AMO, have been deposited in the Electron Microscopy Data Bank (http://www.ebi.ac.uk/pdbe/emdb/). The corresponding atomic coordinates (PDB: 9XK5, 9XKB, 9XJ2 and 9XGS) have been deposited in the Protein Data Bank (http://www.rcsb.org). All other data generated or analyzed during this study are included in this article and its supplementary information files.

## Supplementary Information

The PDF file includes:

Materials and Metphods

Extended Data Figures 1 to 44

Extended Data Tables 1 to 8

References 45-129

## Materials and Methods

### Cultivation of ammonia-oxidizing microorganisms

*Nitrosopumilus maritimus* SCM1, *Nitrosomonas halophila* ZJB1 (recently isolated in our laboratory), and *Nitrosomonas europaea* were cultured as previously described ^6,45^. Briefly, pure cultures of *N. maritimus* SCM1 were maintained in HEPES-buffered synthetic Crenarchaeota medium (per liter: 26 g NaCl, 5 g MgCl_2_·6H_2_O, 5 g MgSO_4_·7H_2_O, 1.5 g CaCl_2_, 0.1 g KBr, 10 mL HEPES buffer [comprising 1 mol L⁻¹ HEPES and 0.6 mol L⁻¹ NaOH], 5 mL KH_2_PO_4_ solution [0.4 g L⁻¹], 2 mL bicarbonate solution [1 mol L⁻¹], 1 mL FeNaEDTA solution [7.5 mol L⁻¹], and a mixture consisting of 1 mL vitamin solution, 1 mL non-chelated trace element solution (TES), 1 mL streptomycin [100 mg mL⁻¹], and 1 mL amphotericin B [1 mg mL⁻¹]) ^6,46,47^. *N. halophila* ZJB1 was maintained in a modified synthetic seawater medium (per liter: 5 g NaCl, 0.25 g KCl, 0.04 g MgSO_4_·7H_2_O, 0.1 g CaCl_2_·2H_2_O, 10 mL HEPES buffer, 5 mL KH_2_PO_4_, 2 mL bicarbonate solution, 1 mL TES solution and 1 mL phenol red [0.05%] as pH indicator) ^48^. *N. europaea* was grown in a modified synthetic freshwater medium (per liter: 0.5 g NaCl, 0.04 g MgSO_4_·7H_2_O, 0.04 g CaCl_2_·2H_2_O, 10 mL HEPES buffer, 5 mL KH_2_PO_4_, 2 mL bicarbonate solution, 1 mL TES solution and 1 mL phenol red [0.05%] as pH indicator) ^45,48^. The pH of all media was adjusted to 7.5–7.8. Stock cultures were grown in 2 L bottles containing 1.2 L of medium, supplemented with 2 mmol L⁻¹, 10 mmol L⁻¹ and 10 mmol L⁻¹ NH_4_⁺ for *N. maritimus* SCM1, *N. halophila* ZJB1 and *N. europaea*, respectively, as the nitrogen source. *N. maritimus* SCM1 was incubated without agitation, whereas *N. halophila* ZJB1 and *N. europaea* were grown with shaking at 120 rpm. All strains were incubated at 30°C in the dark to prevent light-induced inhibition. Growth was monitored by measuring ammonium consumption, and culture purity was assessed on Luria–Bertani (LB) broth according to a previously described method ^49^.

### Inhibitor activity assays

To evaluate the efficacy of NIs against ammonia-oxidizing microorganisms, we investigated the activity responses to five commonly used NIs: allylthiourea (ATU), 3,4-dimethylpyrazole phosphate (DMPP), dicyandiamide (DCD), amidinothiourea (ASU), and nitrapyrin (NP) (Extended Data Fig. 30). In each experiment, cells of ammonia-oxidizing microorganisms from batch cultures in the exponential growth phase were aliquoted into sterile 100 mL glass bottles (20 mL per bottle). A range of inhibitor concentrations was then added to the culture medium (Extended Data Fig. 30). Cultures containing no NIs but inoculated identically served as controls. Samples were collected at regular intervals (6 h for AOA and 4 h for AOB) to monitor ammonium (NH_4_⁺) consumption, thereby tracking ammonia oxidation activity in ammonia-oxidizing microorganisms.

### Chemical analyses

Ammonium concentrations were determined using the optimized indophenol blue method ^50^. In brief, 200 μL of culture was transferred into sterile 1.5 mL microcentrifuge tubes and centrifuged at 15,000 × g for 10 min at room temperature to obtain pellets of cellular debris and large particulates. The supernatant was then transferred to new sterile 1.5 mL Eppendorf tubes and diluted to a final volume of 1 mL with NH_4_^+^-free ultrapure water. For the optimized indophenol blue assay, 50 μL of phenol-nitroprusside solution and 50 μL of alkaline hypochlorite solution were added. After complete color development, absorbance at 630 nm was measured using a microplate reader (EnSpire) for ammonium quantification. Final concentrations were calculated using standard curves for NH_4_^+^ (0-10 μM, *n* = 3, *R*^2^ > 0.998).

### Archaeal membrane fraction extraction and bacterial AMO protein purification

For archaeal membrane fraction extraction, *N. maritimus* SCM1 cells were cultured in HEPES-buffered synthetic Crenarchaeota medium without inhibitor (for active AMO) or with 1000 μM ATU (for inactivated AMO). A total of 70 L of *N. maritimus* SCM1 culture was harvested by vacuum filtration through a 0.22 µm pore polycarbonate membrane filter (Millipore) and resuspended in 50 mL lysis buffer (50 mM PIPES pH 7.2, 50 mM NaCl, 30 μM CuSO_4_). For inactivated AMO preparation, the lysis buffer was supplemented with 10 μM ATU throughout the preparation. Cell disruption was carried out at 60 W power (2 s on and 2 s off) for 7 min using an ultrasonic disruptor (Xinzhi Instrument, Zhejiang, China). The resulting lysate was centrifuged at 15,000 rpm at 4 °C for 10 min to remove cellular debris and intact cells. The supernatant was collected and ultracentrifuged at 150,000 × g at 4 °C for 1 h. The final membrane pellet was resuspended in 4 mL of lysis buffer, flash-frozen in liquid nitrogen and stored at −80°C for subsequent use.

For purification of bacterial AMO-NI complexes, *N. halophila* ZJB1 cells were maintained in a modified synthetic seawater medium with 10 μM ATU or 100 μM DMPP. Because DMPP can be dephosphorylated under the experimental conditions, the bound density was modeled as DMP, the dephosphorylated active form of DMPP. A total of 120 L of *N. halophila* ZJB1 cells expressing Flag-tagged AmoA were harvested via vacuum filtration using a 0.22 µm pore polycarbonate membrane filter (Millipore). The collected cells were then resuspended in lysis buffer (50 mM PIPES, pH 7.2; 50 mM NaCl; 30 µM CuSO_4_; and either 1 μM ATU or 10 μM DMPP) and lysed by sonication on ice. Subsequently, n-Decyl-β-D-maltopyranoside (DDM) was added to achieve a final concentration of 2%, followed by stirring for 2 h at 4 °C to extract membrane proteins. Subsequently, cell debris was removed by centrifugation at 20,000 × g for 30 min at 4 °C. The resulting supernatant was collected and incubated with anti-FLAG beads for 2 h at 4 °C. The beads were washed extensively with SEC buffer (50 mM PIPES pH 7.2, 50 mM NaCl, 30 µM CuSO_4_, 0.06% digitonin, and either 1 μM ATU or 10 μM DMPP). Proteins were then eluted with SEC buffer containing 200 µg/mL FLAG peptide. The eluent was then loaded onto a Superose S6 3.2/60 column (GE Healthcare) pre-equilibrated with SEC buffer. Peak fractions were collected for cryo-EM sample preparation.

### Lipidomics

Total lipid extracts (TLEs) of all samples were prepared using the Bligh and Dyer method, as modified by Sturt et al ^51^. In brief, frozen *N. maritimus* SCM1 cells or membrane fractions were subjected to two sequential extractions with a methanol (MeOH):dichloromethane (DCM):phosphate buffer (PB) mixture (2:1:0.8, v/v/v), followed by two additional extractions with MeOH:DCM:trichloroacetic acid (TCA) buffer (2:1:0.8, v/v/v). During each extraction step, samples were sonicated for 10 min and then centrifuged at 1,000 × g for 5 min. Following phase separation, the DCM layer was collected, and the residual aqueous phase was further extracted twice with pure DCM. The combined DCM fractions were subsequently dried under a stream of nitrogen and stored at –80°C until analysis.

An aliquot of each TLE was dissolved in MeOH for injection. Archaeal lipid samples were separated using an ACQUITY I-Class Ultra Performance Liquid Chromatography (UPLC) system equipped with a C18 EXCEL UPLC column (2.1 × 150 mm, 2 μm; ACE). The elution gradient was adapted from the protocol described by Zhu et al.^52^: 0–5 min, 100% A; 5–10 min, 0–24% B; 10–36 min, 24–60% B; 36–38 min, 60–90% B; 38–45 min, 90% B; 45–45.1 min, 90–100% A; 45.1–55 min, 100% A. Mobile phase A consisted of MeOH, and mobile phase B consisted of isopropanol; both were supplemented with 0.04% formic acid (>99.0%, Optima™ LC/Mass spectrometry (MS) Grade, Fisher Chemical) and 0.1% ammonium hydroxide (25–30% NH_3_ basis, Sigma-Aldrich). The flow rate was maintained at 0.3 mL/min, and the total run time was 55 min.

MS parameters were configured according to the protocol established by Chen et al.^53^. Data acquisition was performed in FAST-DDA mode. The m/z ranges for MS^1^ and MS^2^ scans were set to 100–2000 and 50–2000, respectively. A ramped collision energy profile—ranging from 10–15 eV at low mass to 55–65 eV at high mass—was applied for collision-induced dissociation (CID) to generate MS^2^ spectra. Mass calibration was performed using a sodium iodide solution (m/z 50–2000; residual mass error < 0.5 ppm), and real-time mass accuracy was maintained by continuous infusion of leucine enkephalin ([M + H]⁺ at m/z 556.2771) as a lock mass. Lock mass correction data were acquired for 0.2 s at 20-s intervals throughout the entire acquisition period.

Raw data files were converted to mzML format using the Waters2mzML script (version 1.2.0) ^54^. Precursor m/z values were subsequently corrected using the mzxml-precursor-corrector script ^55^. Further data processing was conducted using MS-DIAL, with the following settings: minimum peak height of 1000 intensity units, mass slice width of 0.1 Da, retention time tolerance of 0.1 min, and MS^1^ m/z tolerance of 0.05 Da. Lipid annotations were assigned based on a maximum mass error of 0.01 Da for MS^1^ and 0.05 Da for MS^2^, and only features with a total match score exceeding 70% against the reference library ^56^ were considered identified. The feature table exported from MS-DIAL was further processed using a custom Python script to identify potential adducts of annotated features and extract corresponding peak areas, which were then aggregated into lipid compounds for downstream proxy calculations.

### Preparation of cryo-EM samples and data acquisition

For cryo-EM grid preparation, 4 µL of purified *Nh*AMO in complex with NIs, or membrane-bound *Nm*AMO prepared with and without NIs, were applied to freshly glow-discharged gold grids (Quantifoil R2/1, 300 mesh; 10 mA, 10 s) coated with a single layer of graphene (kindly provided by Jiayue Su, Tsinghua University). Grids were blotted for 3.5 s at 100% humidity and 4 °C and plunge-frozen in liquid ethane using a Vitrobot Mark IV device (Thermo Fisher Scientific).

Cryo-EM data were acquired on a Titan Krios G3 (Thermo Fisher Scientific) operating at 300 kV, equipped with a K3 direct electron detector (Gatan) and a BioQuantum energy filter (Gatan) using a 20 eV slit. A total of 15,310, 10,073, 16,914 and 15,382 micrographs were collected automatically for *Nh*AMO-DMP, *Nh*AMO-ATU, active *Nm*AMO and inactivated *Nm*AMO, respectively, in super-resolution mode at a nominal magnification of 130,000×, yielding a calibrated pixel size of 0.668 Å. Each exposure was dose-fractionated into 32 frames with a total dose of 50 e⁻⁄Å². Data acquisition was performed using EPU software ^57^ over a defocus range of –1.5 to –2.0 µm.

### Image processing

The beam-induced motion of each entire micrograph was corrected using MotionCor2 ^58^. All subsequent data processing steps were performed in cryoSPARC version 4.3.2 ^59^. Contrast transfer function (CTF) parameters were estimated by Patch CTF estimation for the AMO datasets, and micrographs with a poor CTF fit, as determined by Manually Curate Exposures, were discarded. Particle picking was carried out using Blob Picker with a diameter range of 8 to 20 nm. Auto-picked particles were extracted with a box size of 224 pixels and binned by a factor of 2. Following multiple rounds of two-dimensional (2D) classifications, good classes were used to train Topaz ^60^. These datasets underwent additional rounds of 2D classification, Topaz training and particle re-extraction; this process was repeated four times until no further improvement in class quality was observed. Particles from all iterations were merged, and duplicate picks were removed. The final particle set was subjected to ab initio 3D reconstruction, followed by heterogeneous refinement, resulting in distinct EM maps. The map featuring the AMO complex was subjected to several rounds of AMO heterogeneous refinement, using previously generated maps as references, to remove poorly aligned or bad particles. The highest-quality map was selected for subsequent non-uniform (NU) refinement ^61^ under C3 symmetry, achieving an overall resolution as reported in Extended Data Table 1. Finally, DeepEMhancer ^62^ was applied for the post-processing of refined maps to enhance features for model building. As a control, we also performed processing under C1 symmetry; this essentially yielded identical structures with only a marginal reduction in resolution.

### Model building and refinement

For archaeal AMO, initial models of *Nm*AmoA–*Nm*AmoF were generated using AlphaFold3 ^63^ based on their respective amino acid sequences. The *Nm*AmoG sequence was identified within the *amo* gene cluster and similarly modeled using AlphaFold3. *Nm*AmoH was modeled as polyA, with the exception of several residues exhibiting large side chains. Archaeal lipids were assigned based on the lipidomics data and the corresponding features of cryo-EM density. The resulting subunit models and lipid components were docked into the cryo-EM map using UCSF Chimera ^64^, followed by manual rebuilding in Coot ^65^. The atomic model was iteratively refined in Phenix ^66^ using the Real-space refinement module with secondary structure and geometric restraints, interspersed with manual adjustments in Coot to optimize stereochemistry and side-chain conformations.

For bacterial AMO in complex with DMP or ATU, our recently resolved structure (PDB 9LEG) was used as the initial template, fitted into the cryo-EM map using Chimera, manually rebuilt in Coot, and refined using Real-space refinement in Phenix. Data collection, refinement and validation statistics are summarized in Extended Data Table 1. Structural figures were prepared using UCSF Chimera, Chimera X ^67^ and PyMOL (http://www.pymol.org).

### Expression and purification of cupredoxin domains and *Nmar*_1504-encoded protein

The DNA sequences encoding the cupredoxin domains of *Nm*AmoB (residues 35– 162), *Nh*AmoB (residues 38–420, excluding the transmembrane helix), and *Nvi*AmoB (residues 25–158) were commercially synthesized by Jiangsu CoWin Biotech Co., Ltd. (Nantong, China) and individually cloned into a modified pET28a (+) vector featuring an N-terminal 3×Flag tag and a C-terminal 6×His tag. Similarly, the coding sequence of *Nmar*_1504 was synthesized and subcloned into a modified pGEX-6P-1 vector bearing an N-terminal 6×His–GST dual affinity tag. All recombinant plasmids were transformed into *Escherichia coli* BL21(DE3) competent cells. Protein expression was induced with 0.2 mM isopropyl-β-D-thiogalactopyranoside (IPTG) at 16 °C for 12 h. Cells were harvested by centrifugation at 5,000 × g for 10 min at 4 °C, then resuspended in lysis buffer (25 mM HEPES, pH 7.4; 500 mM NaCl). Cell lysis was performed by sonication on ice, followed by centrifugation at 12,000 × g for 30 min at 4 °C to remove insoluble debris. The clarified supernatant was incubated with Ni–NTA agarose resin at 4 °C for 30 min under gentle agitation. After binding, the resin was washed sequentially with lysis buffer containing 20 mM, 50 mM, and 80 mM imidazole to eliminate nonspecifically bound proteins. Target proteins were eluted with elution buffer (25 mM HEPES, pH 7.4; 150 mM NaCl) supplemented with 200 mM imidazole. Eluates were pooled, aliquoted, and stored at −80 °C until further use.

### Ammonium binding capacity of archaeal and bacterial cupredoxin-domain proteins

Purified cupredoxin-domain proteins from archaeal and bacterial AMOs were buffer-exchanged into the same assay buffer before analysis. To enable direct comparison of ammonium binding capacity, all protein-containing samples were adjusted to contain the same molar amount of cupredoxin-domain proteins. A protein-free buffer sample was included as a control to correct for background ammonium levels and nonspecific ammonium loss during incubation and ultrafiltration.

NH_4_Cl was added to each protein sample and to the buffer control to a final concentration of 10 μM. After gentle mixing, a 100-μL aliquot was immediately collected from each sample to determine the initial total ammonium concentration at 0 h. The remaining samples were incubated overnight at 4 °C to allow ammonium binding to reach equilibrium. After incubation, each sample was transferred to a 30-kDa molecular-weight-cutoff centrifugal filter unit (Merck, Germany) and centrifuged at 7,000 × g for 15 min at 4 °C. Under these conditions, cupredoxin-domain proteins were retained in the upper chamber, whereas unbound ammonium passed into the filtrate. The filtrate was collected and used to determine the concentration of free, unbound ammonium using the optimized indophenol blue assay described above.

The ammonium binding capacity of each cupredoxin-domain protein was calculated from the decrease in free ammonium concentration after incubation and ultrafiltration, after correction using the protein-free buffer control. Binding capacity was expressed as the amount of ammonium retained by the protein-containing fraction per equal molar amount of cupredoxin-domain protein.

### Isothermal Titration Calorimetry

Isothermal titration calorimetry (ITC) experiments were performed at 25 °C using a Nano ITC instrument (TA Instruments, New Castle, DE, USA). Purified archaeal and bacterial cupredoxin-domain proteins were buffer-exchanged into the same assay buffer before measurement. To enable direct comparison of NH_4_⁺-binding affinity, all protein samples were adjusted to the same molar concentration. For each experiment, the sample cell contained 300 μL of an individual cupredoxin-domain protein at 0.005 mM, whereas the titration syringe contained 0.2 mM NH_4_Cl prepared in the identical assay buffer.

Each titration consisted of twenty consecutive 2-μL injections of NH_4_Cl into the protein solution. The sample cell was stirred at 350 rpm, with a 120-s interval between injections. Control titrations were performed by injecting NH_4_Cl into assay buffer alone under identical conditions, and the resulting heat of dilution was subtracted from the corresponding protein titration data before fitting.

The corrected ITC thermograms were analyzed using the one-site binding model implemented in Origin 7.0 software supplied with the instrument (OriginLab, USA). The dissociation constant, Kd, was calculated as the reciprocal of the fitted association constant, Ka. The ITC titration syringe and sample cell were thoroughly cleaned with mild detergent and water after each run according to the manufacturer’s instructions.

### Substrate-dependent oxygen uptake measurement

Cellular substrate oxidation kinetics were determined from instantaneous substrate-dependent oxygen uptake rates, as previously described ^16,17,68,69^. Oxygen uptake was measured using a microrespiration (MR) system consisting of a 6-mL glass MR chamber fitted with an MR injection lid, a glass-coated magnetic stir bar, a microsensor multimeter, and an OX-MR oxygen microsensor (Rank Brothers Ltd., Cambridge, UK). The oxygen microsensor was polarized continuously for at least 24 h before use.

Active *N. europaea* and *N. halophila* ZJB1 cells were harvested from 1 L of NH_4_^+^-replete cultures by centrifugation at 10,000 × g for 20 min at 20 °C. The resulting cell pellets were washed and resuspended in 20 mL of the corresponding NH_4_^+^-free growth medium. Because *N. maritimus* SCM1 cells lost detectable oxygen-uptake activity after centrifugation or filtration, kinetic measurements for this strain were performed using unconcentrated active cultures, as reported previously ^16,17^. All samples of ammonia-oxidizing microorganisms were equilibrated at 30 °C in a recirculating water bath for at least 20 min before transfer to the MR chamber.

For each measurement, the MR chamber was filled headspace-free with 5 mL of cell cultures, sealed with MR injection lids, and submerged in a recirculating water bath at 30 °C. An OX-MR microsensor was inserted into the MR chamber and allowed to equilibrate for approximately 1 h. Before substrate addition, background sensor drift and endogenous oxygen consumption were recorded for at least 10 min, and this background rate was subtracted from all subsequent oxygen-uptake rates. Substrate-dependent oxygen uptake was initiated by sequential injections of NH_4_Cl solutions using a sterile syringe to generate increasing total ammonium concentrations in the MR chamber. Oxygen concentration was continuously recorded after each injection. Once the signal stabilized, the instantaneous oxygen-uptake rate was calculated from the linear slope of oxygen depletion following each substrate addition. Immediately after each experiment, the total ammonium concentration and pH of the MR chamber contents were measured to determine the actual substrate concentration used for kinetic analysis.

After oxygen-uptake measurements, cells were collected and stored at −20 °C for protein quantification. Cells were lysed using an ultrasonic homogenizer (Xinzhi Instrument, China), and total protein content was determined using a BCA protein assay kit (Beyotime, Shanghai, China) according to the manufacturer’s instructions. The resulting substrate-dependent oxygen-uptake rates were used for subsequent calculation of apparent kinetic parameters, including *K*_m(app)_ and *V*_max_, as described below.

### Calculation of kinetic properties

The apparent half-saturation constant (*K*_m(app)_) and maximum reaction rate (*V*_max_) were calculated from substrate-dependent oxygen uptake measurements. Background-corrected oxygen uptake rates were converted to total ammonium oxidation rates using the stoichiometric ratio of total ammonium (NH_3_ + NH_4_^+^) oxidized to oxygen consumed of 1:1.5, as described previously ^16,17,68^. The resulting total ammonium uptake rates were fitted to the Michaelis-Menten model using Origin 7.0 software (OriginLab, USA) according to the following equation:

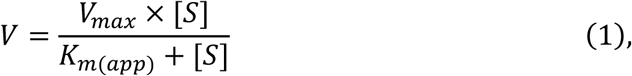

where *V* is the total ammonium oxidation rate (μM h⁻¹), *V*_max_ is the maximum oxidation rate (μM h⁻¹), [*S*] is the total ammonium concentration (μM), *K*_m(app)_ is the reaction half-saturation concentration for total ammonium (μM). A nonlinear least squares regression analysis was used to estimate *K*_m(app)_ and *V*_max_. The *K*_m(app)_ for NH_3_ for each strain was calculated based on the *K*_m(app)_ for total ammonium, incubation temperature, salinity and pH ^70^. The *V*_max_ values of the pure cultures were normalized according to the protein content of the cultures.

### Microcosm enrichment assay comparing the competitive responses of AOA and AOB

Surface water was collected from the Shenzhen Bay site SZB1 (22.5242°N, 113.9861°E) and used to establish microcosm enrichment experiments under different ammonium regimes. Water samples were dispensed into 2-L Duran bottles and amended with NH_4_Cl to final total ammonium concentrations of 1 μM or 10 μM, representing low- and higher-ammonium conditions, respectively. Three biological replicates were established for each treatment. The bottles were sealed with breathable cotton stoppers and incubated in the dark at 25 °C for three weeks in temperature-controlled chambers. During incubation, the bottles were gently shaken to maintain oxygen availability for microbial growth.

The NH_4_^+^ concentration in each microcosm was monitored daily. When NH_4_^+^ was depleted, NH_4_Cl was replenished to restore the designated treatment concentration, thereby maintaining the intended ammonium regime throughout the enrichment period. Initial in situ water samples and enriched samples collected after three weeks were filtered through 0.22-μm-pore-size polycarbonate membranes (Millipore). The filters were immediately frozen and stored at −80 °C until DNA extraction.

DNA was extracted from the filters using a soil genomic DNA extraction kit (Tiangen, China) according to the manufacturer’s instructions. The abundances of AOA and β-AOB were quantified by quantitative PCR targeting archaeal and bacterial *amoA* genes, respectively, using the primers listed in Extended Data Table 2. qPCR reactions were performed on a QuantStudio 5 Real-Time PCR system (Applied Biosystems, CA, USA). Each 20-μL reaction contained 10 μL ChamQ™ Universal SYBR qPCR Master Mix (Vazyme, Nanjing, China), 0.5 μL forward primer, 0.5 μL reverse primer, 1 μL DNA template, and nuclease-free water. All samples and standard reactions were analyzed in triplicate. The resulting *amoA* gene abundances were used to compare the competitive responses of AOA and β-AOB under low- and higher-ammonium enrichment conditions.

### Quantitative proteomics and AMO abundance normalization

### Protein extraction and peptide preparation

Total cellular proteins were extracted from growing *N. maritimus* SCM1, *N. halophila* ZJB1, and *N. europaea* cells. Cell pellets were resuspended in 200 µL RIPA lysis buffer (Beyotime Biotechnology, Shanghai, China) and disrupted with steel beads at 45 Hz for 2 min, repeated once, followed by sonication in an ice-water bath for 20 min. Lysates were centrifuged at 12,000 × g for 10 min at 4 °C, and the supernatants were collected for total-protein quantification. Protein concentrations were determined using a bicinchoninic acid (BCA) assay with BSA standards (Beyotime Biotechnology). For each sample, 30 µg of total protein was subjected to reduction, alkylation, and enzymatic digestion using a commercial sample-preparation kit (Magigene, Guangzhou, China; reagents A to J) according to the manufacturer’s instructions. Briefly, samples were incubated with reagent H, reduced at 95 °C for 20 min, alkylated with reagent I at 37 °C for 10 min, and digested at 37 °C for 2.5 h. The resulting peptides were desalted on a desalting plate, eluted with reagent F, vacuum-dried, and stored at −20 °C until LC-MS/MS analysis.

### LC-MS/MS proteomic analysis

Peptide samples were analyzed using a Vanquish Neo nano-UPLC system coupled to an Astral Zoom mass spectrometer (Thermo Scientific, USA) equipped with a nano-electrospray ion source. Peptides were separated on an EASY-Spray reversed-phase analytical column (150 µm × 15 cm; Thermo Scientific, USA). Mobile phase A consisted of water with 0.1% formic acid, and mobile phase B consisted of 80% acetonitrile with 0.1% formic acid. Data-independent acquisition (DIA) was performed in positive-ion mode. The electrospray voltage was set to 1,800 V and the ion-transfer temperature to 280 °C. MS1 scans were acquired over an m/z range of 380 to 980 at a resolution of 240,000, with an AGC target of 500% and a maximum injection time of 3 ms. MS2 scans were acquired over an m/z range of 150 to 2,000 with a maximum injection time of 2.5 ms, a 2-Th isolation window, an RF lens setting of 40%, higher-energy collisional dissociation (HCD) activation, a normalized collision energy of 25%, and a cycle time of 0.6 s.

### Database search and protein quantification

Vendor raw files were processed with DIA-NN software (version 2.2.0). Spectra were searched against species- or strain-specific protein FASTA databases, with carbamidomethylation of cysteine specified as a fixed modification and methionine oxidation and protein N-terminal acetylation specified as variable modifications. Trypsin was selected as the digestion enzyme, allowing up to two missed cleavages. The false discovery rate (FDR) threshold was set to 0.01 at the PSM and peptide levels. Initial precursor and fragment mass tolerances were both set to 20 ppm. All other parameters were kept at the DIA-NN default settings. Protein-group quantities exported from DIA-NN were used for downstream abundance calculations.

### Estimation of cellular AMO abundance and activity normalization

To compare ammonia oxidation activity after accounting for differences in AMO expression, cellular AMO abundance was estimated from the DIA-NN protein-group quantities. For each strain and replicate, the summed quantity of annotated AMO-complex subunits was divided by the summed quantity of all quantified protein groups in the same sample, yielding the AMO protein fraction of the total quantified proteome. For archaeal *N. maritimus* SCM1, AMO-complex subunits included the structurally assigned *Nm*AmoA to *Nm*AmoG subunits when quantified. For bacterial *N. halophila* ZJB1 and *N. europaea*, annotated AMO subunits were included according to the corresponding genome annotations. The AMO fraction was calculated independently for each replicate and then averaged for each organism.

Bulk ammonia oxidation rates (AORs) measured from matched cultures were normalized to cellular AMO abundance using the following relationship:

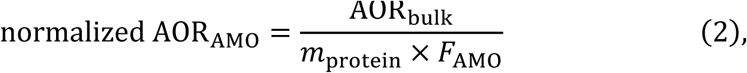

where normalized AOR_AMO_ is the AORs normalized to the protein content of pure cultures (μmol N mg protein^-1^ h^-1^), AOR_bulk_ is the bulk AORs (μmol L^-1^ h^-1^), *m*_protein_ is the total protein content of pure cultures, *F*_AMO_ is the AMO protein fraction estimated from quantitative proteomics. Based on quantitative proteomics, the *F*_AMO_ values for AOA (*N. maritimus* SCM1) and AOB (*N. europaea* and *N. halophila* ZJB1) were approximately 3 and 5% of the total protein respectively, consistent with previous proteomic estimates ^71,72^. This normalization was used to compare the apparent AMO abundance-normalized activities of *N. halophila* ZJB1, *N. europaea*, and *N. maritimus* SCM1. Data are presented as mean ± SD from biological replicates, and statistical comparisons were performed as described for Fig. 2g.

### Copper quantification by inductively coupled plasma mass spectrometry (ICP-MS)

Purified AMO samples were dried using a vacuum centrifugal concentrator (Jiaimu, Beijing, China). The dried samples were accurately weighed to 0.0001 g and transferred into acid-cleaned polytetrafluoroethylene digestion vessels. Concentrated nitric acid and hydrochloric acid were added at a ratio of 3:1 (v/v; 6 ml HNO_3_ and 2 ml HCl), and the samples were pre-digested at room temperature for approximately 30 min. Digestion was then performed on a hotplate using a stepwise temperature program: 100 °C for 10 min, 120 °C for 10 min, and 200 °C for 60 min. Additional acid mixture was added when necessary to prevent complete drying during digestion. After complete digestion, the solution was gently evaporated to approximately 0.5 ml to remove excess acid, cooled to room temperature in a fume hood, and quantitatively transferred to a 10-ml volumetric flask. The digestion vessel was rinsed three times with metal-free water, and the rinses were combined with the digest. The final volume was adjusted to 10 ml with metal-free water and mixed thoroughly. Copper content was determined using an Agilent 7500CE ICP-MS instrument (Agilent Technologies, USA). Copper stoichiometry was calculated by normalizing the measured copper concentration to the molar concentration of AMO protomers used for digestion (Extended Data Table 3).

### Protein thermal stability (PTS) assay

The thermal stability of purified AMO proteins was assessed using a fluorescence-based differential scanning fluorimetry (DSF) assay with SYPRO Orange dye as the hydrophobic fluorescent probe ^73,74^. Purified archaeal *Nm*AMO and bacterial *Nh*AMO samples were prepared under identical buffer conditions before analysis. Both proteins were adjusted to a final concentration of approximately 0.2 mg mL^-1^ in assay buffer containing 50 mM PIPES, pH 7.2, 50 mM NaCl, and 30 μM CuSO_4_. Each 20-μL reaction contained AMO protein, 10× SYPRO Orange dye (Invitrogen, USA), and assay buffer. Reactions were assembled in 96-well optical PCR plates. Buffer-only wells containing SYPRO Orange but no protein were included as background controls. All measurements were performed using a QuantStudio 5 Real-Time PCR system (Applied Biosystems, CA, USA), with three biological replicates and technical triplicates for each condition. After plate sealing with optical film and brief centrifugation to remove bubbles, samples were heated from 10 to 99 °C at a ramp rate of 0.05 °C s⁻¹. Fluorescence was recorded at 0.5 °C intervals using excitation and emission wavelengths of 490 and 530 nm, respectively. Raw fluorescence signals were corrected by subtracting the buffer-only background. The apparent melting temperature (Tm) of each AMO sample was determined by fitting the thermal unfolding curve to a Boltzmann sigmoidal equation using GraphPad Prism software version 10.0 (GraphPad Software, CA, USA). The resulting apparent Tm values were used to compare the relative thermal stability of *Nm*AMO and *Nh*AMO under identical assay conditions.

### System setup and molecular dynamics simulations

All molecular dynamics (MD) simulations were performed using the Amber Package, version 22 (University of California, San Francisco, CA) ^75^. The initial models of both active and inactivated *Nm*AMO determined in this study were employed. For the titratable residues (His, Asp, Glu), protonation states were assigned based on pKa values calculated in the PlayMolecule website (https://open.playmolecule.org) ^76^ and detailed visual examination of the regional hydrogen-bonded networks. Within the active *Nm*AMO monomer, His9 and His121 (in *Nm*AmoA), His132 (in *Nm*AmoB), His48, His61, His67, His120 and His133 (in *Nm*AmoC), and His55 and His73 (in *Nm*AmoF) were protonated at the *δ* position, whereas His35 and His130 (in *Nm*AmoB), and His67 (in *Nm*AmoD) were protonated at the *ε* position. For Asp and Glu residues, Asp40 and Asp44 (in *Nm*AmoC) were protonated, while the remaining residues were deprotonated. Within *Nh*AMO, His41, His116, His171, His189 and His257 (in *Nh*AmoA), His69, His77, His113, His141, His144, His263, His336, and His405 (in *Nh*AmoB), His54, His74, His140, His153, His211, and His225 (in *Nh*AmoC) were protonated at the *δ* position, whereas His19, His43, and His214 (in *Nh*AmoA), His 38, His142, and His197 (in *Nh*AmoB) were protonated at the *ε* position. For Asp and Glu residues, Asp353 (in *Nh*AmoB), Glu103 and Glu267 (in *Nh*AmoA), and Glu355 and Glu400 (in *Nh*AmoB) were protonated; the remaining residues were deprotonated. The membrane environment of *Nh*AMO was constructed using the Packmol-Memgen program ^77^, with the phospholipid bilayer consisting of 60% phosphatidylvinylethanolamine (PVPE), 20% phosphatidylvinylglycerol (PVPG), and 20% 1-palmitoyl-2-oleoyl-*sn*-glycero-3-phosphocholine (POPC). The Amber ff14SB ^78^, Lipid21 ^79^, and GAFF ^80^ force fields were applied to parameterize the amino acid residues within AMO, phospholipid bilayer, and reductants CoQ10H_2_, respectively, while the force field parameters for the Cu_B_, Cu_C_ and Cu_D_ sites were tailored using the “MCPB.py” tool ^81,82^ of AmberTools23. The membrane-free simulation system was solvated with a periodic rectangular box (the volume is 130.464 × 129.651 × 121.353 Å^3^) containing 42,841 TIP3P water molecules and an approximate number (6) of sodium counterion to neutralize the charge. The membrane-embedded simulation system was solvated with a periodic rectangular box (the volume is 153.531 × 153.531 × 156.643 Å^3^) containing 81,910 TIP3P water molecules, 666 lipid molecules, and 0.15 M NaCl.

Following appropriate setup, the complex systems were successively fully minimized by combining 10,000 steps of the steepest descent method and 10,000 steps of the conjugate gradient method. Then, each system was gradually heated from 0 K to 300 K for a total of 50 ps using the NVT ensemble. To achieve a uniform density after heating dynamics, 1 ns of density equilibrium was executed under the NPT ensemble, during which the temperature and pressure of the system were maintained at 300 K and 1.0 ATM using the Langevin thermostat and the Berendsen barostat ^83,84^, respectively.

Subsequently, all complexes were equilibrated for 2 ns without any restraints in order to relieve minor unfavorable ligand-protein steric interactions that might have been still present. Finally, a productive MD simulation of 200 ns was carried out under the NPT ensemble. Throughout the process, the simulation integration step was set to 2 fs, and the conformational configurations of the complex systems were saved every 100 ps in trajectory files. The covalent bonds connecting hydrogen atoms were then restricted with the SHAKE algorithm ^85^. The short-range nonbonded interactions were adopted using a cutoff radius of 8 Å, while the long-range electrostatic interactions were modeled utilizing the Particle mesh Ewald (PME) method with a grid point density of 0.1 nm and an interpolation order of 4 ^86^. The results of MD simulations were presented by using the Visual Molecular Dynamics version 1.9.3a ^87^, and PyMOL software was used to exhibit the graphics of the MD simulations.

### QM Calculations

To evaluate the reaction mechanisms and energetics of the Cu_C_, Cu_D_, and Cu_C_-Cu_D_ sites in the oxidation of NH_3_ to NH_2_OH (Extended Data Figs. 3c, 3d, 4i and 10), density functional theory (DFT) calculations were carried out using the Gaussian 16 software package (https://gaussian.com/citation/). Geometry optimizations were performed at the MN15/def2-SVP and B3LYP/def2-SVP levels of theory in conjunction with the SMD implicit solvation model ^88^, followed by single-point energy refinements at the B3LYP/def2-TZVP and MN15/def2-TZVP levels. Given that the AMO active site is embedded within the protein–membrane environment, chlorobenzene was selected as the solvent in the SMD model ^88^ to approximate the hydrophobic milieu. The B3LYP functional has been demonstrated to provide a reliable description of mononuclear ^89^ and certain binuclear copper systems ^90^, and dispersion effects were included using Grimme’s D3 correction ^91,92^. In parallel, MN15 was employed based on a comprehensive benchmark by Odoh et al., in which 19 density functionals were evaluated for multicopper systems, revealing that MN15 yields the lowest mean absolute deviation (1.2 kcal mol⁻¹) relative to DLPNO-CCSD(T) calculations extrapolated to the complete basis set limit ^93^. Considering these results, both MN15 (Extended Data Figs. 3c, 3d and 4i) and B3LYP (Extended Data Fig. 10) were applied to assess the ammonia oxidation pathway, with all reported energies further refined using the larger def2-TZVP basis set for all atoms. The energy data are shown in Extended Data Tables 4 and 5, the spin density data are shown in Extended Data Table 6.

### Umbrella sampling

Umbrella sampling simulations were performed to characterize the free energy landscapes associated with long-range molecular events in *Nh*AMO and *Nm*AMO, including the approach of the dinuclear copper center in *Nh*AMO, the entry and dissociation of CoQ10H_2_ in membrane-bound *Nh*AMO, and the dissociation of the inhibitor from the active sites of *Nm*AMO and *Nh*AMO. Umbrella sampling windows were generated along each reaction coordinate at 0.2 Å intervals. For each window, 10 ns of MD simulations were performed under a harmonic biasing potential with a force constant of 50 kcal/mol/Å^2^. The unbiased potentials of mean force (PMFs) were then reconstructed using the weighted histogram analysis method (WHAM) ^94^, yielding quantitative free-energy landscapes for dinuclear copper center approach and inhibitor dissociation.

### QM/MM-MD and metadynamics simulations

All QM/MM Born−Oppenheimer MD simulations were performed utilizing the CP2K program version 2024.1 (https://www.cp2k.org). The initial structures for QM/MM MD simulation were derived from the representative conformations extracted from the classical MD trajectories. The interaction energies of the QM and MM regions were calculated separately using QUICKSTEP ^95^ and the FIST module within the CP2K program, and the real-space multigrid technique ^96^ was employed to calculate the electrostatic coupling between the QM and MM regions. The QM region was processed by using a mixed Gaussian and plane wave (GPW) basis set at the DFT (MN15) level ^97^, and the MM region was parametrized in the same way as the classical MD simulation. All QM regions comprised the dinuclear copper active site and the side chains of the residues Asp136, His153, His140, Asn207, His211 and His225. Specifically, for the binding mode of CoQ10H_2_ within the dinuclear copper active site, the Cu_D_(II)−O_2_^•–^ species and the hydroxy-substituted aromatic ring of CoQ10H_2_ were incorporated into the QM region. For the process of (*μ*-superoxo)Cu_C_(II)Cu_D_(I) and (*μ*-hydroperoxo)Cu_C_(II)Cu_D_(I) intermediate formation, the Cu_D_(II)−O_2_^•–^ and Cu_D_(II)−OOH^−^ species were incorporated into the QM region, respectively. All QM atoms at the boundary employed hydrogen-capped atoms to fill the empty valence. The wave function was expanded using a Gaussian double-ζ valence-polarized basis set (DZVP) ^98^. An auxiliary plane-wave basis set with a cutoff energy of 360 Ry was implemented to converge the electron density, and combined with the Goedecker−Teter−Hutter (GTH) pseudopotential to handle the core electrons ^99^. All QM/MM-MD simulations of 50 ps were performed under the NVT system using 0.5 fs integration steps and the simulated temperature was controlled by canonical sampling through velocity rescaling (CSVR) with a time constant of 10 fs ^100^.

Furthermore, the well-tempered metadynamics method ^101,102^ was used to explore the free energy profile for the process of (*μ*-superoxo)Cu_C_(I)Cu_D_(II) and (*μ*-hydroperoxo)Cu_C_(II)Cu_D_(I) intermediate formation. The collective variables were defined as the distance between Cu_C_ site and the distal oxygen of Cu_D_(II)−O_2_^•−^ and Cu_D_(II)−OOH^−^, respectively. Gaussian-shaped potential hills were defined by a width of 0.1 Å, and a height of 0.6 kcal/mol, with a deposition interval of 10 fs.

### RAMD simulations

Random acceleration molecular dynamics (RAMD) simulations were carried out using GROMACS version 2024.1 to identify and characterize potential inhibitor dissociation pathways from the active sites of *Nm*AMO and *Nh*AMO. In RAMD, a randomly oriented external force is applied to the inhibitor’s center of mass, whereas the remainder of the system evolves under standard MD conditions ^103^. The force direction was maintained for a predefined number of MD steps (N). After each interval of N steps, the displacement of the inhibitor is evaluated. If the inhibitor moves less than a predefined minimum distance (r_min_), a new force direction was randomly assigned; otherwise, the same direction was retained for an additional N steps. This protocol enables unbiased exploration of possible dissociation pathways ^104^. In the present study, a random acceleration of 15 kcal/mol/Å was applied. The parameter N was set to 1,000 MD steps, and r_min_, defined as the distance between the inhibitor and the Cu_D_ site, was set to 10 Å. Twenty snapshot structures extracted from equilibrated classical MD trajectories were used as starting conformations for the RAMD simulations. For each snapshot, 20 independent RAMD trajectories were generated using different random number generator seeds, resulting in a total of 800 RAMD simulations for inhibitor dissociation from the active sites of *Nm*AMO and *Nh*AMO. Dissociation times from individual trajectories were compared to identify the most probable and kinetically favorable dissociation channels for *Nm*AMO and *Nh*AMO.

### QM/MM calculations

The last snapshot extracted from MD or QM/MM MD trajectories was used for the QM/MM calculations. All QM/MM calculations were performed using ChemShell ^105,106^, combining turbomole ^107^ for the QM region and DL_POLY ^108^ for the MM region. The electrostatic embedding scheme ^109^ was used to account for the polarizing effect of the protein environment on the QM region. Hydrogen link atoms with the charge-shift model were applied to treat the QM/MM boundary. During QM/MM geometry optimizations, the QM region was studied with the hybrid UMN15 ^93,110,111^ density functional with two levels of theory. For geometry optimization, the double-ζ basis set def2-SVP were used. The energies were further corrected with the larger basis set def2-TZVP for all atoms. Dispersion corrections computed with Grimme’s D3 method ^92,112,113^ were included in all QM calculations. The transition states (TSs) were determined as the highest point of potential energy surface along the reaction coordinates, and a small increment of 0.02 Å was used for scanning the transition states. All the TSs were obtained from the finely scanned energy surfaces. All the minima were optimized without symmetry restraints. The DL-FIND ^114^ optimizer was used in the geometry optimization. For the first stage, corresponding to HAT from CoQ_10_H_2_ to the Cu_D_(II)–O_2_^•⁻^ species, the QM region comprised the polar headgroup of CoQ_10_H_2_, the Cu_D_(II)–superoxo moiety, and its directly coordinated residues Asn207, His211, and His225. In the second stage, describing the formation of the binuclear *μ*-*η*²:*η*²-peroxo-dicopper(II) intermediate, the QM region was expanded to include, in addition to the Cu_D_(II)–OOH⁻ unit, the bridged Cu_C_(II) center together with its coordinating residues Asp136, His140, and His153. For the initial Cu_D_(II)–OOH⁻ formation step, QM/MM calculations were performed for both the open-shell singlet and triplet spin states. In contrast, the subsequent *μ*-*η*²:*η*²-peroxo-dicopper(II) formation was validated exclusively on the doublet spin surface. The computed relative energies were electronic energies from the QM/MM calculation (Extended Data Table 7). The previous work in other metalloenzymes confirmed that the electronic energy barrier is close to the free energy barrier ^115–122^. The spin density of QM/MM simulated species are shown in Extended Data Table 6.

### Constructions of mutant ammonia-oxidizing microorganisms

Mutated plasmid DNA containing point mutations at specific amino acid residues associated with inhibitors were generated following standard molecular cloning procedures. PCR amplification was carried out using the primers listed in Extended Data Table 8, and the resulting products were purified. A plasmid carrying the AMO-encoded genes and conferring kanamycin (KmR) and ampicillin (AmpR) resistance was constructed in our laboratory and linearized using the Pjtu-F1/R1 primer pair. Subsequently, PCR was performed separately with the corresponding site-directed mutagenesis primers and the Amp-F1/R1 primer pair. The resulting products were purified using a DNA purification kit (Shenggong Biotech, Shanghai). The fragments were then ligated using a ligation kit (Vazyme, Nanjing) and transformed into DH5α competent cells. The transformed cells were spread onto LB agar plates containing 50 μg/mL kanamycin and incubated statically at 37 °C. After 24 h, single colonies were picked and inoculated into liquid LB medium supplemented with 50 μg/mL kanamycin, followed by incubation at 37°C with shaking at 180 rpm for 16 h. Finally, the plasmid DNA was extracted and introduced into wild-type *N. halophila* ZJB1 cells via electroporation following a standard protocol ^123^. Transformants were subsequently selected on agar plates supplemented with 25 mg/L kanamycin, yielding seven kanamycin-resistant mutant strains.

### Activity assays of ammonia-oxidizing microorganisms

Ammonia oxidation activity was assessed by monitoring NH_4_⁺ consumption in AMO-containing strains with mononuclear copper centers (*N. maritimus* SCM1 and *N. europaea*) and a binuclear copper center (*N. halophila* ZJB1). Briefly, batch cultures of the three strains were grown to the late exponential phase. Each culture was then adjusted to an OD_600_ of approximately 0.07 (with AOB mutant strains adjusted to approximately 0.1), transferred into 20 mL of their respective medium and supplemented with NH_4_⁺ to a final concentration of ∼2 mM along with other essential nutrients. *N. maritimus* SCM1 was grown without agitation, whereas AOB strains were grown with shaking at 120 rpm; all strains were incubated at 30°C. Aliquots were collected at predetermined time intervals to measure NH_4_⁺ concentration and evaluate ammonia oxidation activity in each strain. ammonia oxidation rates were determined by NH_4_⁺ consumption during the initial 12-h (for AOB) or 24-h (for AOA) incubation period.

### Concatenated marker gene phylogeny

A set of representative AOA or AOB genomes (10 and 6 taxa, respectively), encompassing currently known lineages, were used for phylogenomic analysis. Phylogenetic trees were inferred using a concatenated set of 122 archaeal-specific single-copy marker genes and 120 bacterial-specific marker genes in GTDB (https://gtdb.ecogenomic.org/). Orthologs of these marker genes in the AOA or AOB metagenome-assembled genomes and reference genomes were identified using the GTDB-Tk tool ^124^ based on hidden Markov models. Maximum-likelihood trees were constructed with IQ-TREE ^125^ using the following command: “-m LG+F+I+G4, -bb 1000” for AOA, and “-m JTT+F+I+G4, -bb 1000” for AOB. Trees were edited using iTOL (https://itol.embl.de/), using the non-ammonia-oxidizing *Thaumarchaeota* and *Nitrosospira* lineages as outgroups for the AOA or AOB phylogenomic trees, respectively. The final trees were refined in Adobe Illustrator.

---

**Extended Data Figure 1.**
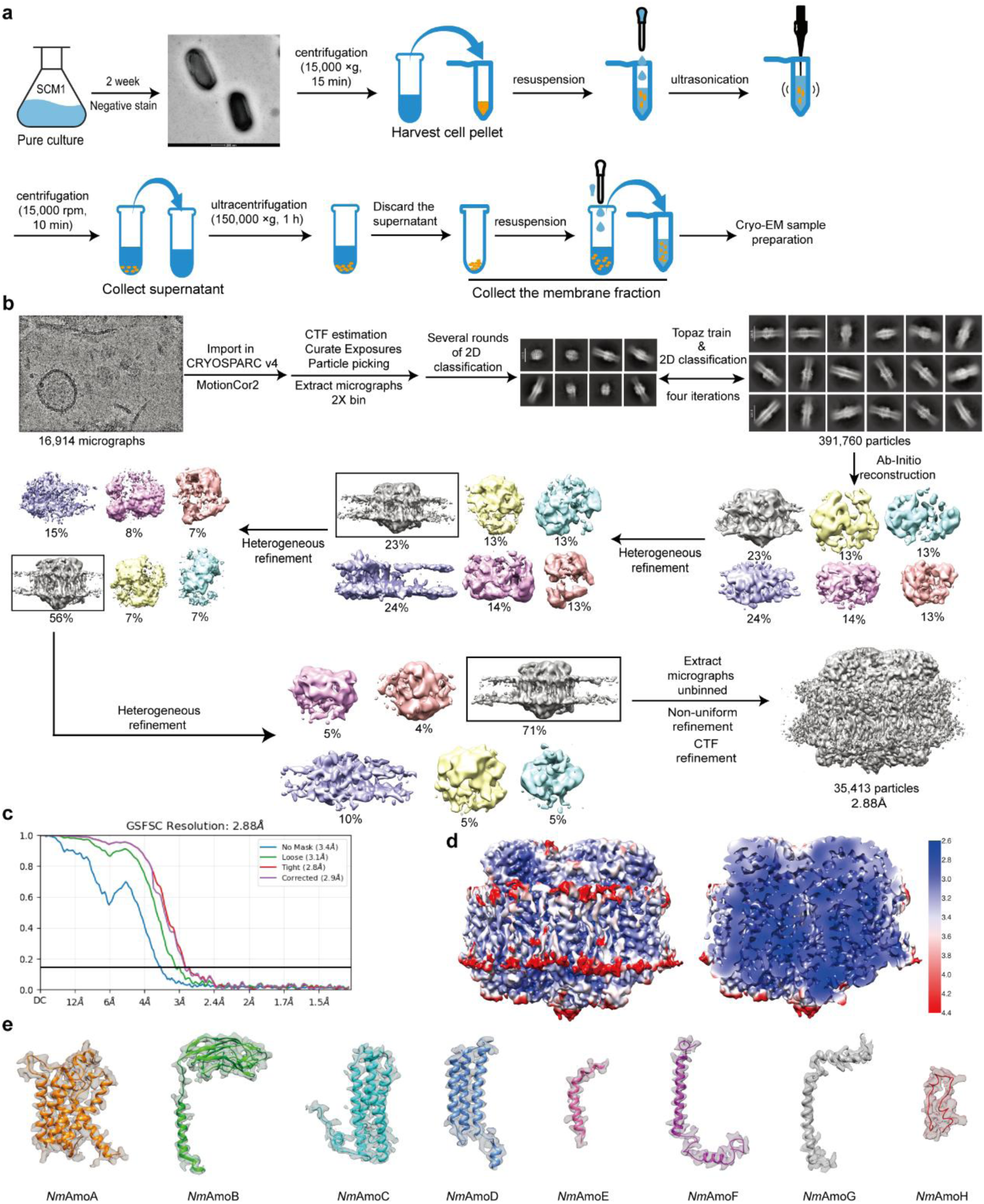
Preparation and cryo-EM data analysis of archaeal AMO. **a.** Schematic diagram depicting the extraction of archaeal membranes. **b.** The flowchart of cryo-EM data processing of active *Nm*AMO. Details can be found in the ‘Method’ section. **c.** FSC curves for the cryo-EM map of active *Nm*AMO. The threshold of 0.143 was used to determine the overall resolution of the map. **d.** Local resolution maps for the overall reconstruction (left) and a central slice (right) of active *Nm*AMO. **e.** Representative EM maps for subunits from active *Nm*AMO.

**Extended Data Figure 2.**
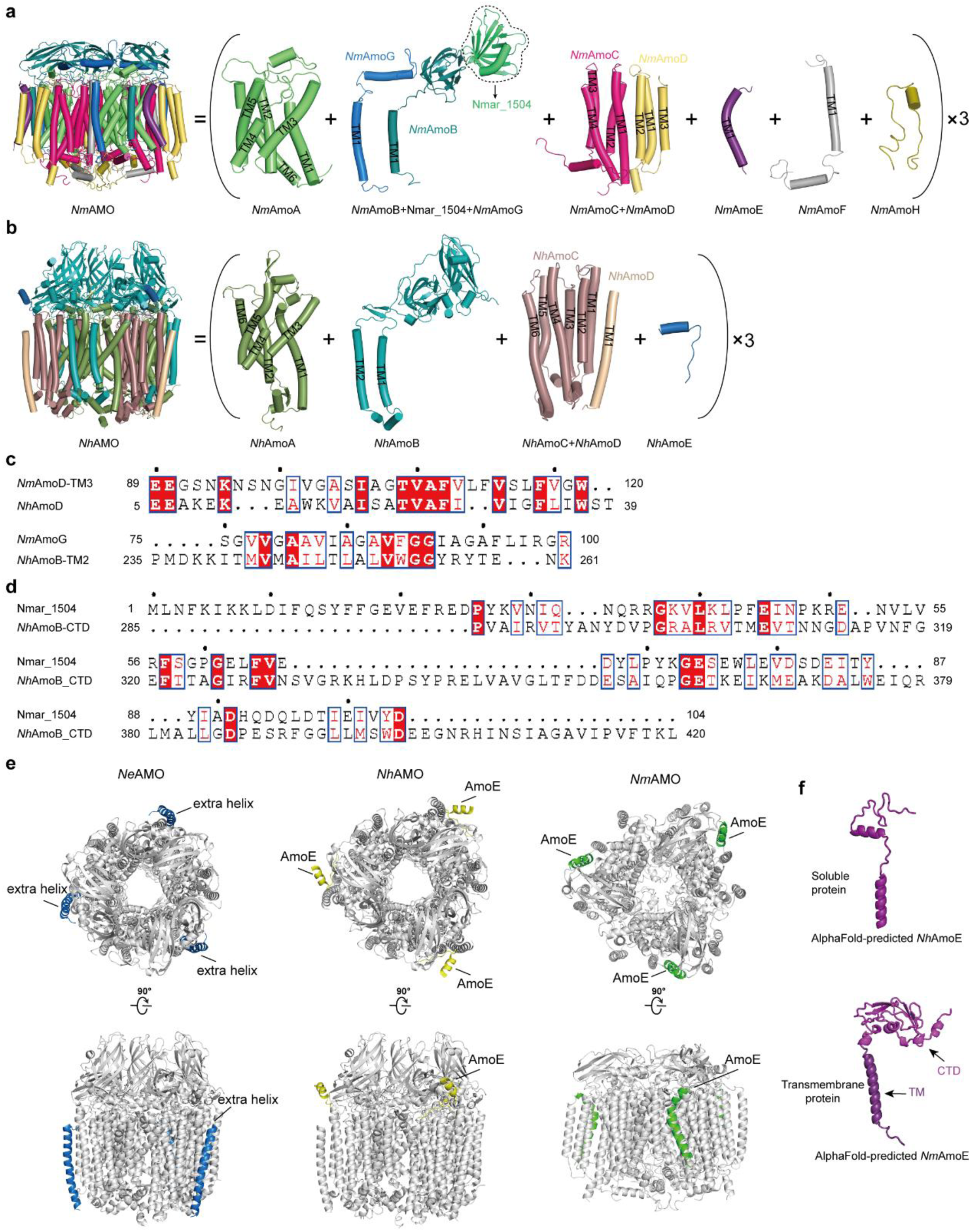
Structural and compositional comparison of archaeal and bacterial AMO. a-b. Comparison of the 3D structures and subunit composition of *Nm*AMO (a) and *Nh*AMO (b). Subunits within an AMO protomer are shown as cartoon and color-coded. Nmar_1504, corresponding to the C-terminal cupredoxin domain of *Nh*AmoB, is indicated by dashed line. c. Sequence alignments of TM3 of *Nm*AmoD and *Nh*AmoD (upper panel), and of *Nm*AmoG and TM2 of *Nh*AmoB (lower panel), revealing potential evolutionary conservation. d. Sequence alignment of the Nmar_1504 and C-terminal cupredoxin domain of *Nh*AmoB. e. Comparison of AmoE and related accessory helices in AMO structures. Top and side views of *Ne*AMO, *Nh*AMO, and *Nm*AMO highlighting the positions of accessory helical elements. The extra helices in *Ne*AMO (PDB 9CL6), bacterial AmoE in *Nh*AMO (PDB 9LEG), and archaeal AmoE in *Nm*AMO (PDB 9XJ2) occupy distinct positions and display different topologies, indicating that archaeal AmoE is not directly equivalent to the supernumerary helical element reported in NeAMO. f. Structural comparison of *Nh*AmoE and *Nm*AmoE.

**Extended Data Figure 3.**
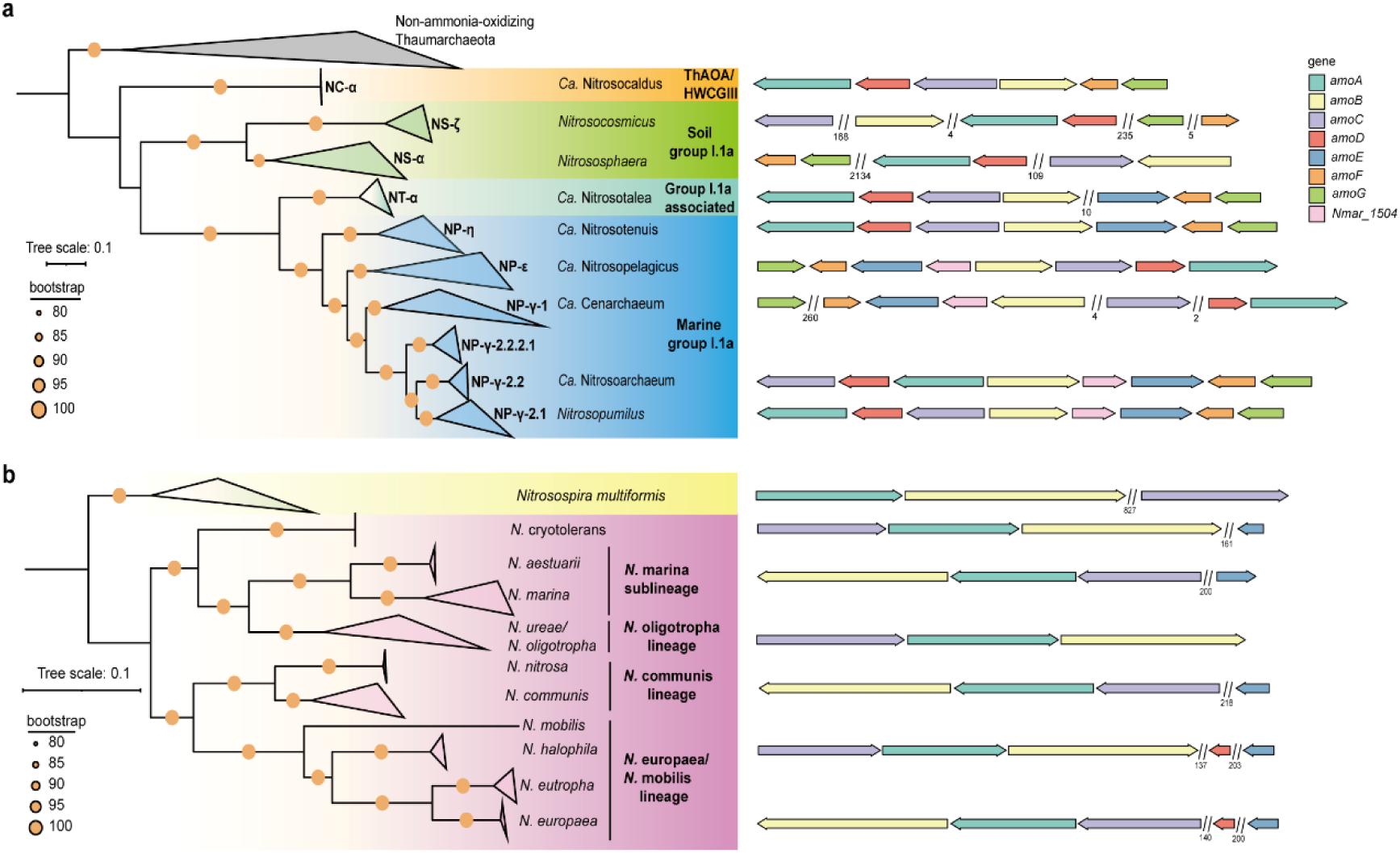
Genomic synteny of AMO subunits in AOA (a) and AOB. **(b).** Left: Maximum-likelihood trees of AOA **(a)** and AOB **(b)**. Nodes with ultrafast bootstrap values ≥80% are marked with orange circles. Clades in bold were included in syntenic analysis. Clade nomenclature follows Alves et al. (2018)^126^ for AOA and Purkhold et al. (2003)^127^ for AOB. Right: Representation of general syntenic patterns in different clades of AOA **(a)** and AOB **(b)**. Gaps between genes on the same contig are marked by a double forward slash. Numbers under the double forward slash represent number of genes between amo subunit genes.

**Extended Data Figure 4.**
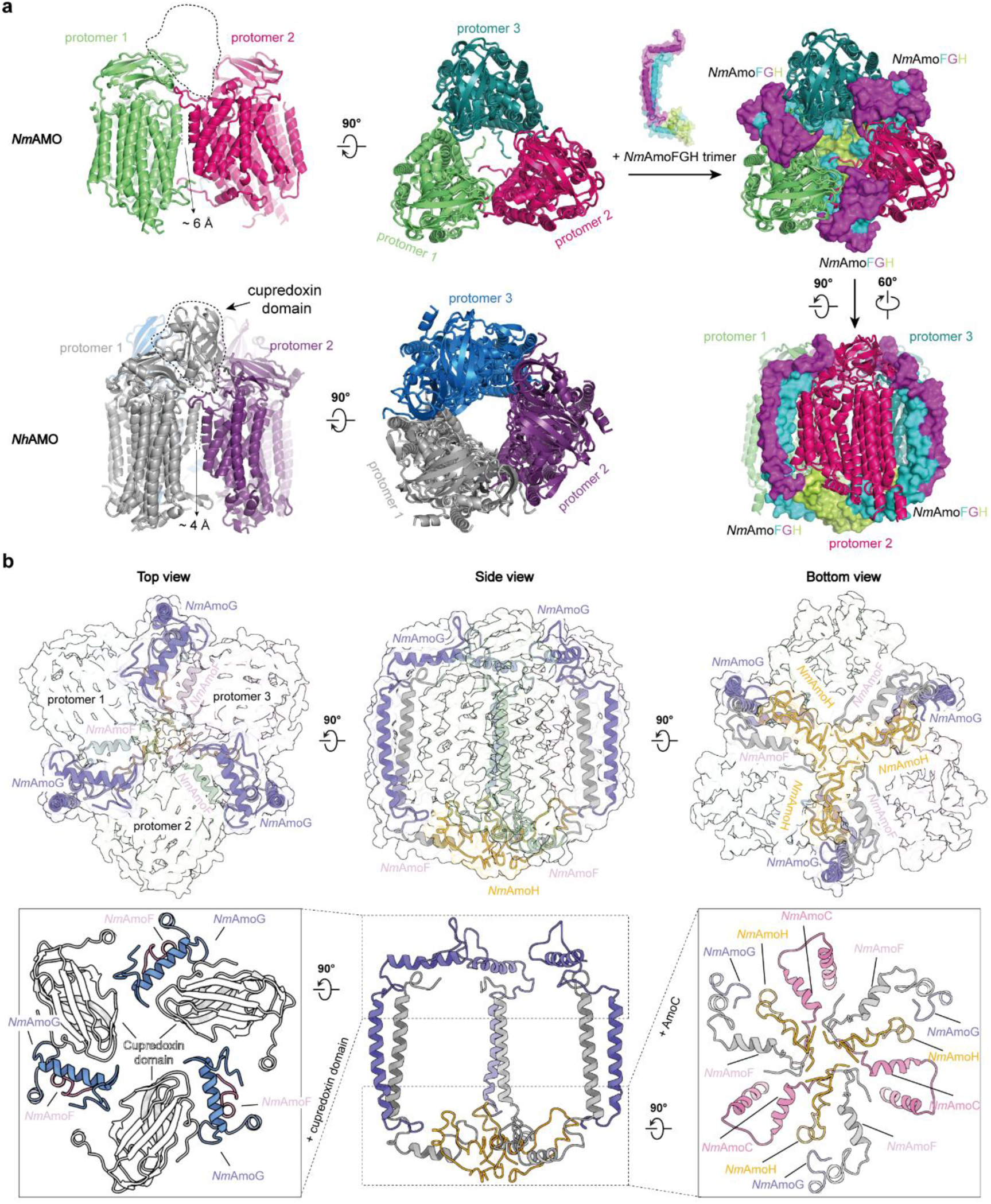
*Nm*AmoFGH stabilizes the *Nm*AMO structure. a. Structural comparison of *Nm*AMO and *Nh*AMO revealed that *Nm*AmoFGH heterotrimers stabilized *Nm*AMO structure through mediating interactions between adjacent protomers. In *Nh*AMO, the C-terminal cupredoxin domain of *Nh*AmoB serves as a major interface for inter-protomer interaction, whereas this structural element is absent in *Nm*AMO (indicated by dashed lines), resulting in a larger distance between adjacent protomers in *Nm*AMO in the absence of *Nm*AmoFGH compared to *Nh*AMO. b. The *Nm*AmoFG heterodimer formed a clamp-like structure that stabilized the *Nm*AMO complex. The N-termini of *Nm*AmoF and *Nm*AmoG mediated interactions between the periplasmic domains of adjacent protomers, while their C-termini mediated interactions between cytoplasmic domains and associated with AmoH to form a closed base for the *Nm*AMO complex in the cytoplasm.

**Extended Data Figure 5.**
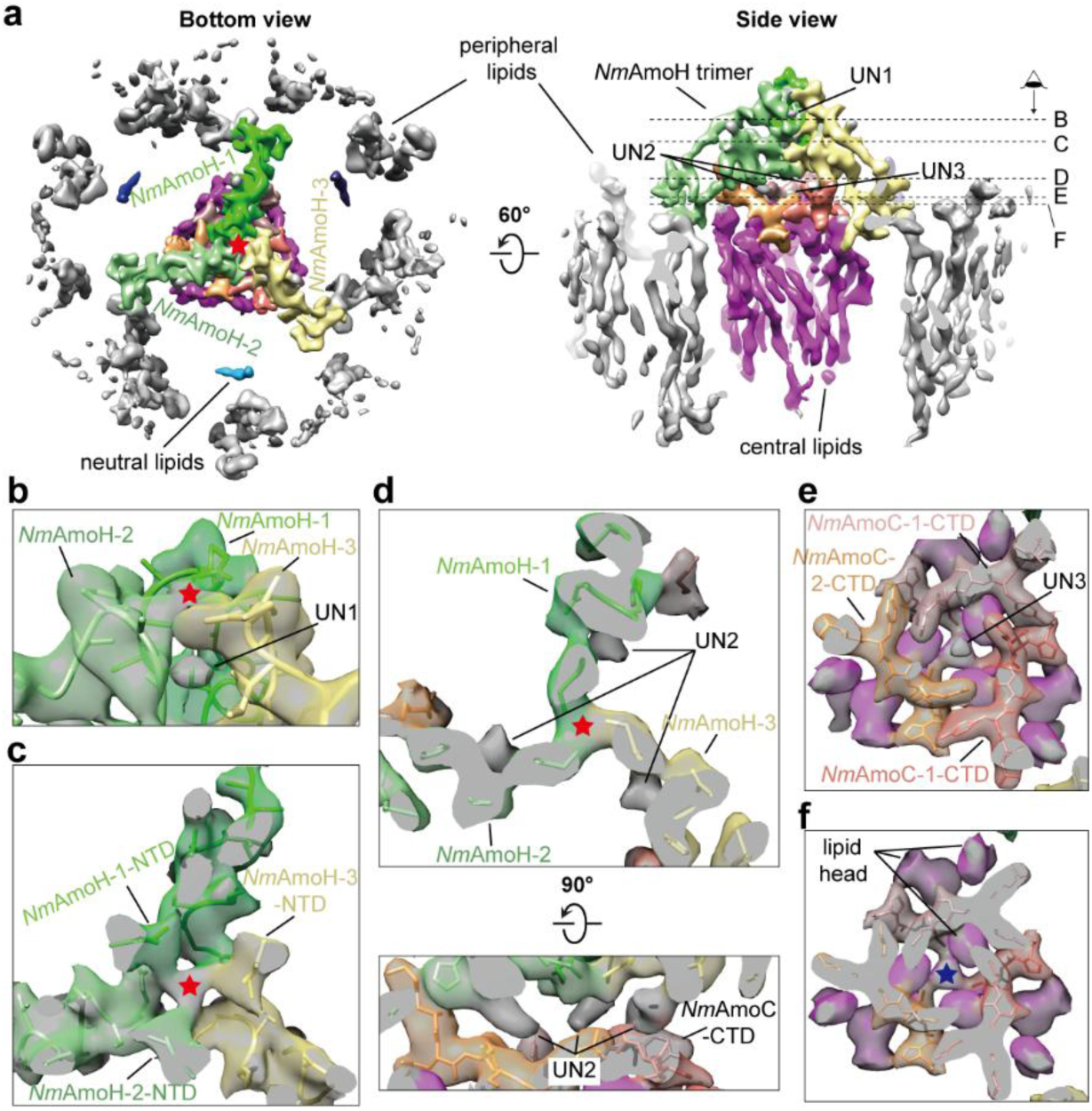
Three-layered architecture of *Nm*AmoH. **a.** Overall structure of the *Nm*AmoH homotrimer and associated archaeal lipid components. Archaeal lipids are categorized into three regions: peripheral (grey), central (purple), and neutral (cyan). The trimeric *Nm*AmoH adopted a propeller-shaped conformation and the three subunits are color-coded. Peripheral regions of the three *Nm*AmoH subunits interacted with peripheral lipids, while the trimeric *Nm*AmoH was positioned beneath the central lipids and interacted with the C-terminus of *Nm*AmoC. Bottom (left panel) and side (right panel) views are shown. Sliced layer views are presented in **(b– f)**. Three unassigned spherical densities are labeled UN1, UN2, and UN3. **b-d.** First (**b**), second (**c**), and third (**d**) layered structures of the trimeric *Nm*AmoH. Red stars indicate the center of each layer, with corresponding unknown densities annotated. **e-f.** Views of two sliced layers near the base of the central lipids. UN3 and the polar heads of the central lipids are indicated. The blue star in (**f**) marks the center at the base of the central lipids.

**Extended Data Figure 6.**
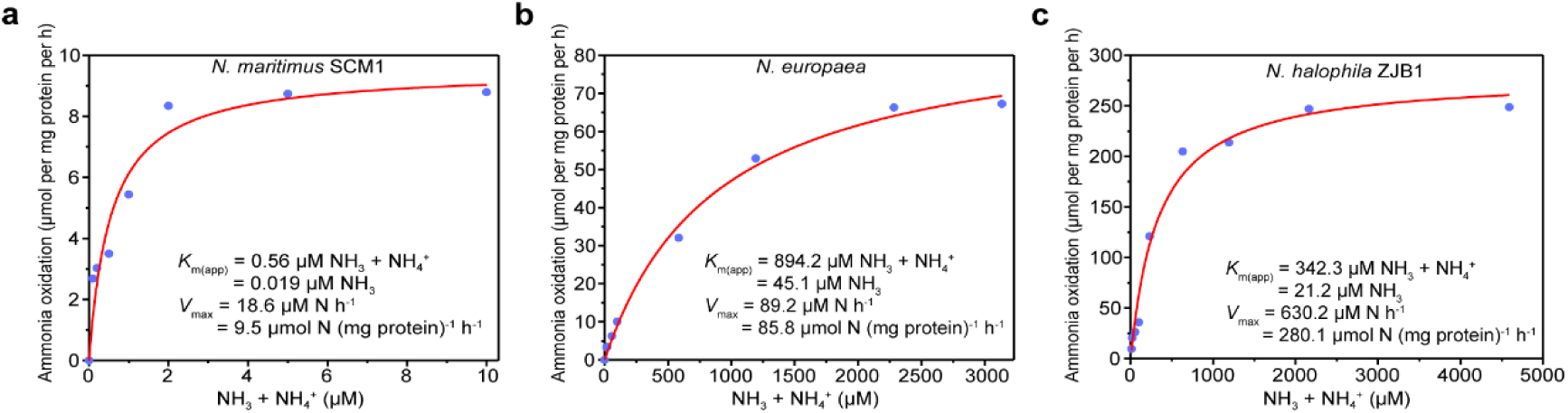
Ammonia oxidation kinetics of representative AOA and AOB. Michaelis–Menten kinetic analysis of ammonia oxidation by *N. maritimus* SCM1 (**a**), *N. europaea* (**b**), and *N. halophila* ZJB1 (**c**). Initial ammonia oxidation rates were measured over increasing NH_3_ + NH_4_^+^ concentrations and fitted by nonlinear regression. The apparent *K*_m(app)_ and *V*_max_ values indicate higher substrate affinity but lower maximum activity in AOA compared with AOB.

**Extended Data Figure 7.**
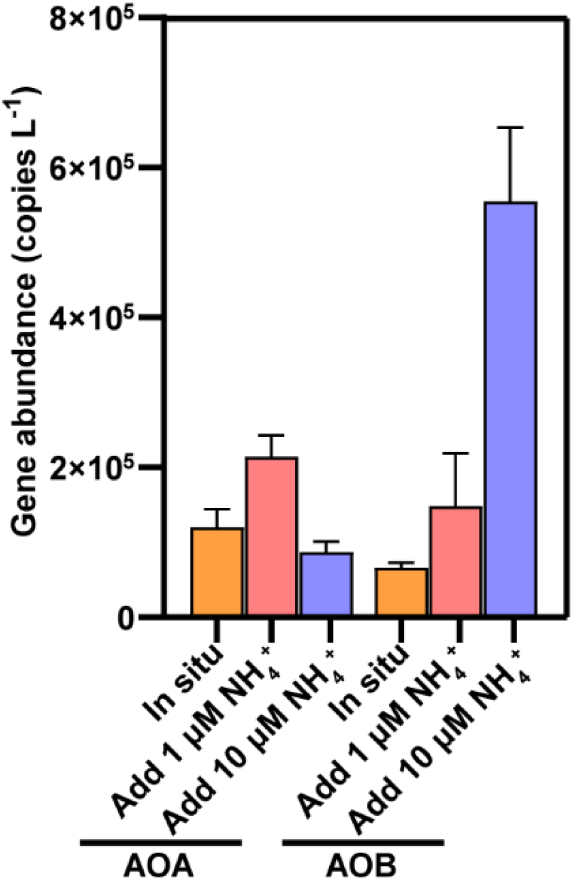
Responses of AOA and AOB to different ammonium regimes. Gene abundance of AOA and AOB (indicated by *amoA* gene) in natural seawater incubations under in situ conditions or after enrichment with 1 μM or 10 μM NH_4_^+^. Low ammonium addition preferentially promotes AOA, whereas higher ammonium addition more strongly increases AOB abundance, consistent with differential substrate adaptation of AOA and AOB.

**Extended Data Figure 8.**
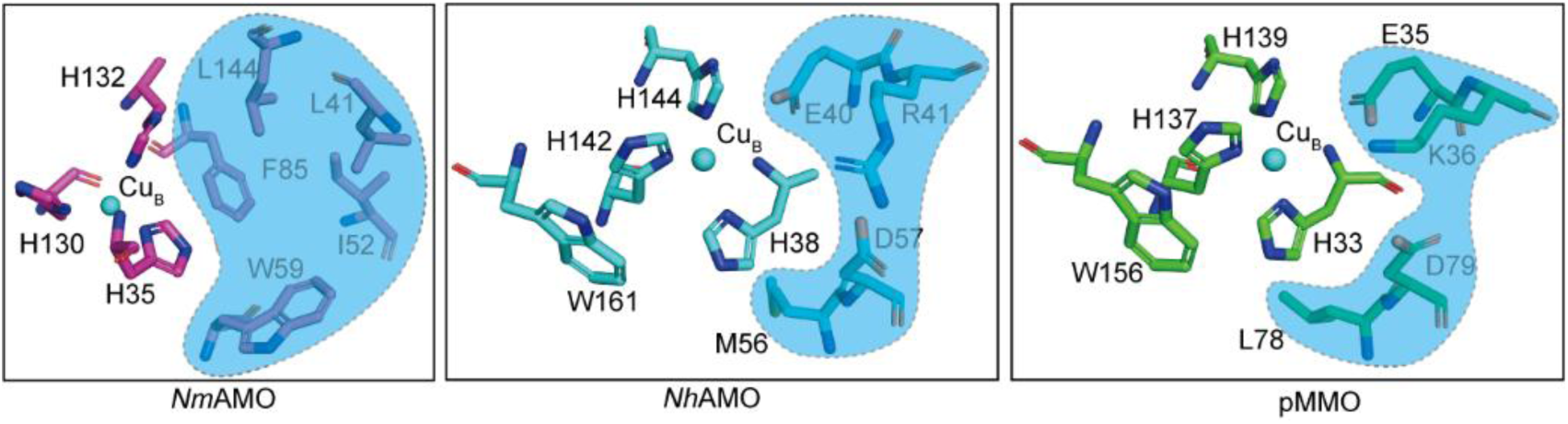
Comparison of the Cu_B_-binding sites in *Nm*AMO, *Nh*AMO and pMMO. The blue shaded regions highlight the distinct coordination environments among the three enzymes.

**Extended Data Figure 9.**
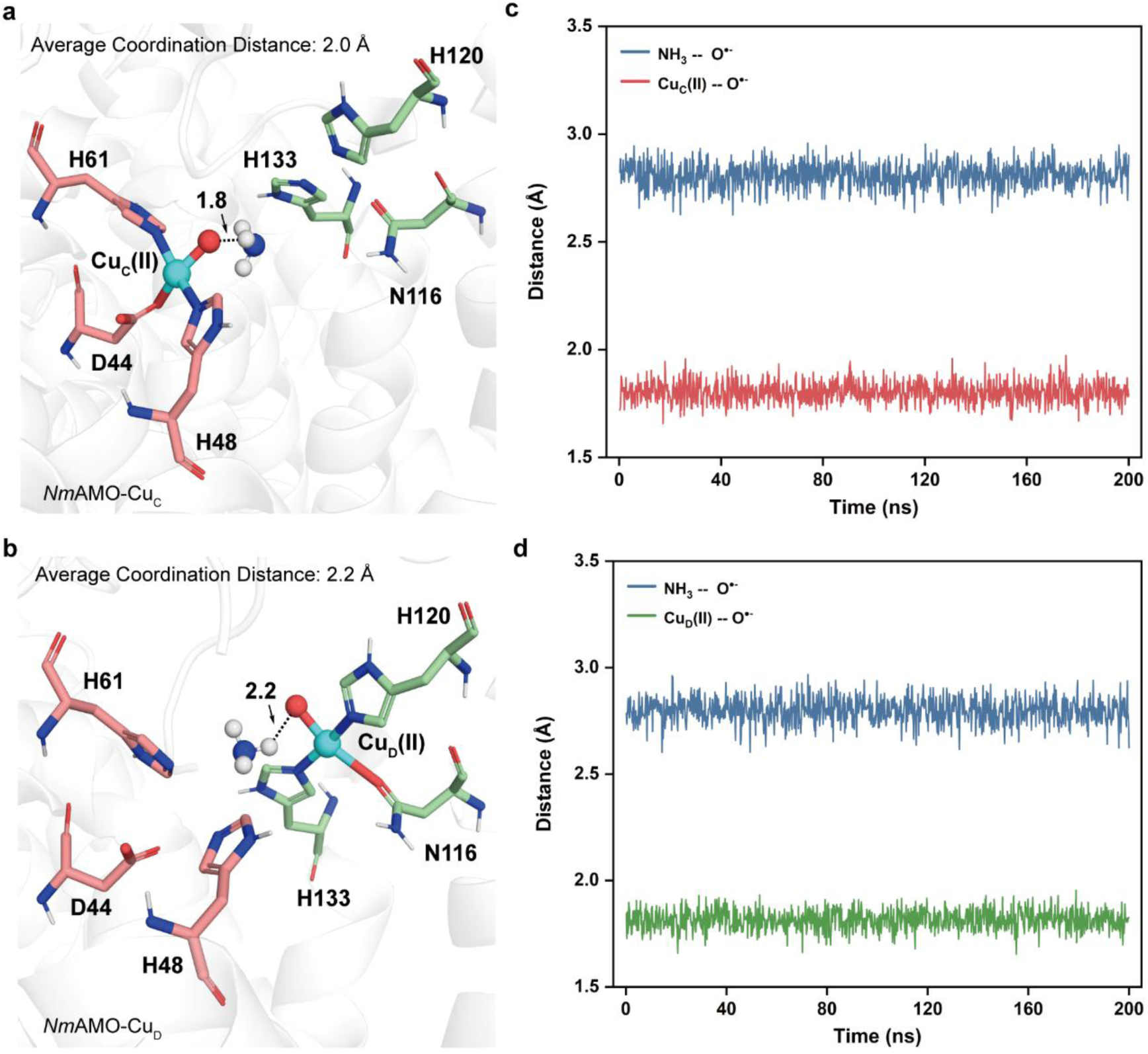
The binding modes between potential *Nm*AMO reactive oxygen species and the substrate ammonia. **a-b.** MD-averaged binding conformations of ammonia with the mononuclear reactive oxygen species Cu_C_(II)−O^•–^ (**a**) and Cu_D_(II)−O^•–^ (**b**). **c-d.** Time-dependent changes in key coordination distances and in the distances between ammonia and the mononuclear copper species Cu_C_(II)– O^•–^ (**c**) and Cu_D_(II)–O^•–^ (**d**).

**Extended Data Figure 10.**
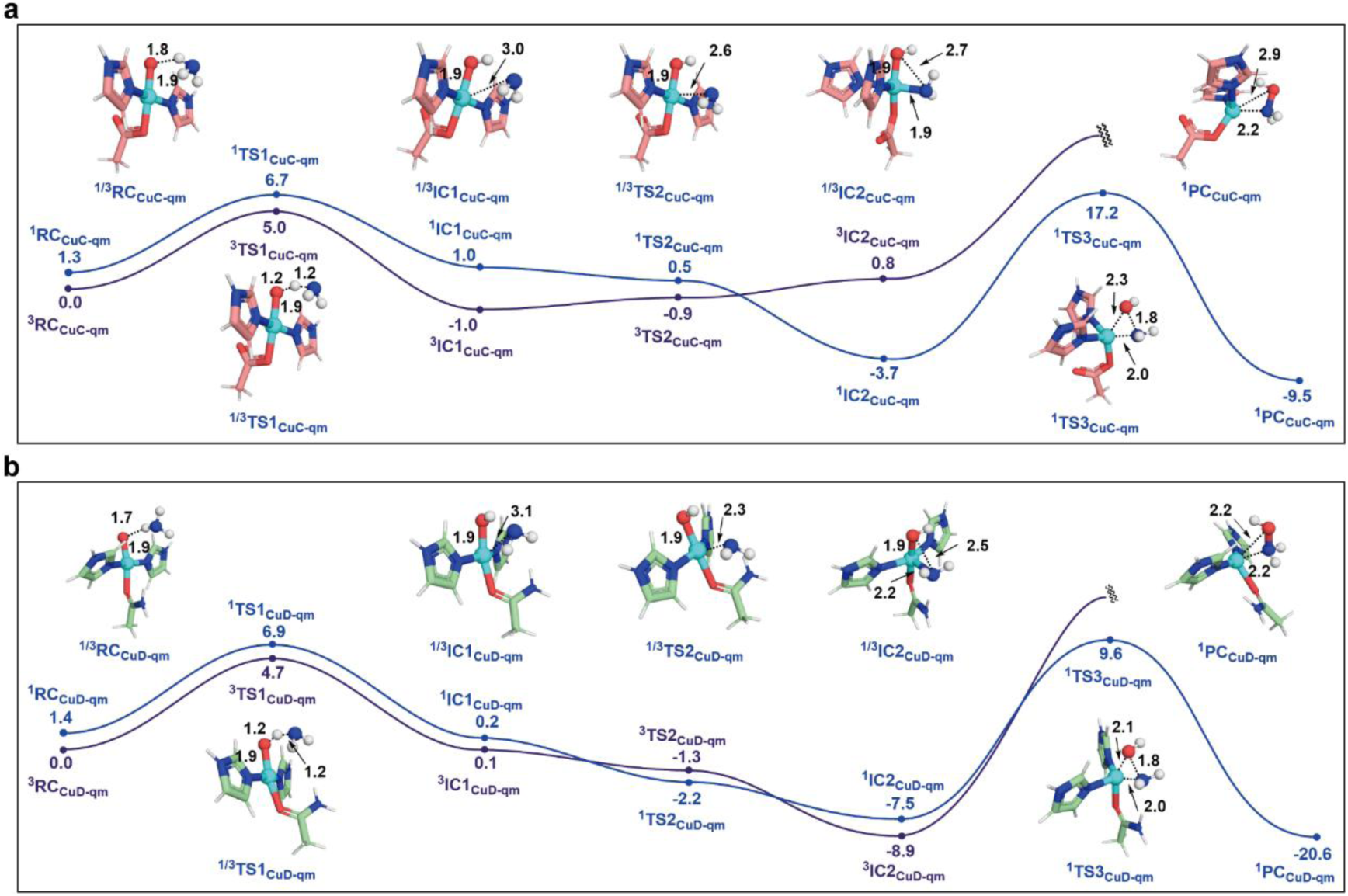
Comparison of the catalysis of potential *Nm*AMO reactive oxygen species. QM (UB3LYP/def2-TZVP//def2-SVP) calculated potential energy profile (in kcal/mol) for Cu_C_(II)−O^•–^ (**a**) and Cu_D_(II)−O^•–^ (**b**) mediated ammonia hydroxylation to hydroxylamine in both open-shell singlet and triplet states. Key distances are given in Å. Starting from the Cu_C_(II)–O^•^⁻ oxygen species (^3^RC_CuC_), HAT from NH_3_ proceeds with a low energy barrier of 5.0 kcal·mol⁻¹ (^3^RC_CuC-qm_ → ^3^TS1_CuC-qm_), yielding a Cu_C_(II)–OH⁻ and NH_2_^•^ intermediate (^3^IC1_CuC-qm_). The subsequent NH_2_^•^ rebound to the Cu_C_(II) site occurs with a small barrier of ∼ 0.5 kcal/mol, generating the more stable open-shell singlet intermediate (^1^IC2_CuC-qm_), which protects the NH_2_^•^ radical. The ensuing coupling between the hydroxyl and amino groups requires an activation barrier of 20.9 kcal·mol⁻¹ (^1^IC2_CuC-qm_ → ^1^TS3_CuC-qm_) to form NH_2_OH, releasing 9.5 kcal·mol⁻¹ (^1^PC_CuC-qm_). Similarly, when initiated from Cu_D_(II)–O^•^⁻ (^3^RC_CuD-qm_), hydrogen abstraction proceeds via ^3^RC_CuD-qm_ to ^1^TS1_CuD-qm_ with a barrier of 4.7 kcal·mol⁻¹, forming the Cu_D_(II)–OH⁻ and NH_2_^•^ (^1^IC1_CuD-qm_). The NH_2_^•^ rebound step is endothermic, followed by hydroxyl–amino coupling (^1^IC1_CuD-qm_ → ^1^TS3_CuD-qm_) with a barrier of 18.5 kcal·mol⁻¹ to yield NH_2_OH, releasing 20.6 kcal·mol⁻¹ (^1^PC_CuD-qm_).

**Extended Data Figure 11.**
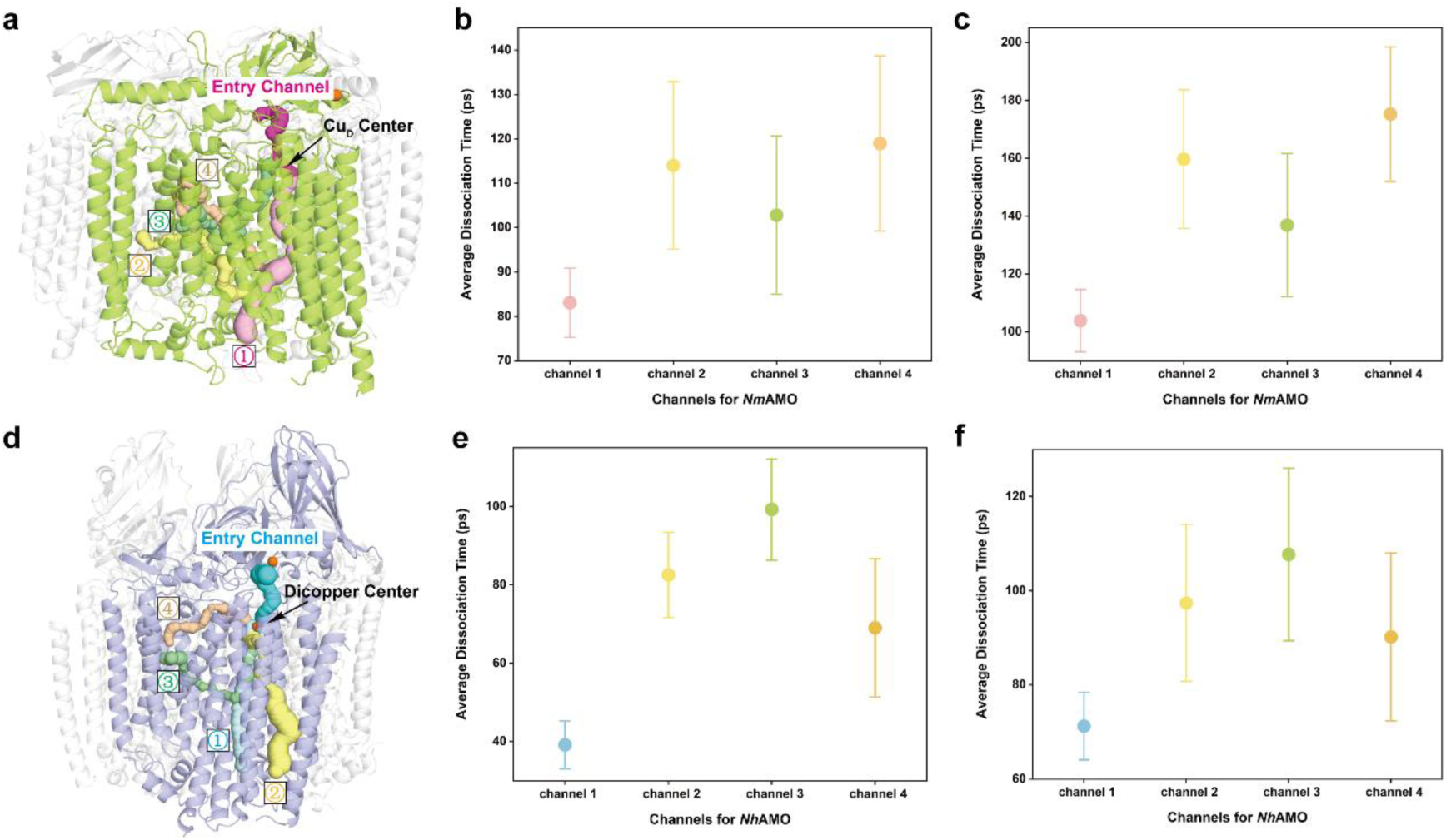
Evaluation of the channels for inhibitors dissociation from *Nm*AMO and *Nh*AMO. **a.** Overview of the entry channel (highlighted in magenta) and all potential dissociation channels in *Nm*AMO. **b-c.** RAMD-calculated dissociation times of DMP (**b**) and ATU (**c**) exiting from *Nm*AMO. **d.** Overview of the entry channel (highlighted in cyan) and all potential dissociation channels in *Nh*AMO. **e-f.** RAMD-calculated dissociation times of DMP (**e**) and ATU (**f**) exiting from *Nh*AMO. Comparison of dissociation times shows that ATU dissociates more slowly than DMP in both *Nm*AMO and *Nh*AMO, and that DMP (39 ps vs. 83 ps) and ATU (71 ps vs. 103 ps) dissociating more readily from *Nh*AMO than from *Nm*AMO.

**Extended Data Figure 12.**
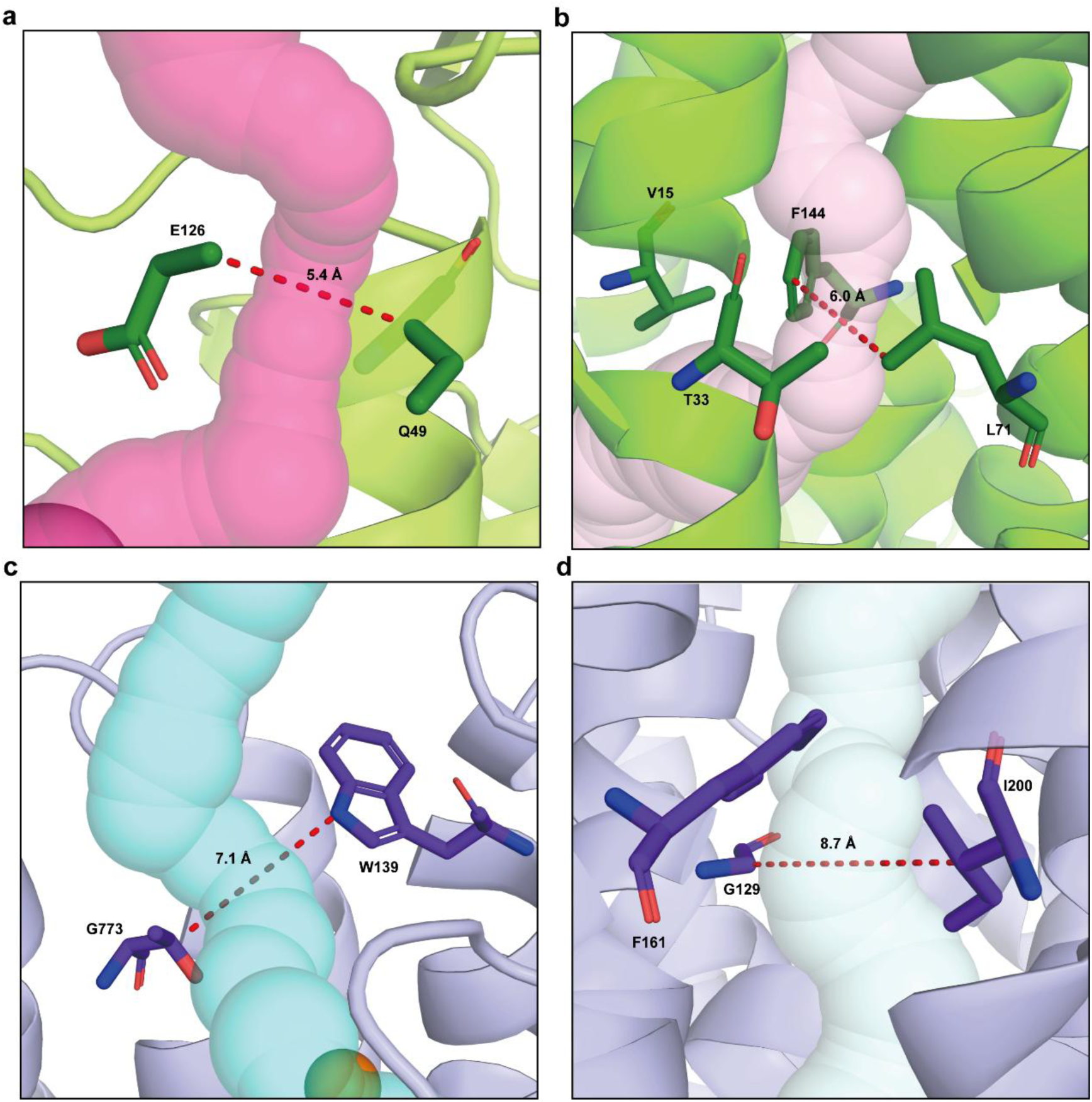
Comparison of the narrowest distance along the inhibitor entry and dissociation channels in *Nm*AMO and *Nh*AMO. **a-b.** Channels for exogenous inhibitors entering (**a**) and dissociating (**b**) from the Cu_D_ site of *Nm*AMO. **c-d.** Channels for exogenous inhibitors entering (**c**) and dissociating (**d**) from the dicopper center of *Nh*AMO. The amino acid residues located at the narrowest region of the *Nm*AMO and *Nh*AMO channel are depicted as green and purple stick models, respectively. Comparison of the entry channels of *Nm*AMO and *Nh*AMO shows that *Nm*AMO exhibits a narrower channel radius than *Nh*AMO, indicating reduced accessibility of exogenous inhibitors to the Cu_D_ sites of *Nm*AMO.

**Extended Data Figure 13.**
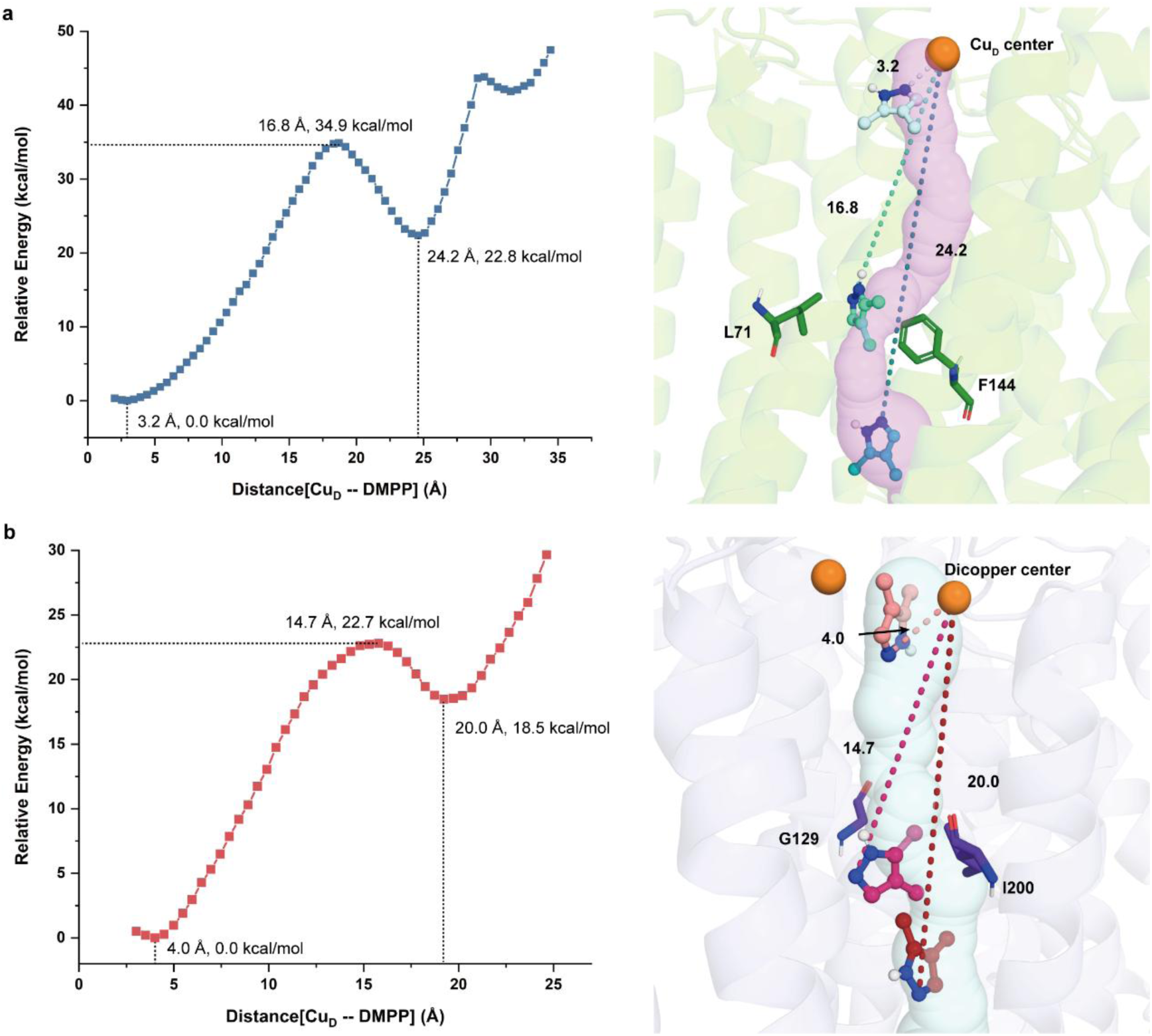
Evaluation of the mechanism of DMP dissociation from the active sites of *Nm*AMO and *Nh*AMO. **a.** Umbrella sampling calculated free energy profiles (kcal/mol) for the exit of DMPP from the Cu_D_ center to the outside via the dissociation channel of *Nm*AMO. **b.** Umbrella sampling calculated free energy profiles (kcal/mol) for the exit of DMP from the dicopper center to the outside via the dissociation channel of *Nh*AMO. The reaction coordinate is defined as the distance between the Cu_D_ sites and the nitrogen atom of inhibitor DMP. The initial conformation of DMP and its conformations during the dissociation process are represented by ball-and-stick models. The amino acid residues at the narrowest regions of the *Nm*AMO and *Nh*AMO channels are displayed as green and purple stick models, respectively. Key distances are given in Å.

**Extended Data Figure 14.**
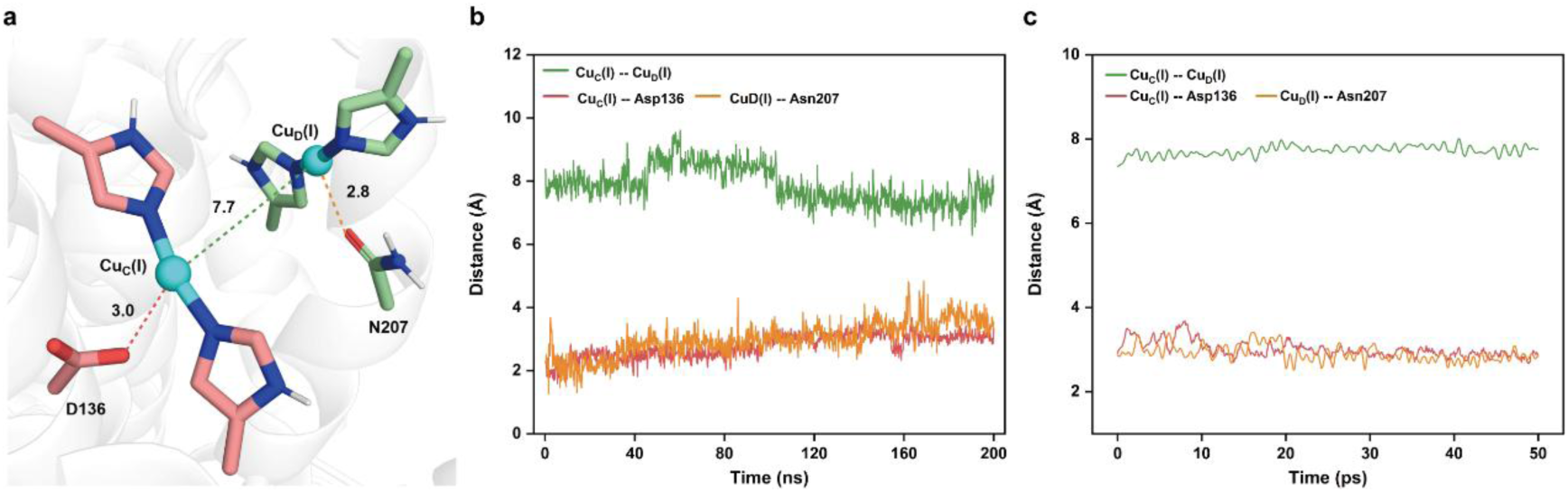
The coordination geometries of the dinuclear copper active site in *Nh*AMO. **a.** Representative geometry of the dinuclear copper active site from QM/MM MD simulation. Key distances are given in Å. **b-c.** Time-dependent changes in the distances between dinuclear copper center and their surrounding key amino acid residues during the MD (**b**) and QM/MM MD simulations (**c**). Analysis of representative conformations and coordination environments revealed that, in the absence of O_2_ or reducing cofactors, the dinuclear copper center in NhAMO is stably maintained in the monovalent Cu_C_(I) and Cu_D_(I).

**Extended Data Figure 15.**
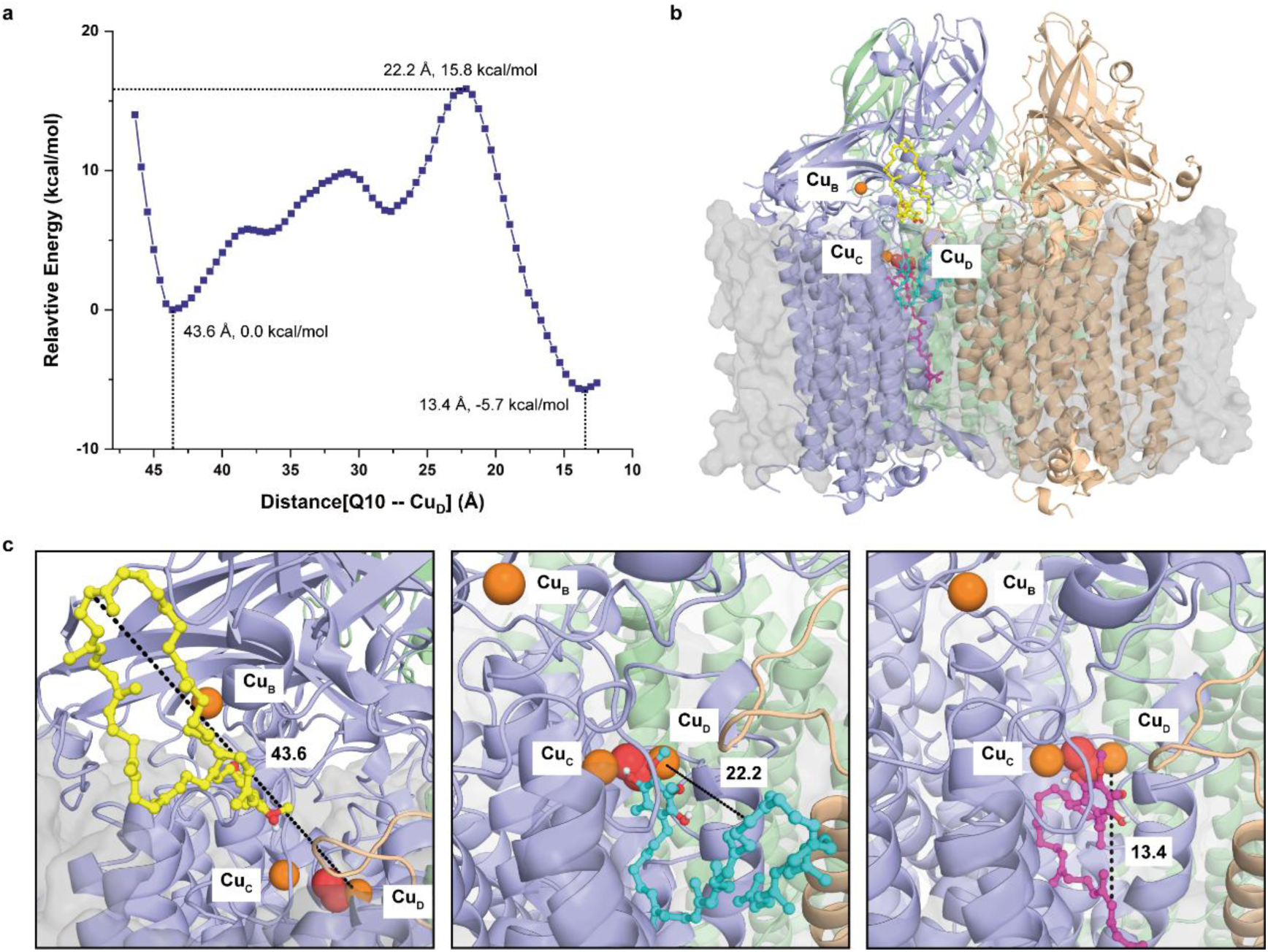
Evaluation of the mechanism of reductant CoQ10H_2_ entry into the active site of *Nh*AMO. **a.** Umbrella sampling calculated free energy profiles (kcal/mol) for the CoQ10H_2_ entry from the extracellular side to the dicopper center of *Nh*AMO. **b-c.** Representative conformations of CoQ10H_2_ along its translocation pathway toward the dicopper center of *Nh*AMO. The reaction coordinate is defined as the distance between the Cu_D_ site and the central carbon atom of CoQ10H_2_. The *Nh*AMO is shown as a cartoon, with the Cu centers depicted as orange spheres and the phospholipid bilayer displayed as a gray surface. The initial conformation of CoQ10H_2_ and two representative intermediates along the entry pathway are shown as yellow, cyan, and magenta ball-and-stick models, respectively. Key distances are given in Å.

**Extended Data Figure 16.**
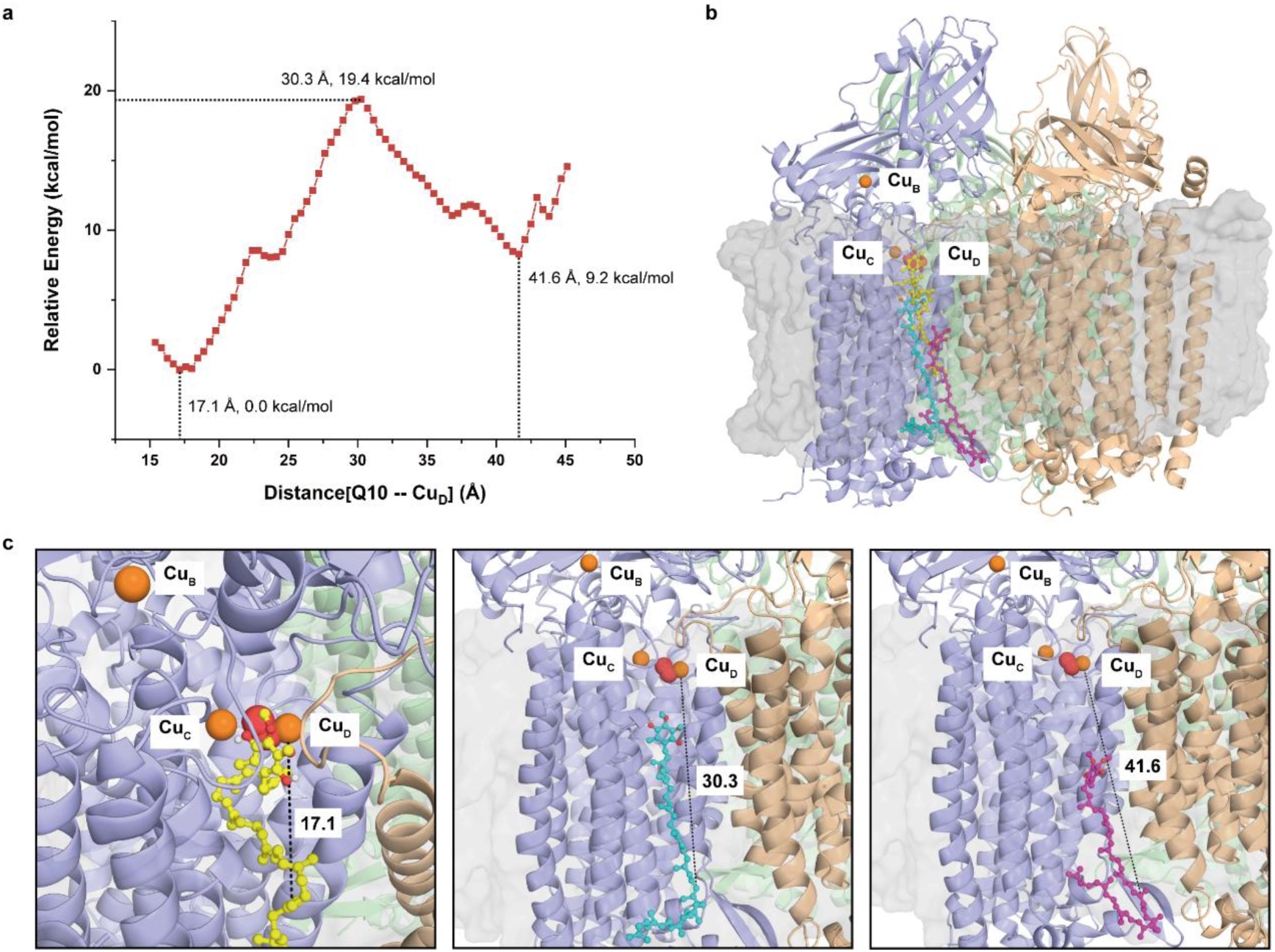
Evaluation of the mechanism of reductant CoQ10H_2_ dissociation from the dicopper center of *Nh*AMO. **a.** Umbrella sampling calculated free energy profiles (kcal/mol) for the exit of CoQ10H_2_ from the dicopper center of *Nh*AMO to the outside. **b-c.** Representative conformations of CoQ10H_2_ along its dissociation pathway from the dicopper center of *Nh*AMO. The reaction coordinate was defined as the distance between the CuD site and the central carbon atom of CoQ10H_2_. The *Nh*AMO is shown as a cartoon, with the Cu centers depicted as orange spheres and the phospholipid bilayer displayed as a gray surface. The initial conformation of CoQ10H_2_ and two representative intermediates along the dissociation pathway are shown as yellow, cyan, and magenta ball-and-stick models, respectively. Key distances are given in Å.

**Extended Data Figure 17.**
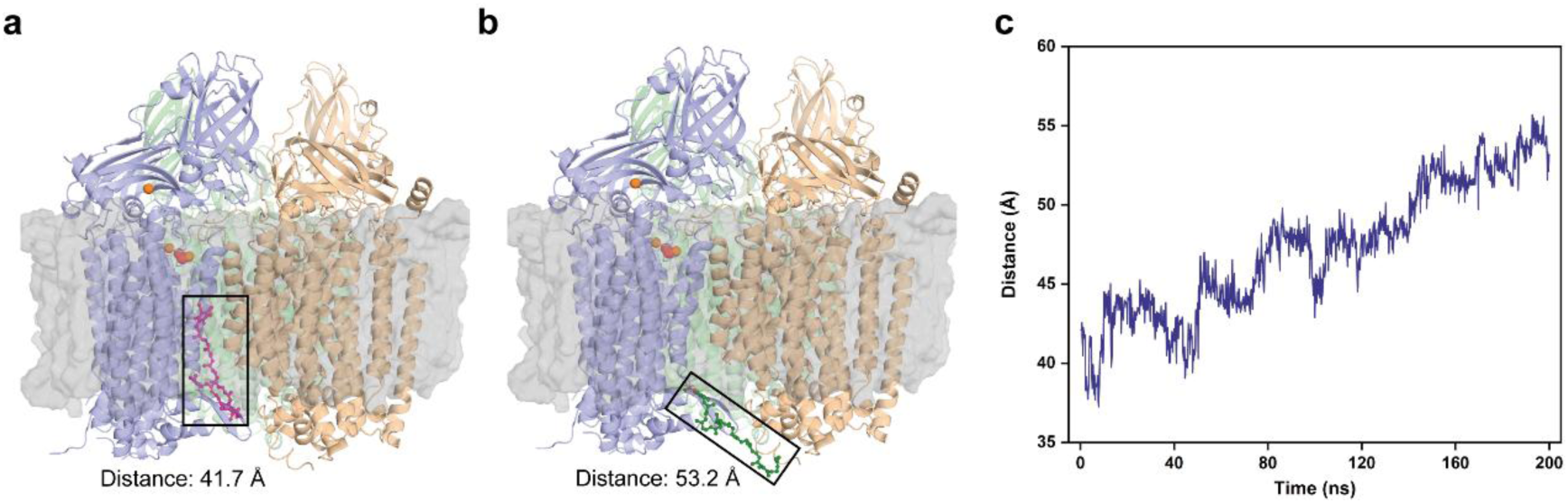
The process of the reductant CoQ10H_2_ dissociation from the dicopper center of *Nh*AMO. **a.** The initial conformation was selected from umbrella sampling, whereby the distance between the Cu_D_ site and the central carbon atom of CoQ10H_2_ was 41.7 Å. **b.** The representative dissociated conformation was selected during the 200 ns unbiased potential MD simulations, whereby the distance between the Cu_D_ site and the central carbon atom of CoQ10H_2_ was 53.2 Å. **c.** Time-dependent changes in the distances between the Cu_D_ site and the central carbon atom of CoQ10H_2_ during the 200 ns unbiased potential MD simulations. Ultimately, CoQ10H_2_ can be successfully released from the dicopper centers of *Nh*AMO into the external environment.

**Extended Data Figure 18.**
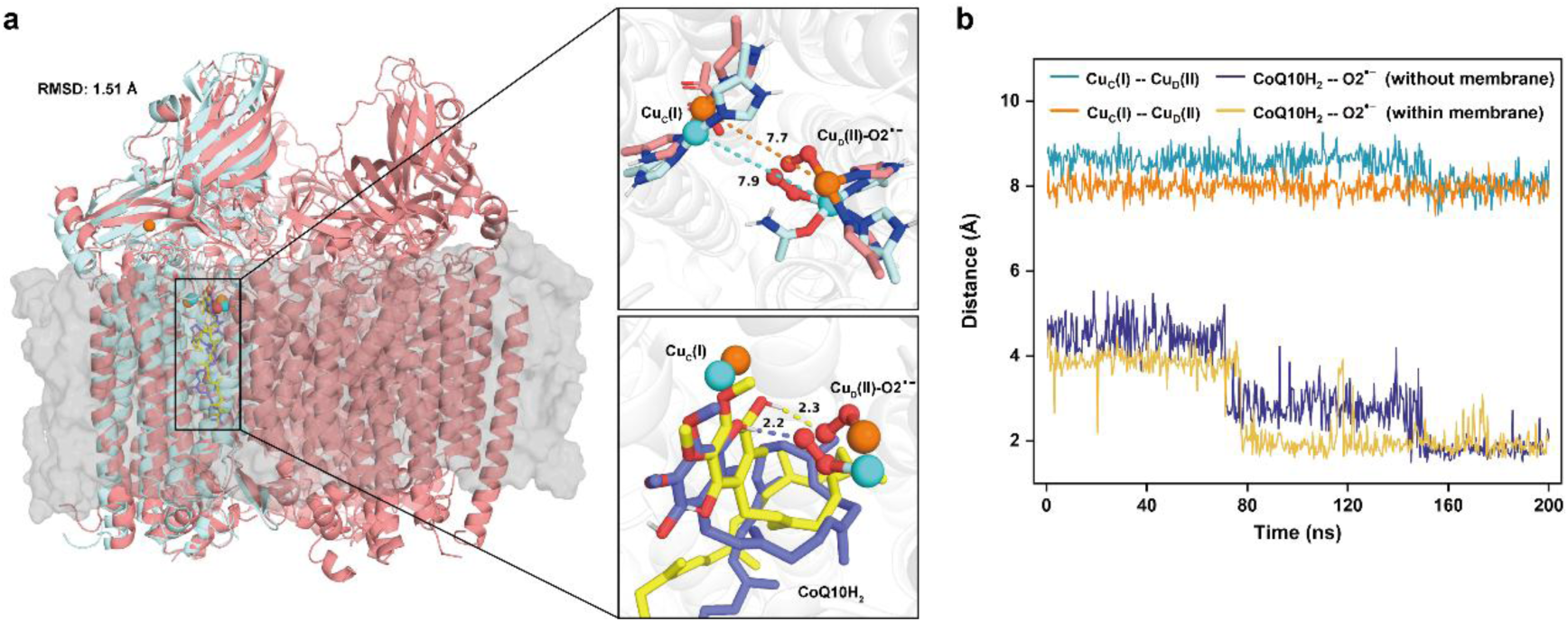
Comparison of the binding modes of CoQ10H_2_ in membrane-bound and membrane-free *Nh*AMO. **a.** Structural superimposition of the overall three-dimensional structures of membrane-bound and membrane-free *Nh*AMO, highlighting the coordination environment of the dicopper center and the binding conformation of CoQ10H_2_. The membrane-bound and membrane-free *Nh*AMO structures are shown as red and cyan cartoons, respectively, with the phospholipid bilayer represented as a gray surface. The dicopper center and its coordinating residues in membrane-bound and membrane-free *Nh*AMO are displayed as cyan and orange spheres and ball-and-stick models, respectively. CoQ10H_2_ in the two structures is depicted as yellow and blue ball-and-stick models, respectively. **b.** Time-dependent changes in the distances between the dinuclear copper sites, as well as between CoQ10H_2_ and the key hydrogen-transfer acceptor Cu_D_(II)−O_2_^•–^, in membrane-bound and membrane-free *Nh*AMO during the MD simulations.

**Extended Data Figure 19.**
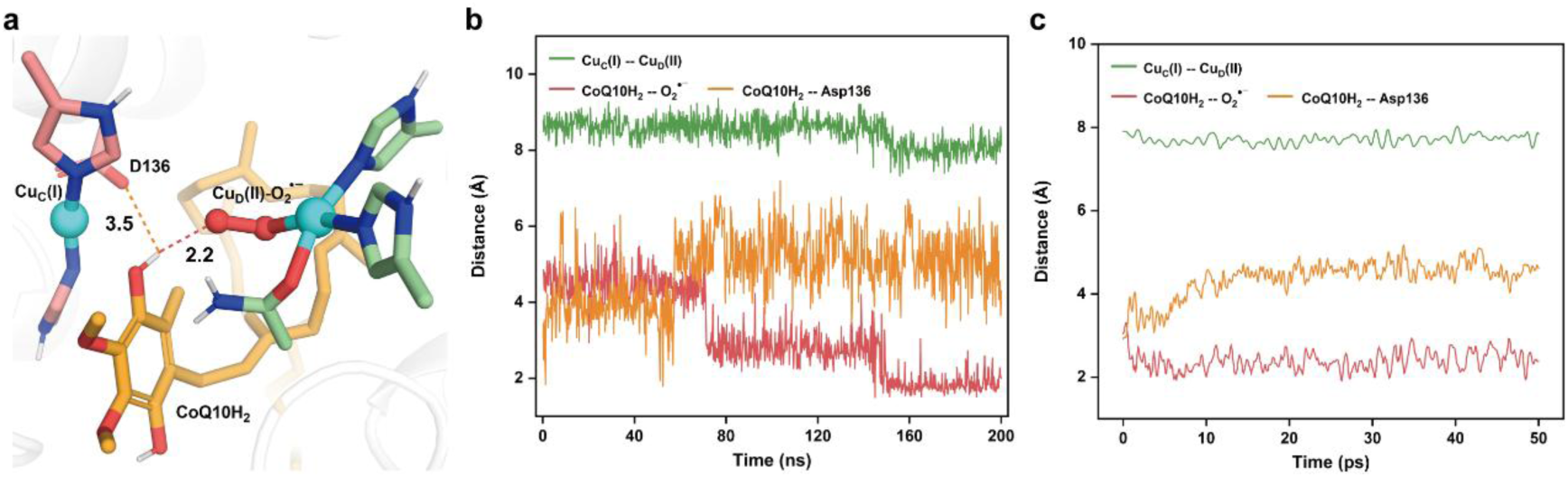
The binding mode of coenzyme Q10 (CoQ10H_2_) within the dinuclear copper active site. **a.** Representative geometry of CoQ10H_2_ within the dinuclear copper active site from QM/MM MD simulation. Key distances are given in Å. **b-c.** Time-dependent changes in the distances between dinuclear copper sites and in the distances between CoQ10H_2_ and its surrounding key proton transfer acceptors (Cu_D_(II)−O_2_^•–^ and Asp136) during the MD (**b**) and QM/MM MD simulations (**c**). MD and QM/MM-MD simulations confirmed that CoQ_10_H_2_ can stably bind within the substrate-accessible pocket, with its hydroxyl group oriented toward Cu_D_(II)–O_2_^•⁻^ through H-bond interaction at a distance of ∼ 2.2 Å. Analysis of representative conformations and distance fluctuations indicates that, upon entry into the binuclear copper center, the phenolic proton of CoQ10H_2_ preferentially transfers to the O_2_ ^•–^ coordinated to the Cu_D_(II) site rather than to Asp136.

**Extended Data Figure 20.**
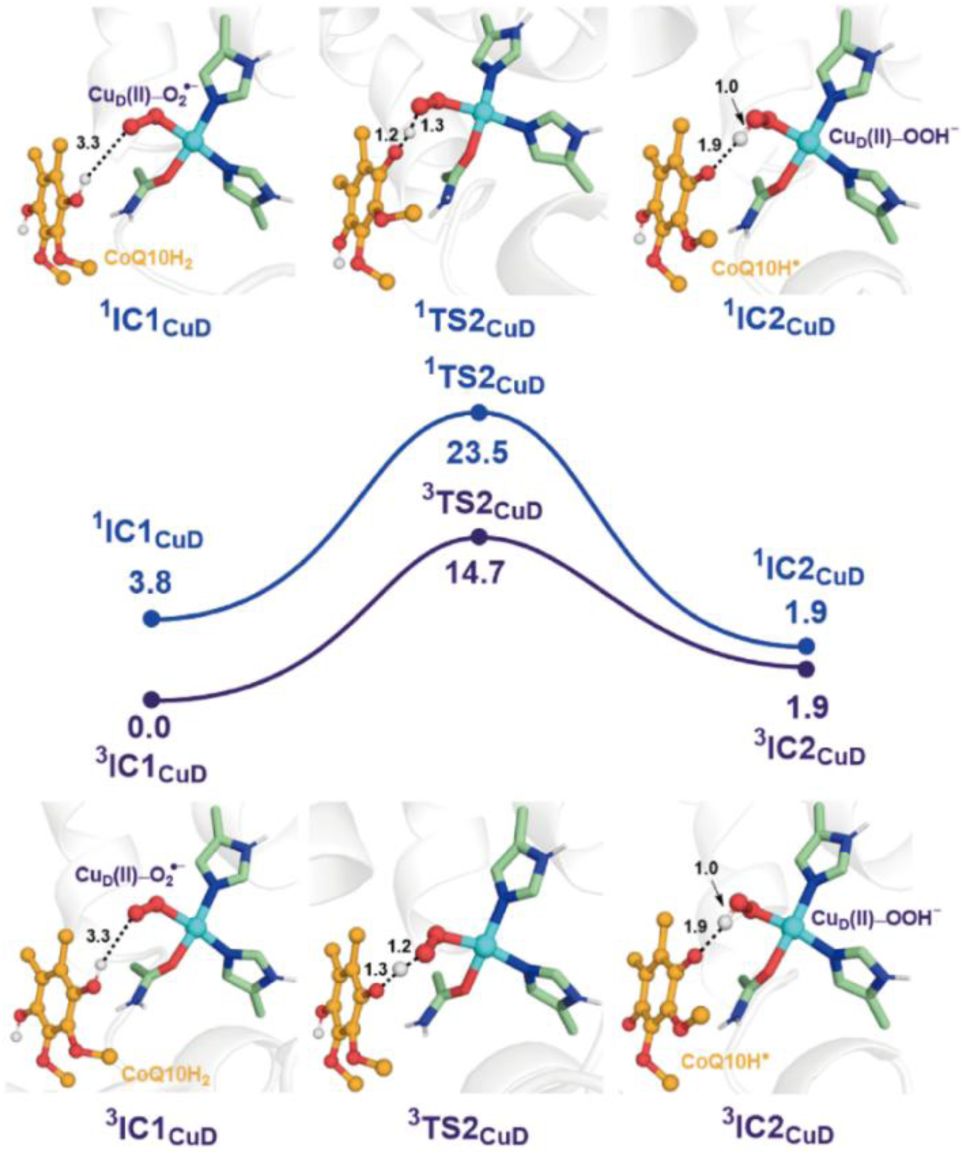
Calculated mechanisms of oxygen activation and subsequent formation of Cu_D_(II)-OOH^−^ species. QM (UMN15-D3/def2-TZVP)/MM relative energies (kcal/mol) for Cu_D_(II)−OOH^−^ (IC2_CuD_) formation from the Cu_D_(II)−O_2_^•−^ and CoQ10H_2_ complex (IC1_CuD_) via the HAT processes in the open-shell broken-symmetry singlet and triplet states. QM(UMN15-D3/def2-SVP)/MM optimized geometries of key species involved in the reaction are presented. Key distances are given in Å. From this preorganized configuration of IC1_CuD_, HAT proceeds with an energy barrier of 14.7 kcal mol⁻¹ (^3^IC1_CuD_ → ^3^TS2_CuD_), yielding a Cu_D_(II)–OOH⁻ species that is slightly endergonic (1.9 kcal mol⁻¹). Notably, this barrier matches that reported for the reduction of Cu_D_(II)–O₂^•⁻^ by duroquinol in pMMO, highlighting an intrinsic propensity of the Cu_D_-bound superoxo species to accept hydrogen (14.2 kcal mol⁻¹) relative to its Cu_C_ counterpart (18.4 kcal mol⁻¹, see ref. 42 in MS).

**Extended Data Figure 21.**
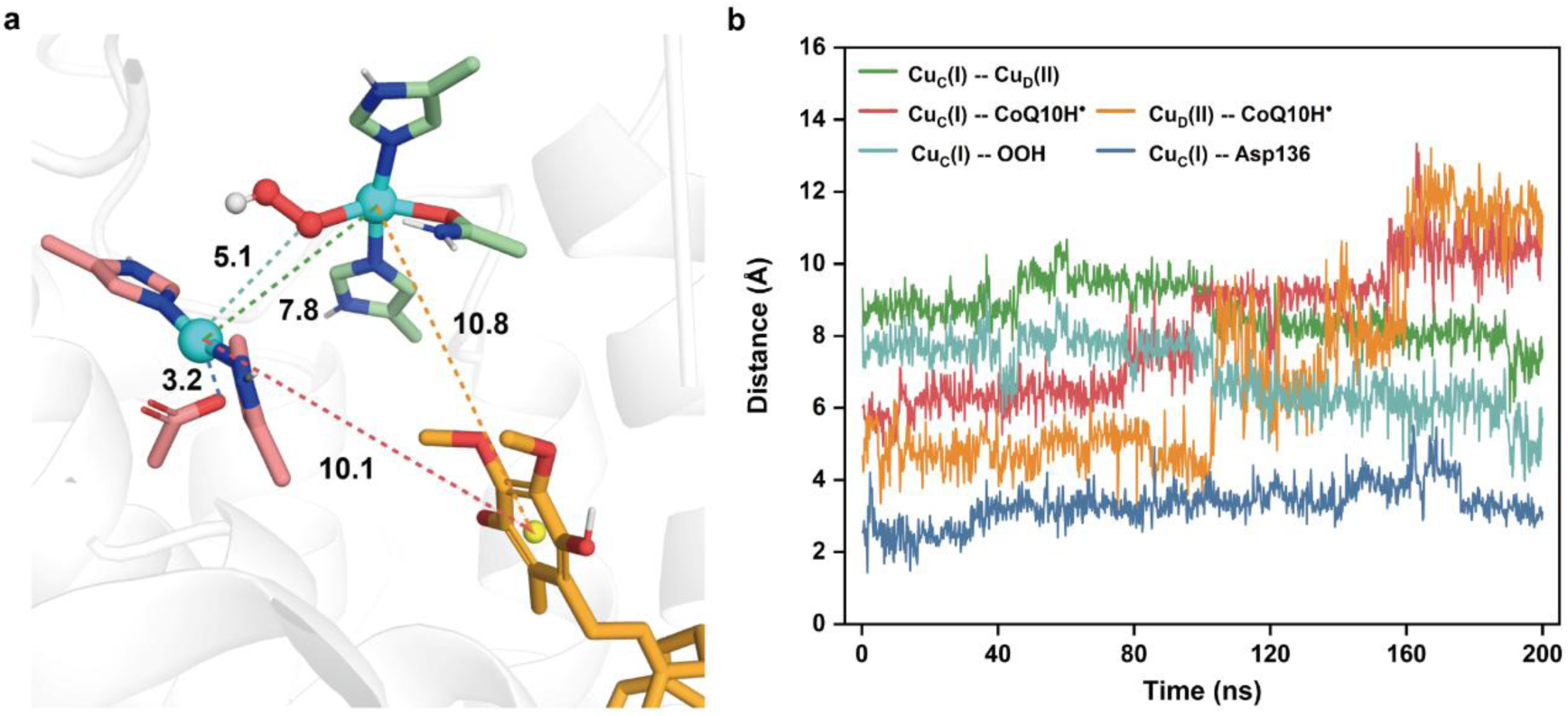
The stability of the CoQ10H^•^ radical binding within the dinuclear copper active site. **a.** Representative geometry of CoQ10H^•^ radical within the dinuclear copper active site from MD simulation. Key distances are given in Å. **b.** Time-dependent changes in the distances between dinuclear copper sites, the distances between CoQ10H^•^ radical and dinuclear copper sites, and the distances between Cu_C_(I) and the surrounding atoms during the MD simulations. Following hydrogen transfer from CoQ10H_2_ to form the oxidized CoQ10H^•^ radical, the CoQ10H^•^ radical moves slightly away from the binuclear copper center. This displacement alleviates steric hindrance between the Cu_C_(I) and Cu_D_(II) sites, thereby facilitating closer approach of the two copper ions.

**Extended Data Figure 22.**
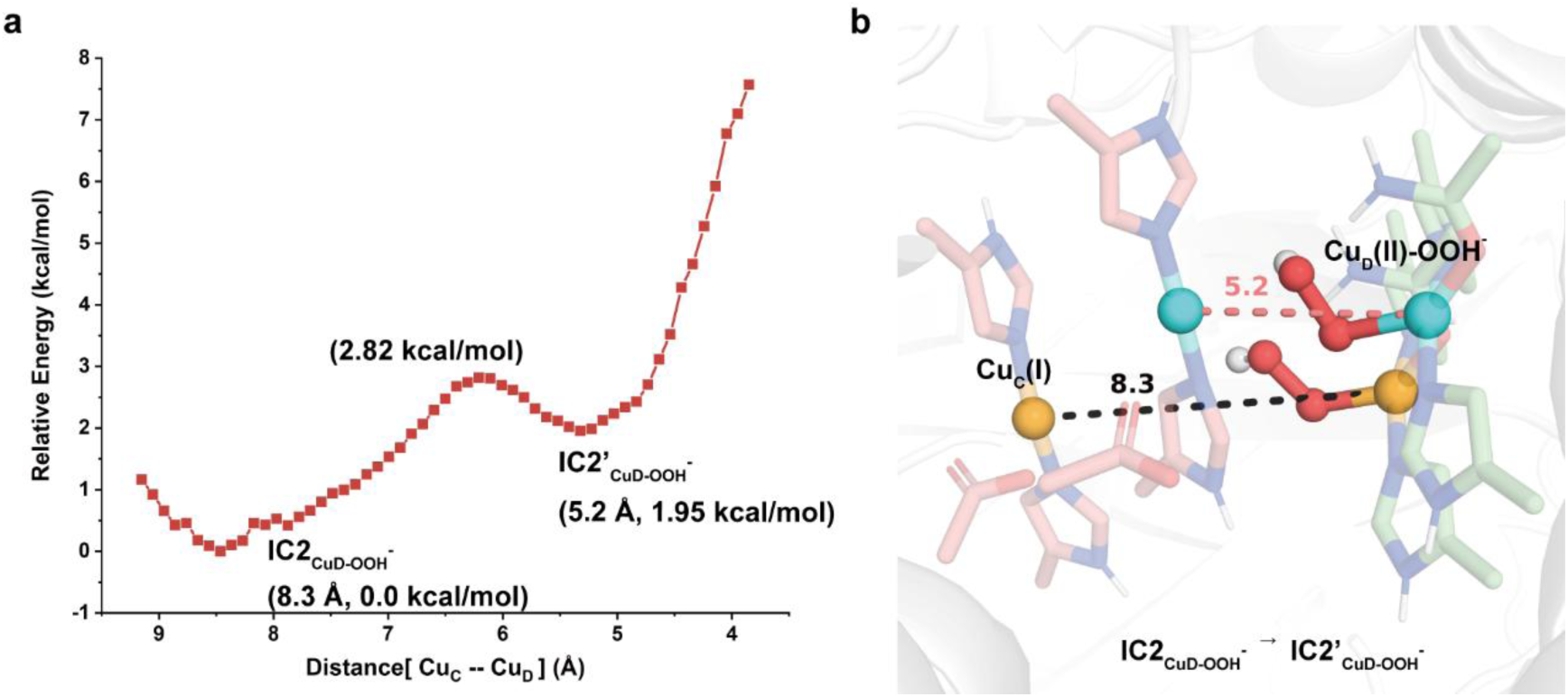
Free energy profile of Cu_C_(I) and Cu_D_(II)−OOH^−^ closure. **a.** Umbrella-sampling-calculated free energy profile (in kcal/mol) of the transition from the initial open state (IC2_CuD-OOH_^−^) to the closed state (IC2’_CuD-OOH_^−^) of the binuclear copper center. The reaction coordinate is defined as the distance between Cu_C_(I) and Cu_D_(II) sites. **b.** Representative conformations of the binuclear copper centre along the transition during the umbrella sampling, with the initial open state (IC2_CuD-OOH_^−^) and final closed state (IC2’_CuD-OOH_^−^) shown as orange and cyan sphere models, respectively. Key distances are given in Å. The approach of Cu_C_(I) and Cu_D_(II)−OOH^−^ proceeds over an energy barrier of 2.82 kcal/mol, corresponding to a reduction in the Cu_C_–Cu_D_ distance from 8.3 Å to 5.2 Å.

**Extended Data Figure 23.**
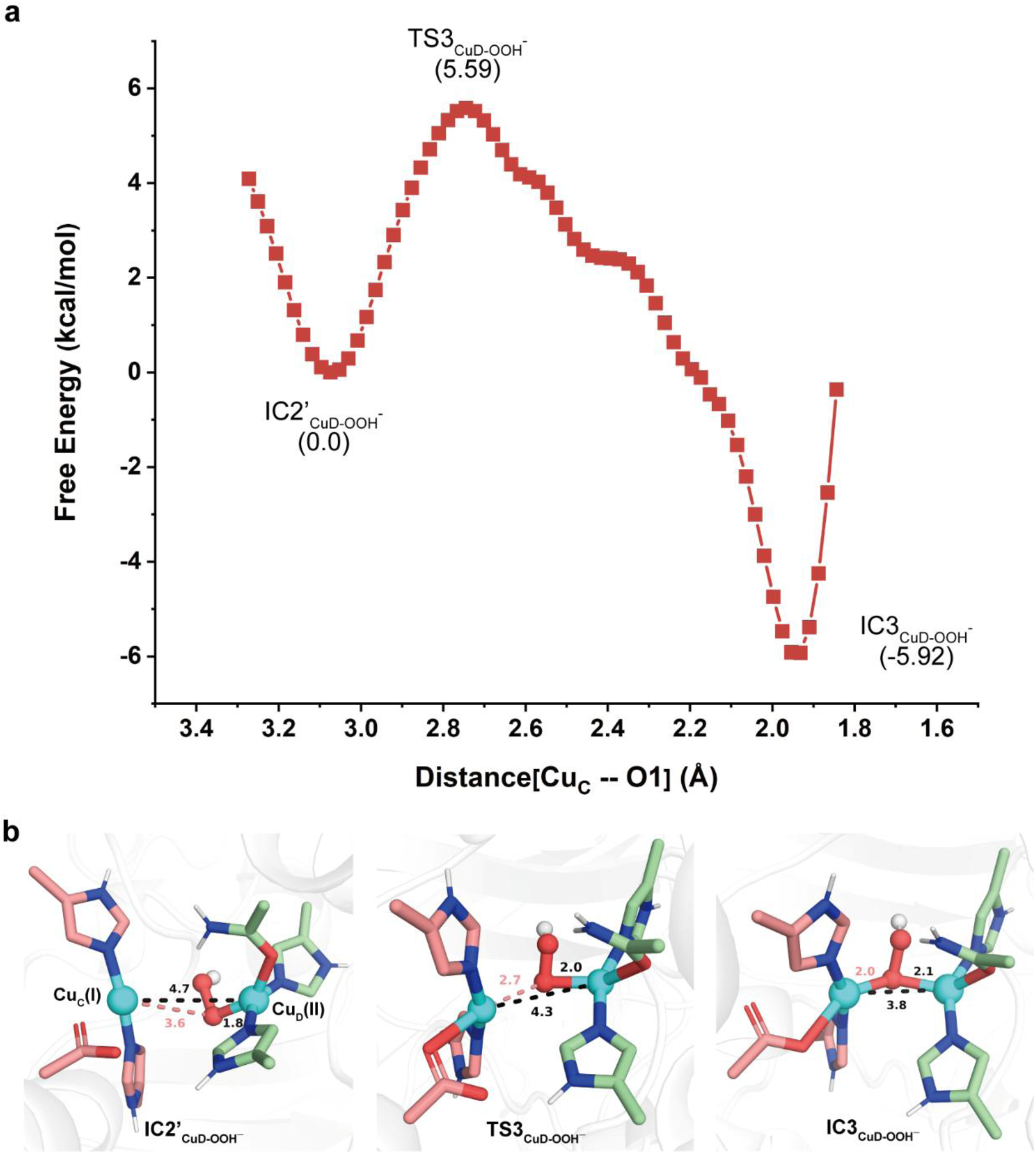
Calculated mechanisms of the (*μ*-hydroperoxo)Cu_C_(II)Cu_D_(I) intermediate formation. **a.** QM/MM metadynamics-calculated free energy profile (in kcal/mol) describing distal-oxygen coordination of Cu_D_–OOH⁻ to the Cu_C_ site, concomitant with electron transfer from Cu_C_(I) to Cu_D_(II) sites, culminating in the formation of the (*μ*-hydroperoxo)Cu_C_(II)Cu_D_(I) species (IC2’_CuD–OOH⁻_ → IC3_CuD–OOH⁻_). The reaction coordinate is defined as the distance between Cu_C_ site and the distal oxygen of Cu_D_(II)−OOH^−^. **b.** The geometry of intermediates and transition states, including the structures of IC2’_CuD−OOH_^−^, TS3_CuD−OOH_^−^ and IC3_CuD−OOH_^−^. Key distances are given in Å. Free energy calculations reveal that the (*μ*-hydroperoxo)Cu_C_(II)Cu_D_(I) species formation involves a substantial barrier of 5.59 kcal/mol and is thermodynamically favorable.

**Extended Data Figure 24.**
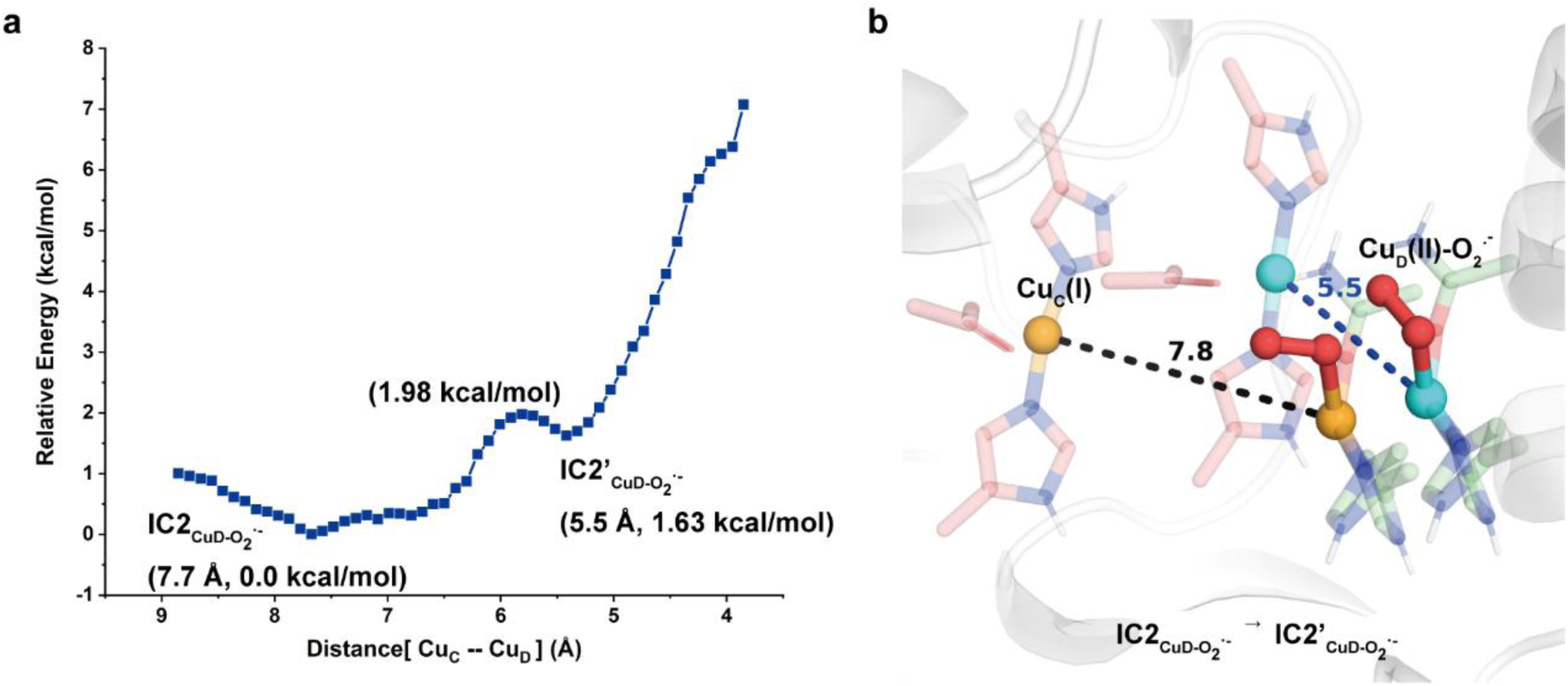
Free energy profile of Cu_C_(I) and Cu_D_(II)−O_2_^•−^ closure. **a.** Umbrella-sampling-calculated free energy profile (in kcal/mol) of the transition from the initial open state (IC2_CuD-O2_^•−^) to the closed state (IC2’_CuD-O2_^•−^) of the binuclear copper center. The reaction coordinate is defined as the distance between Cu_C_(I) and Cu_D_(II) sites. **b.** Representative conformations of the binuclear copper center along the transition during the umbrella sampling, with the initial open state (IC2_CuD-O2_^•−^) and final closed state (IC2’_CuD-O2_^•−^) shown as orange and cyan sphere models, respectively. Key distances are given in Å. The approach of Cu_C_(I) and CuD(II)−O2^•−^ proceeds over an energy barrier of 1.98 kcal/mol, corresponding to a reduction in the Cu_C_–Cu_D_ distance from 7.8 Å to 5.5 Å.

**Extended Data Figure 25.**
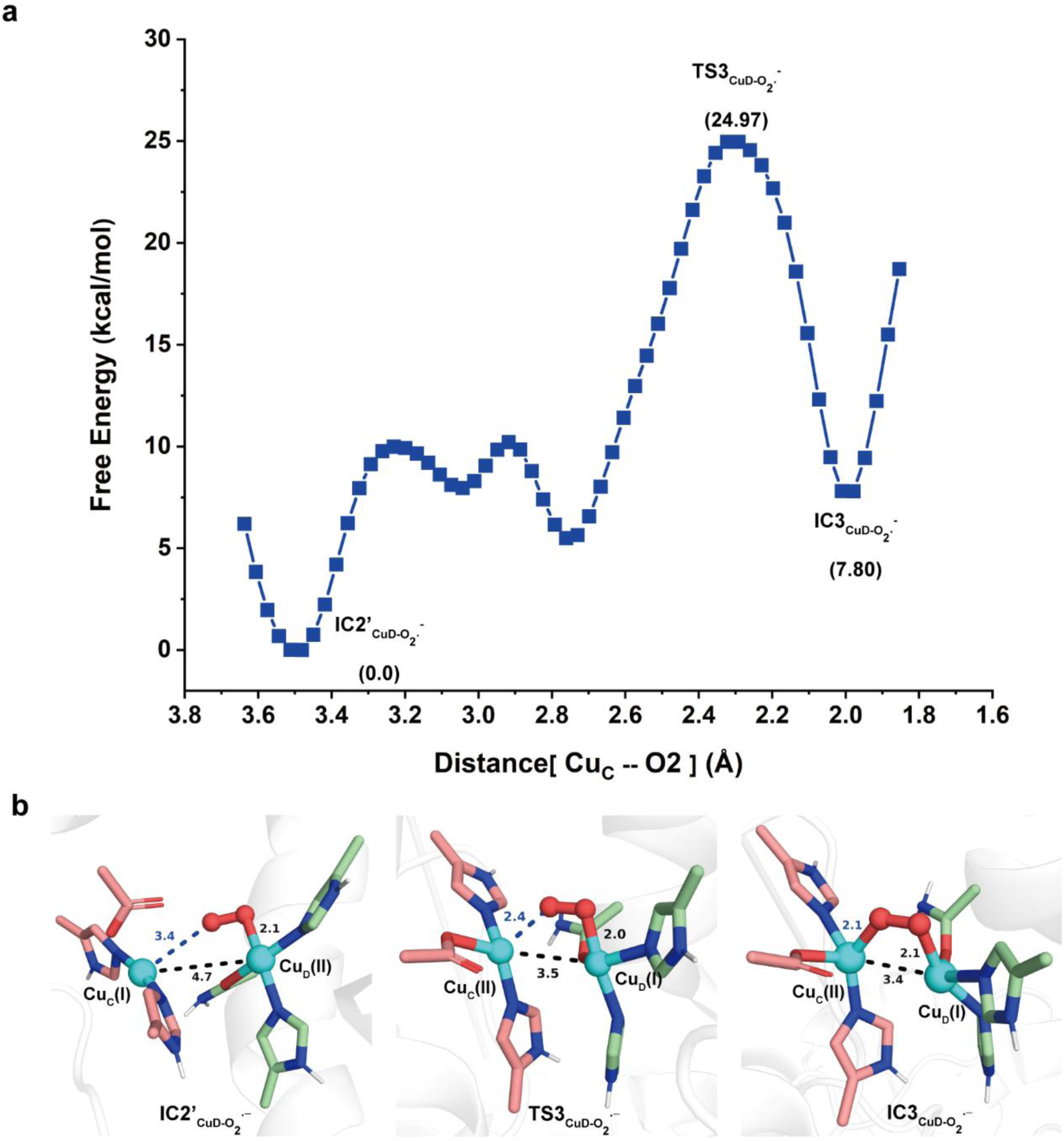
Calculated mechanisms of the (*μ*-superoxo)Cu_C_(II)Cu_D_(I) intermediate formation. **a.** QM/MM metadynamics-calculated free energy profile (in kcal/mol) describing distal-oxygen coordination of Cu_D_−O_2_^•−^ to the Cu_C_ site, concomitant with electron transfer from Cu_C_(I) to Cu_D_(II) sites, culminating in the formation of the (*μ*-superoxo)Cu_C_(II)Cu_D_(I) species (IC2’_CuD−O2_^•−^ → IC3_CuD−O2_^•−^). The reaction coordinate is defined as the distance between Cu_C_ site and the distal oxygen of Cu_D_(II)−O_2_^•−^. **b.** The geometry of intermediates and transition states, including the structures of IC2’_CuD−O2_^•−^, TS3_CuD−O2_^•−^ and IC3_CuD−O2_^•−^. Key distances are given in Å. Free energy calculations reveal that the (*μ*-superoxo)Cu_C_(II)Cu_D_(I) species formation involves a substantial barrier of 24.97 kcal/mol and is thermodynamically unfavorable.

**Extended Data Figure 26.**
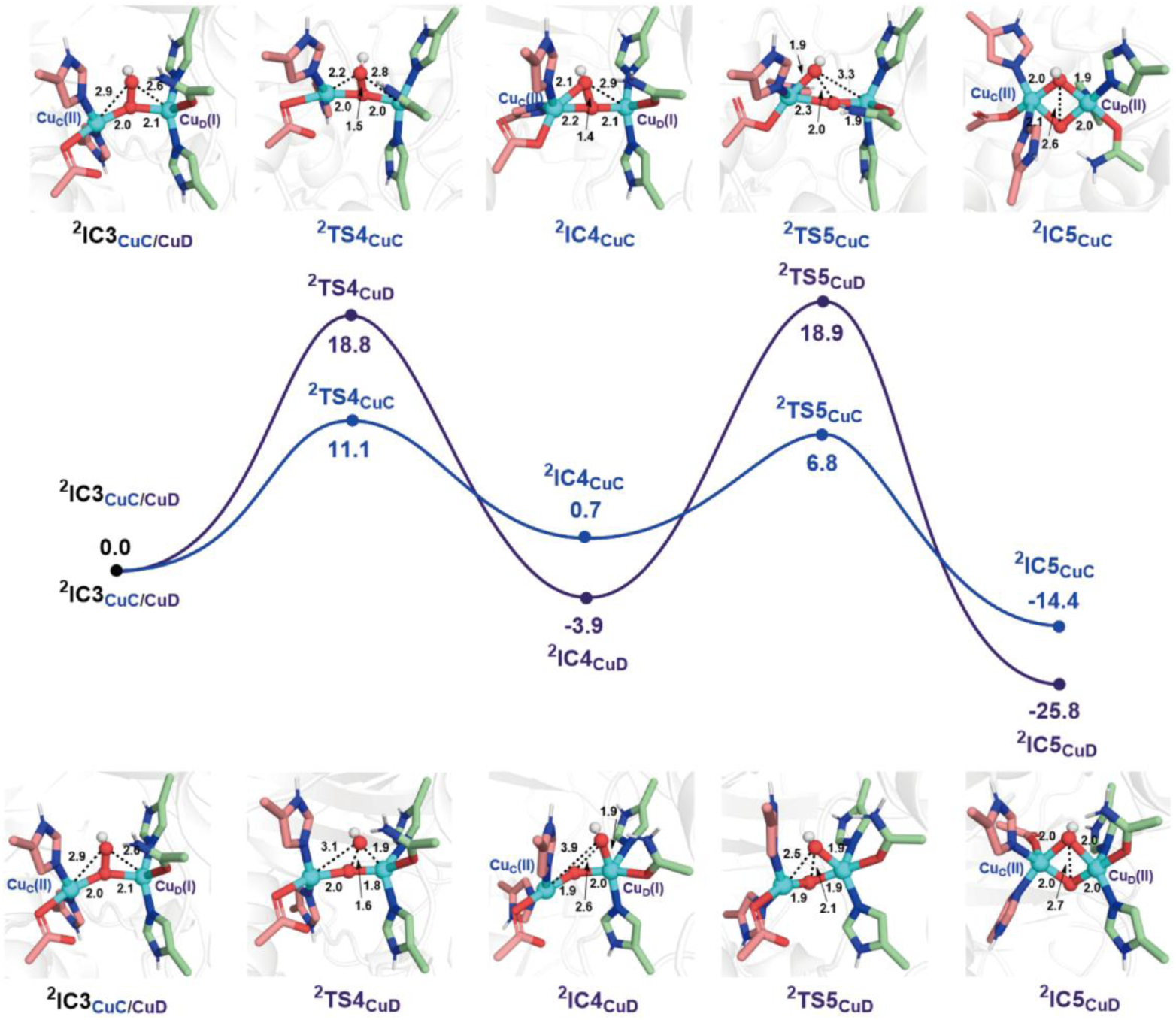
Calculated mechanism of reactive oxygen species generation. QM (UMN15-D3/def2-TZVP)/MM-calculated relative energies (kcal/mol) for reactive oxygen species generation (^2^IC5), which processes rearrangement of the OOH⁻ moiety within the (*μ*-hydroperoxo)Cu_C_(II)Cu_D_(I) specie (^2^IC3 → ^2^IC4). QM(UMN15-D3/def2-SVP)/MM optimized geometries of key species involved in the reaction are presented. Key distances are given in Å. QM/MM calculations indicate that, within the (*μ*-hydroperoxo)Cu_C_(II)Cu_D_(I) specie (^2^IC3), the distal oxygen of the OOH⁻ moiety preferentially coordinates to the Cu_C_ site (^2^IC4_CuC_) rather than to Cu_D_ (^2^IC4_CuD_), ultimately leading to the formation of the (*μ*-oxo)(*μ*-hydro)Cu_C_(II)Cu_D_(II) species (^2^IC5_CuC_).

**Extended Data Figure 27.**
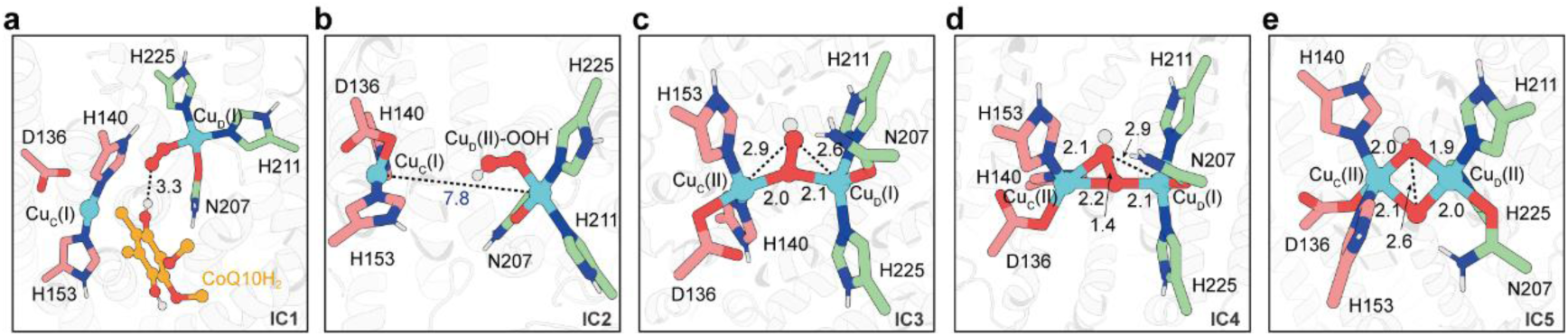
Catalytic intermediates of dicopper center. **a-e.** Structures of key intermediates that capture the stepwise formation of the bimetallic Cu-based reactive oxygen species.

**Extended Data Figure 28.**
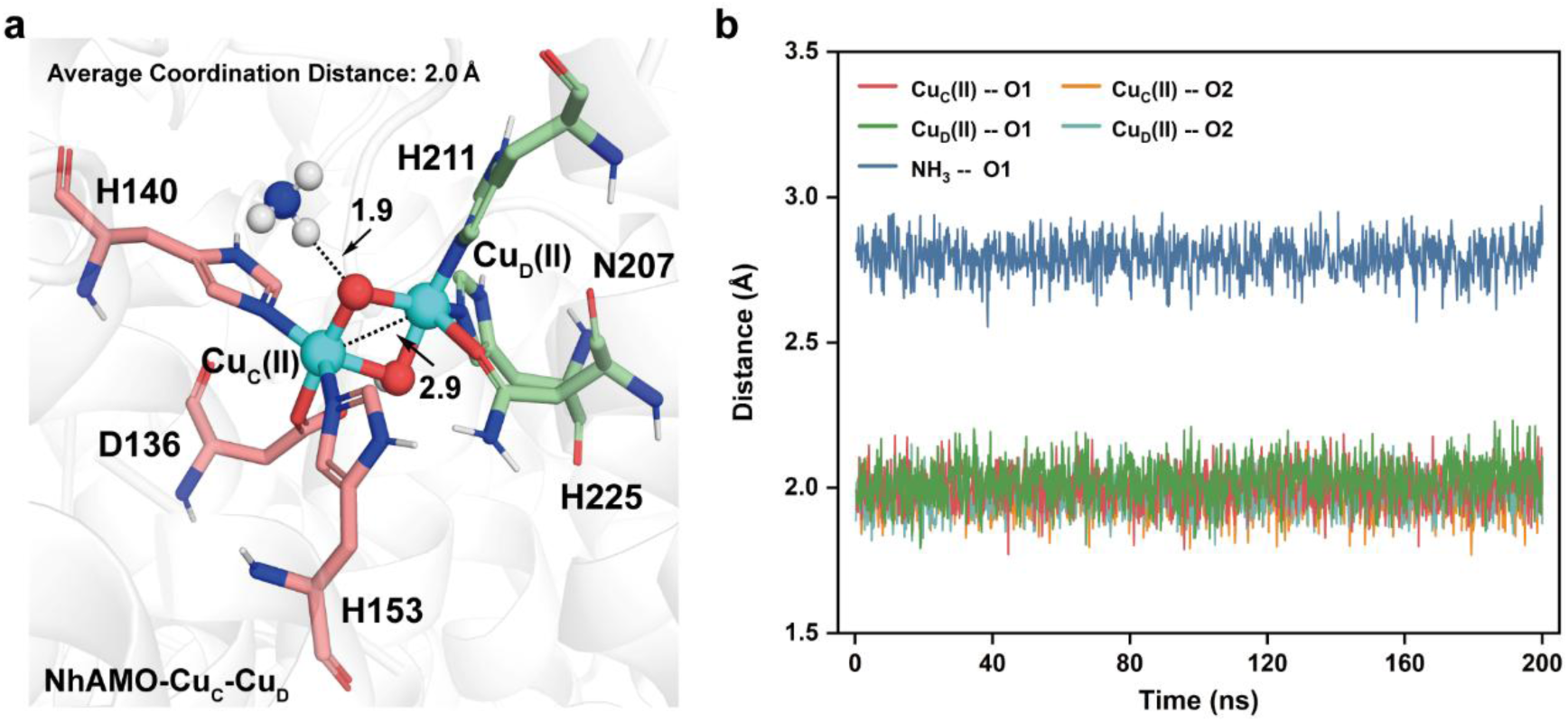
The binding modes between potential *Nh*AMO reactive oxygen species and the substrate ammonia. **a.** MD-averaged binding conformations of the bimetallic Cu-based reactive oxygen species (*μ*-oxo)(*μ*-hydro)Cu_C_(II)Cu_D_(II) and its complex with ammonia. **b.** Time-dependent changes in key coordination distances and in the distances from ammonia to the bimetallic Cu-based species (*μ*-oxo)(*μ*-hydro)Cu_C_(II)Cu_D_(II) species.

**Extended Data Figure 29.**
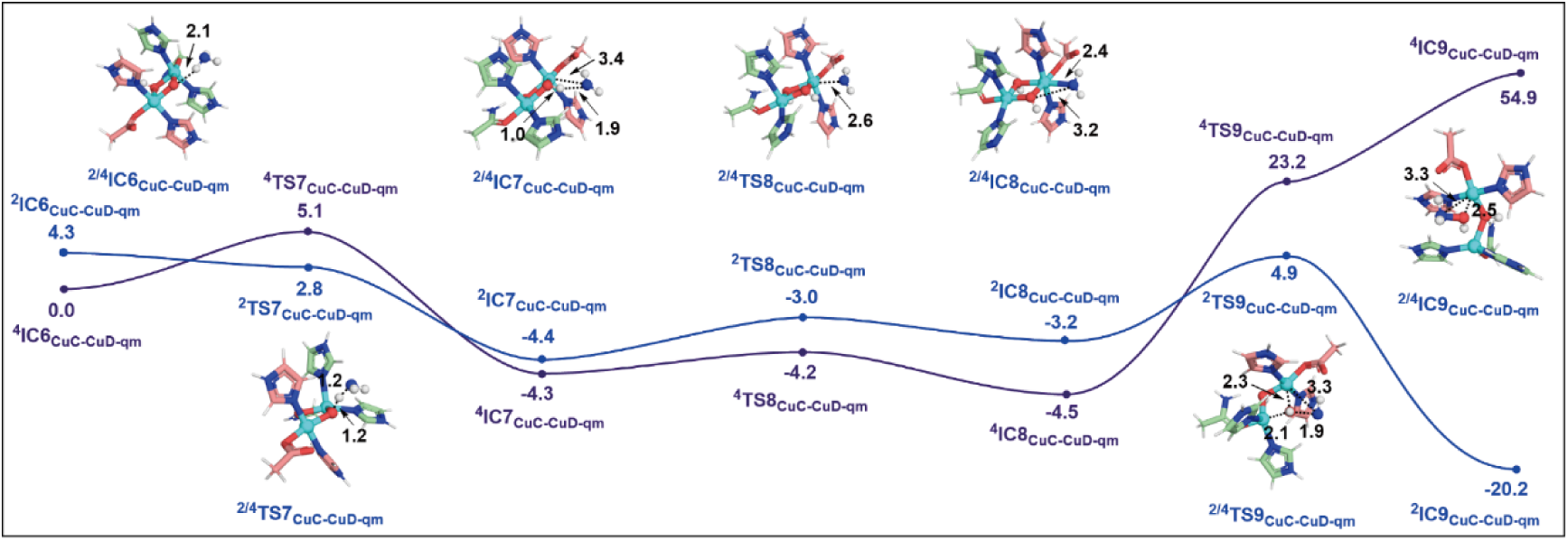
Comparison of the catalysis of potential *Nh*AMO reactive oxygen species. QM (UB3LYP/def2-TZVP//def2-SVP) calculated potential energy profile (in kcal/mol) for (*μ*-oxo)(*μ*-hydro)Cu_C_(II)Cu_D_(II) mediated ammonia hydroxylation to hydroxylamine via the HAT and oxygen rebound mechanism in both doublet and quartet states. Key distances are given in Å. Starting from the *μ*-oxo-*μ*-hydroxo dicopper configuration (IC6_CuC–CuD-qm_), the HAT from NH_3_ requires an energy barrier of 2.8 kcal·mol^-1^, followed by a nearly barrierless NH_2_^•^ rebound step. The final coupling between the singly coordinated amino group and the doubly coordinated hydroxyl proceeds with a 9.4 kcal·mol^-1^ barrier to produce NH_2_OH (IC9_CuC–CuD-qm_). Compared with mononuclear copper centers, the dinuclear Cu_C_–Cu_D_ species promotes NH_3_ oxidation to NH_2_OH more efficiently, highlighting the catalytic potential of the dual-copper site.

**Extended Data Figure 30.**
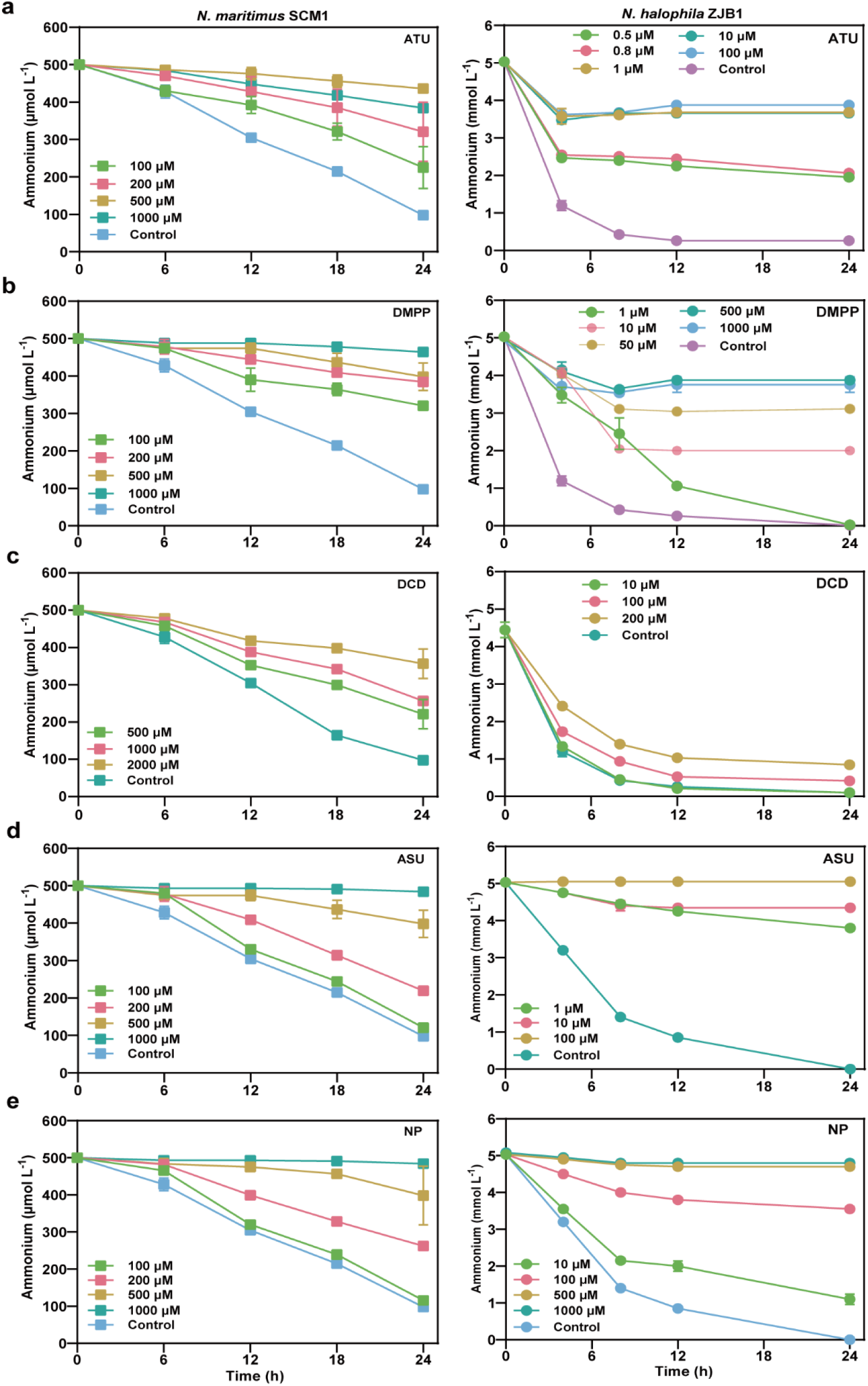
Average inhibition of ammonia oxidation in AOA (left panels) and AOB (right panels) under varying concentrations of NIs. Each subplot corresponds to a specific NI: (**a**) ATU, (**b**) DMPP, (**c**) DCD, (**d**) ASU and (**e**) NP. Different colors and symbols represent different concentrations of the respective inhibitor. Control treatments lack NIs. Error bars indicate standard deviations derived from ≥3 biological replicates, each performed in triplicate.

**Extended Data Figure 31.**
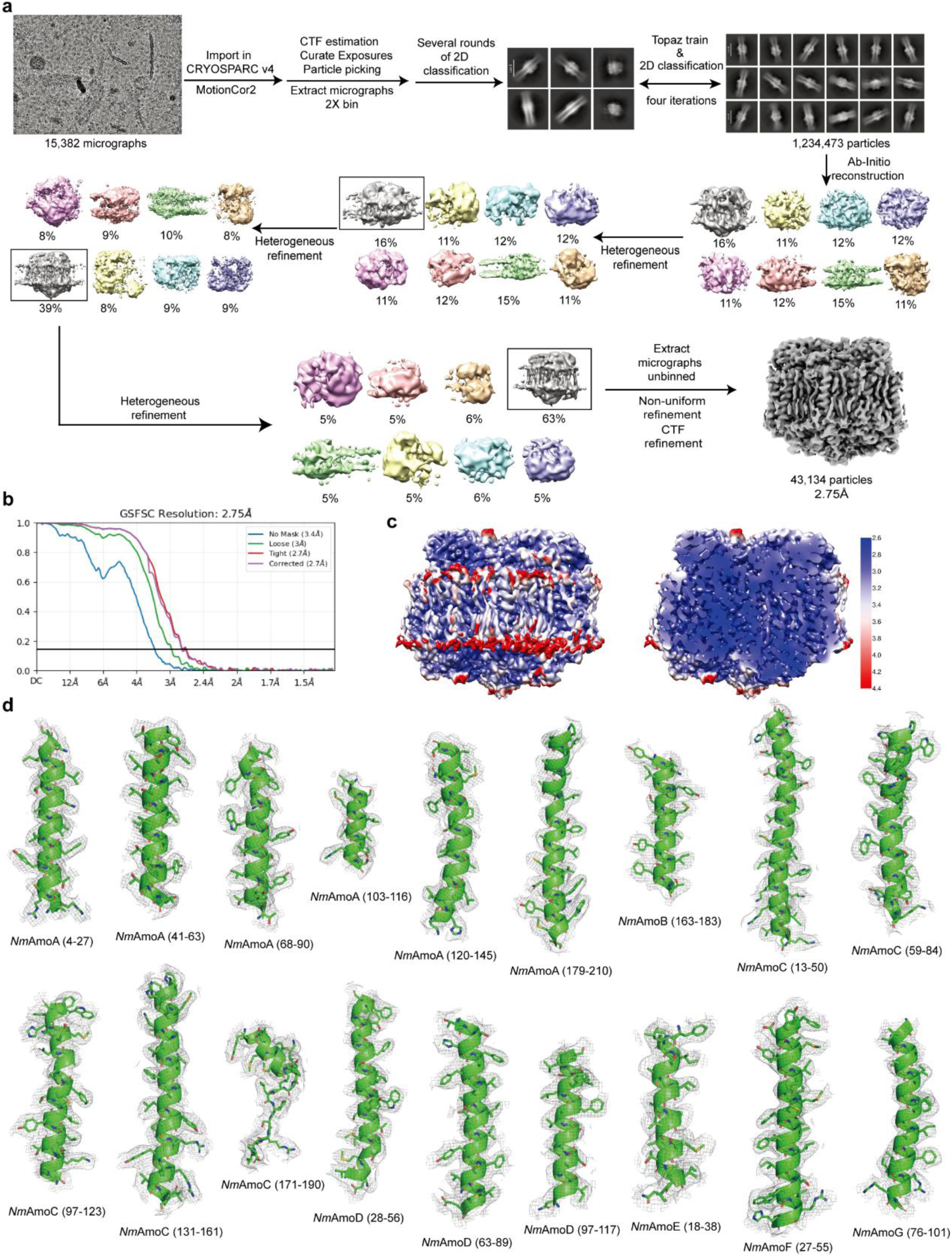
Cryo-EM data analysis of inactivated *Nm*AMO. **a.** The flowchart of cryo-EM data processing of inactivated *Nm*AMO. Details can be found in the ‘Method’ section. **b.** FSC curves for the cryo-EM map of inactivated *Nm*AMO. The threshold of 0.143 was used to determine the overall resolution of the map. **c.** Local resolution maps for the overall reconstruction (left) and a central slice (right) of inactivated *Nm*AMO. **d.** Representative EM maps for subunits from inactivated *Nm*AMO.

**Extended Data Figure 32.**
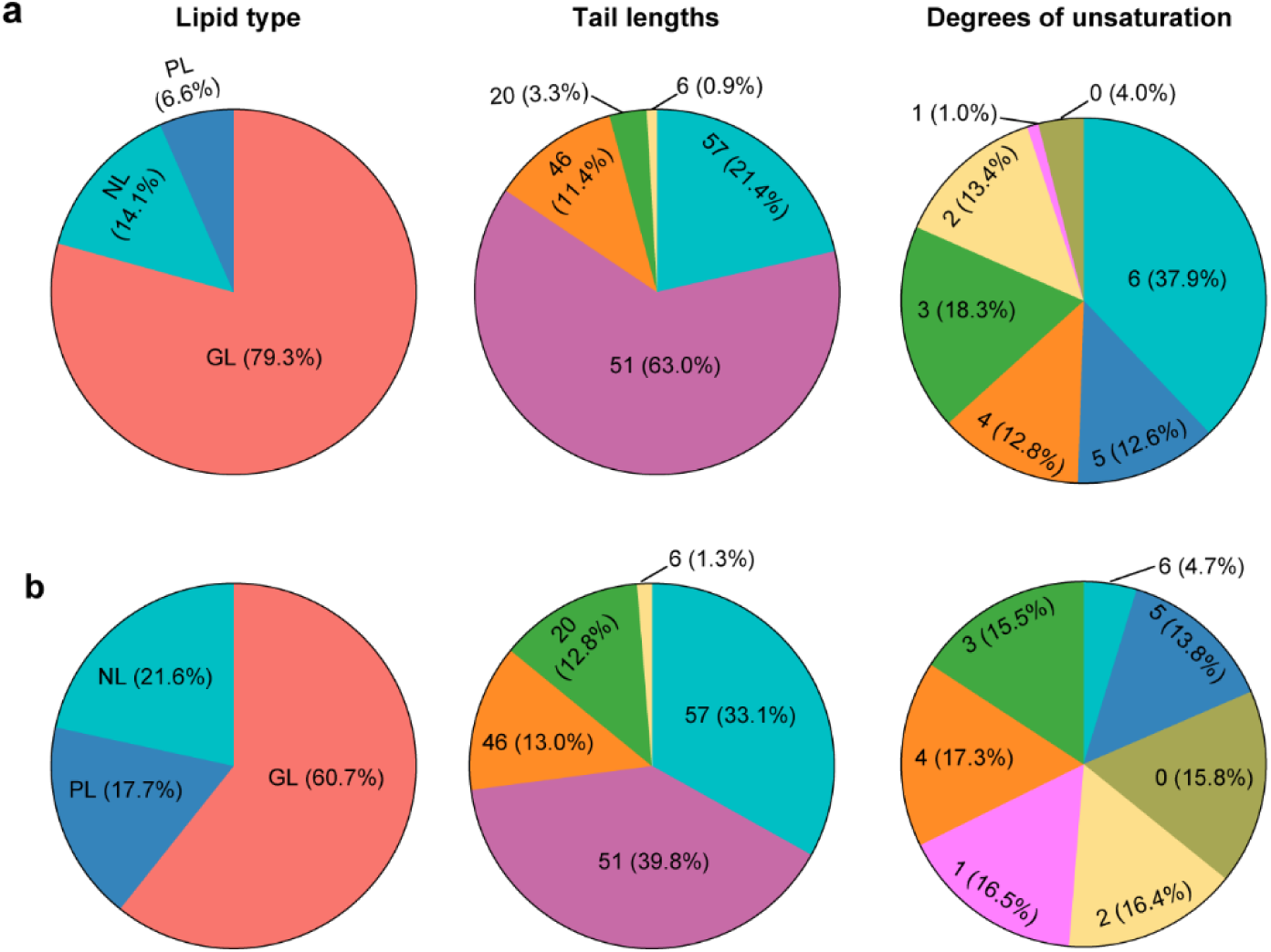
Analysis of the lipidomic data of the SCM1 cells (a) and extracted membrane fraction (b). Lipids are categorized based on the head group type, the degree of unsaturation in lipid tails, and the length of the lipid tails. The identified lipid types include glycolipid (GL), phospholipid (PL), and non-polar lipids (NL).

**Extended Data Figure 33.**
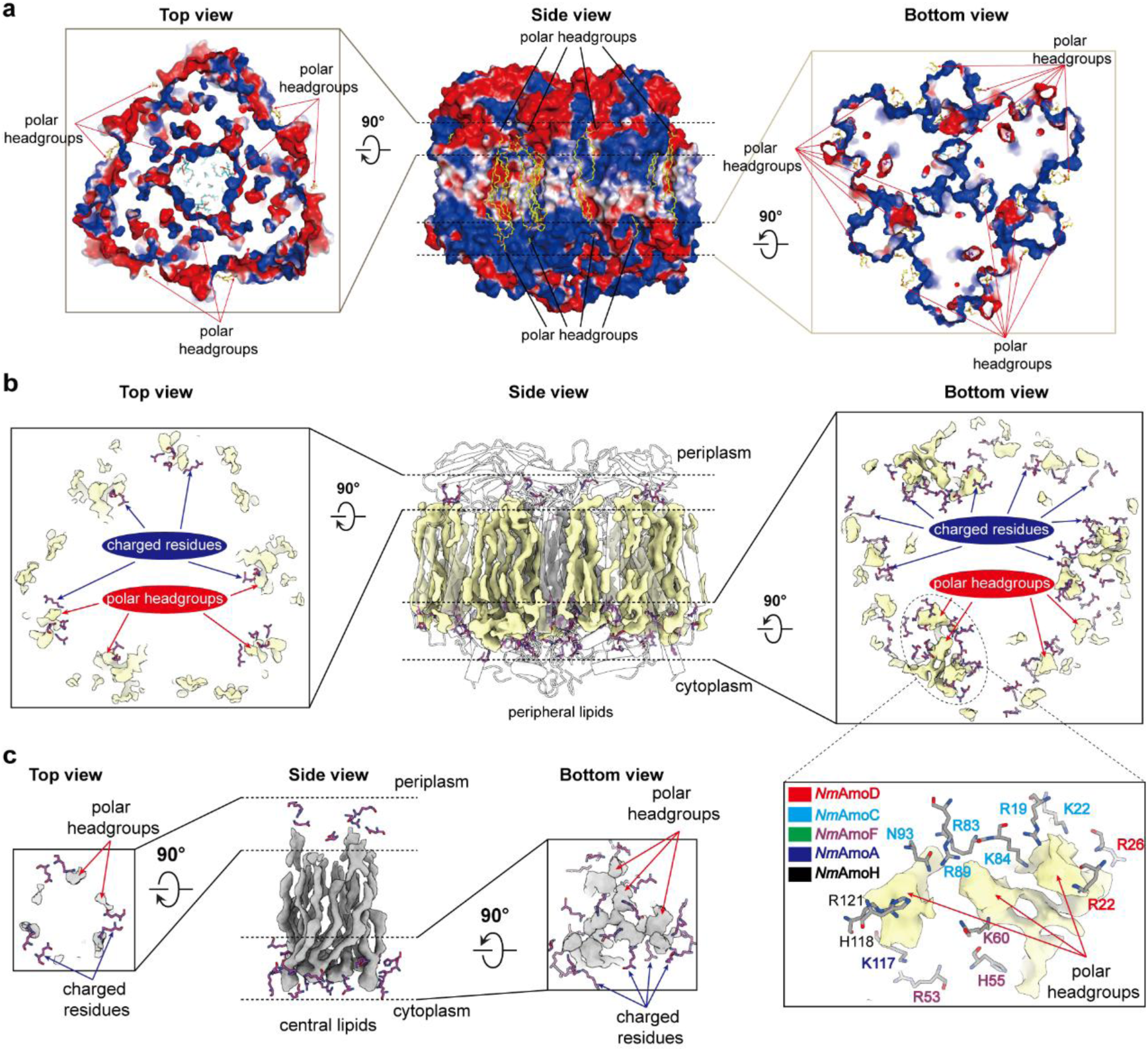
The polar headgroups of archaeal lipids primarily interact with *Nm*AMO via strong electrostatic interactions. **a.** Surface electrostatic potential of *Nm*AMO showing the distribution of polar headgroups of archaeal lipids. **b-c.** Periplasmic (left panels) and cytoplasmic (right panels) polar headgroups from peripheral (**b**) and central (**c**) lipids interacting with charged residues from *Nm*AMO. Three perspectives (Top view, Side view, Bottom view) are shown.

**Extended Data Figure 34.**
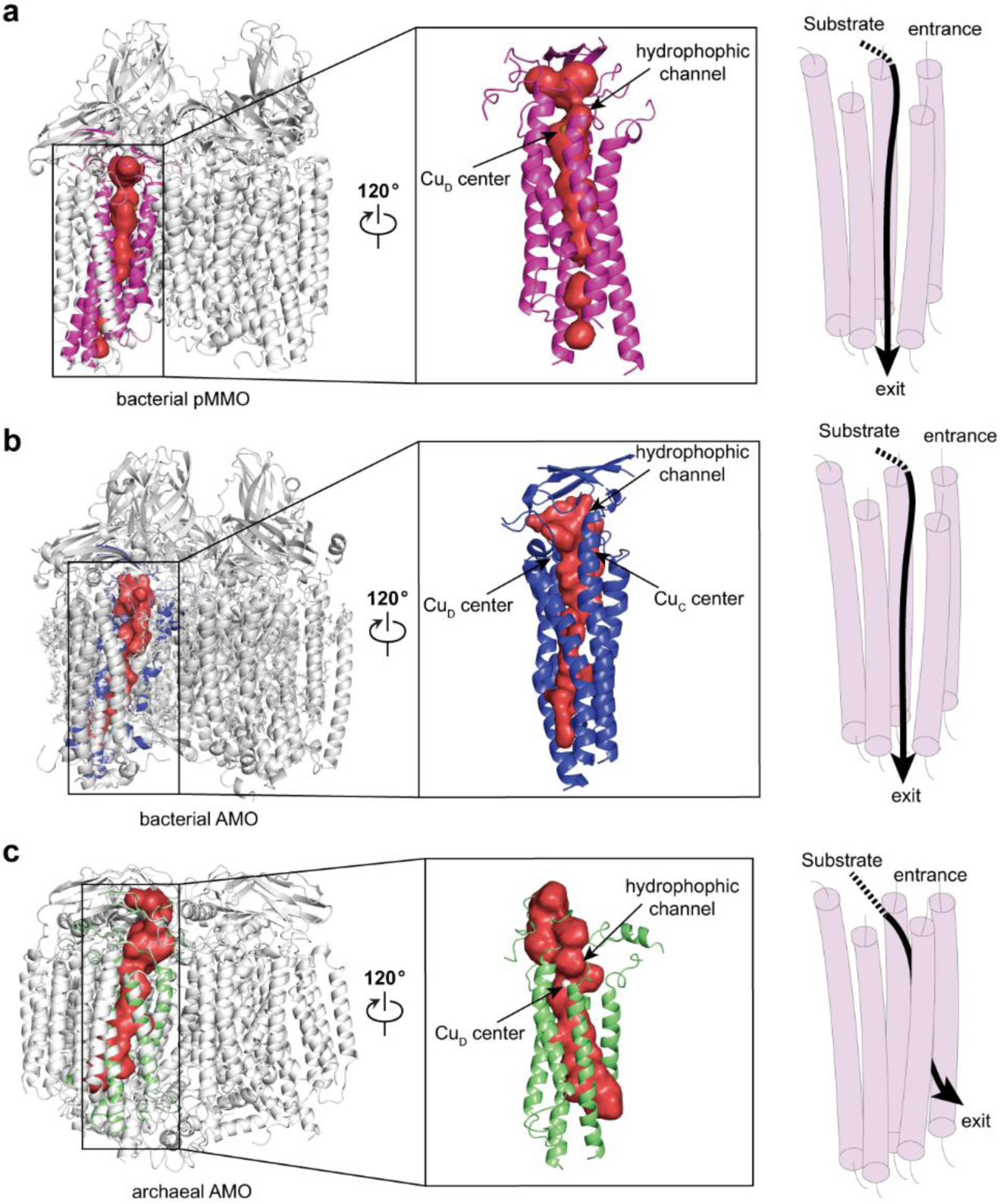
Structural comparison of hydrophobic channels and potential substrate pathways in pMMO and AMOs. The overall structures (left panels), calculated hydrophobic channels (middle panels) and schematic representation of the substrate pathway (right panels) of bacterial pMMO **(a)**, and bacterial **(b)** and archaeal **(c)** AMOs. The central pores generated by the ‘hollow’ program are shown as red surface.

**Extended Data Figure 35.**
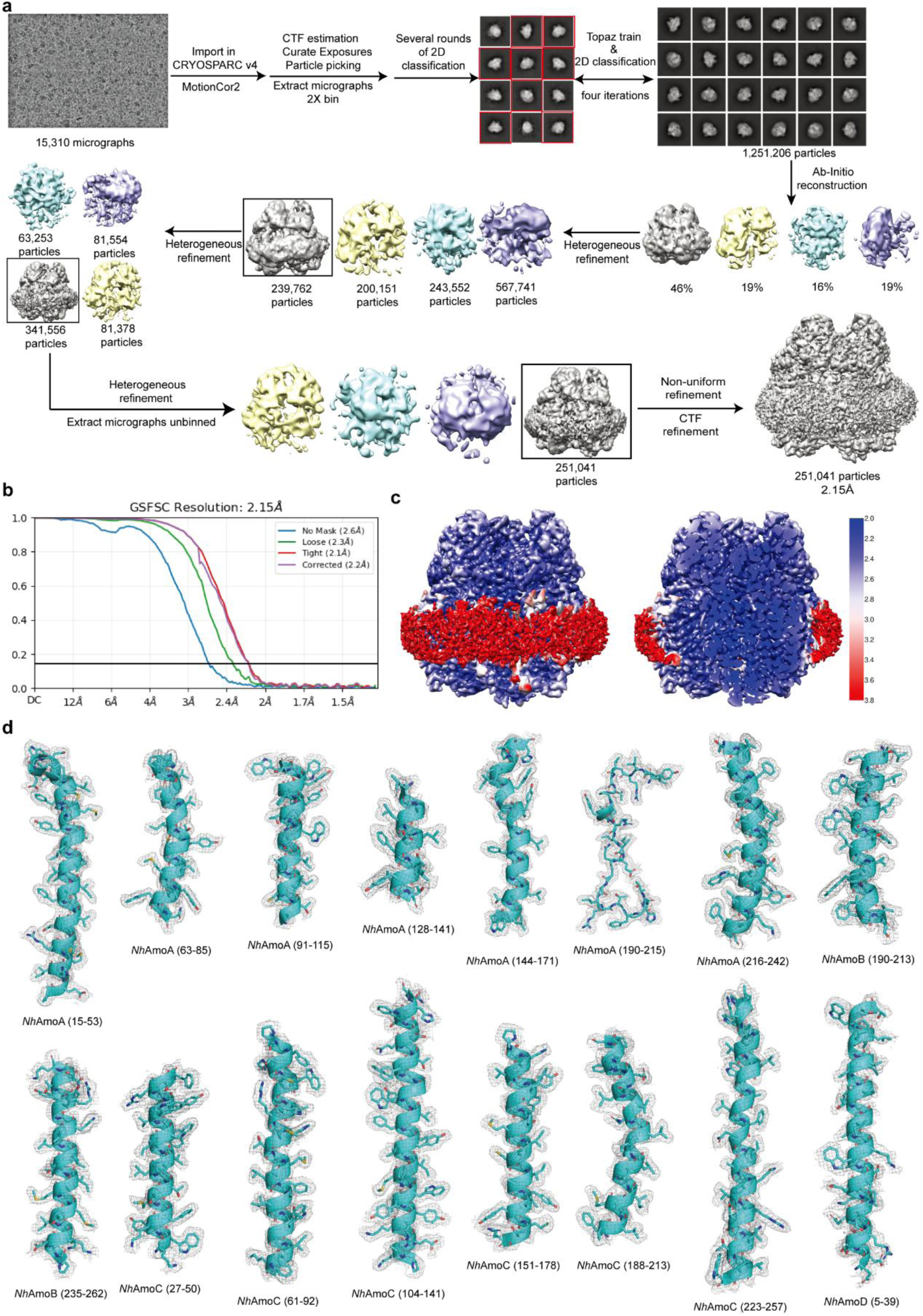
Cryo-EM data analysis of *Nh*AMO-DMP complex. **a.** The flowchart of cryo-EM data processing of *Nh*AMO-DMP complex. Details can be found in the ‘Method’ section. **b.** FSC curves for the cryo-EM map of *Nh*AMO-DMP complex. The threshold of 0.143 was used to determine the overall resolution of the map. **c.** Local resolution maps for the overall reconstruction (left) and a central slice (right) of *Nh*AMO-DMP complex. **d.** Representative EM maps for subunits from *Nh*AMO-DMP complex.

**Extended Data Figure 36.**
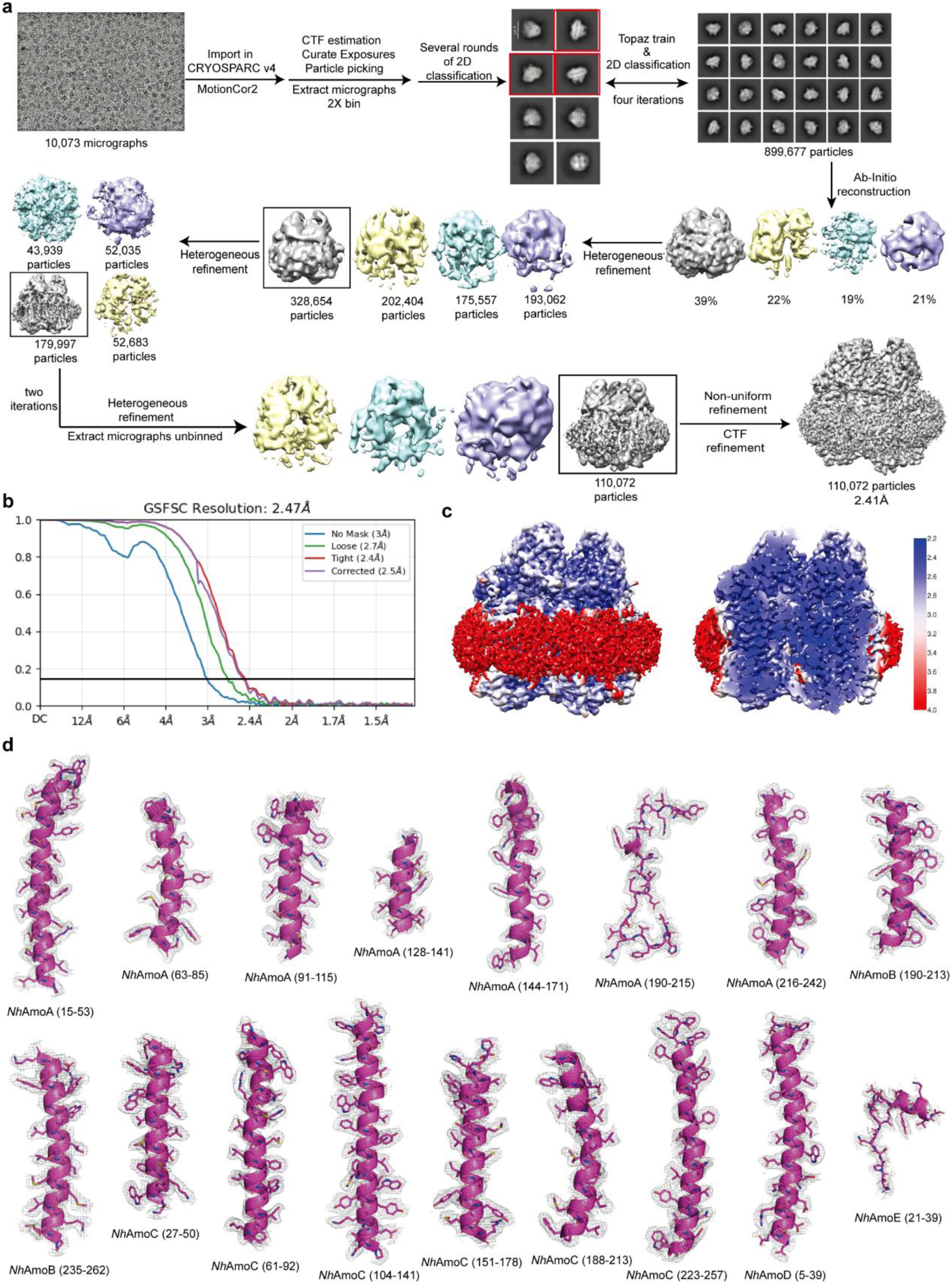
Cryo-EM data analysis of *Nh*AMO-ATU complex. **a.** The flowchart of cryo-EM data processing of *Nh*AMO-ATU complex. Details can be found in the ‘Method’ section. **b.** FSC curves for the cryo-EM map of *Nh*AMO-ATU complex. The threshold of 0.143 was used to determine the overall resolution of the map. **c.** Local resolution maps for the overall reconstruction (left) and a central slice (right) of *Nh*AMO-ATU complex. **d.** Representative EM maps for subunits from *Nh*AMO-ATU complex.

**Extended Data Figure 37.**
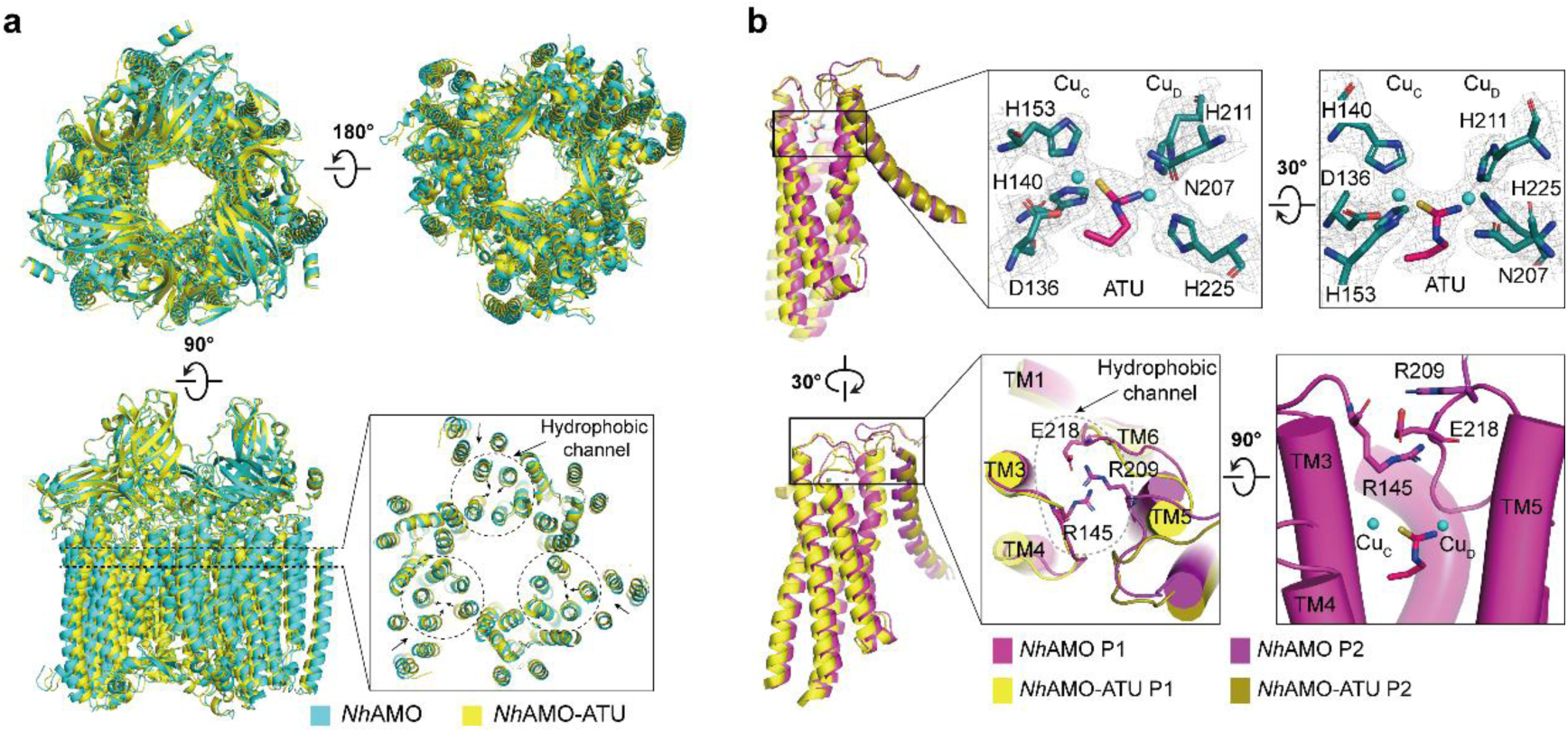
ATU binding induces conformational changes in *Nh*AMO. **a.** Structural superposition of *Nh*AMO and *Nh*AMO-ATU reveals that ATU binding induced the overall contraction of the AMO complex. Contraction of the three hydrophobic channels is highlighted by dashed circles and magnified in (**b**). **b.** ATU bound to the dicopper center, chelated Cu_C_ and Cu_D_, and induced closure and constriction of the hydrophobic channel. Models from adjacent AMO protomers are labeled P1 and P2 and color-coded. These results correspond to those shown in Fig. 6.

**Extended Data Figure 38.**
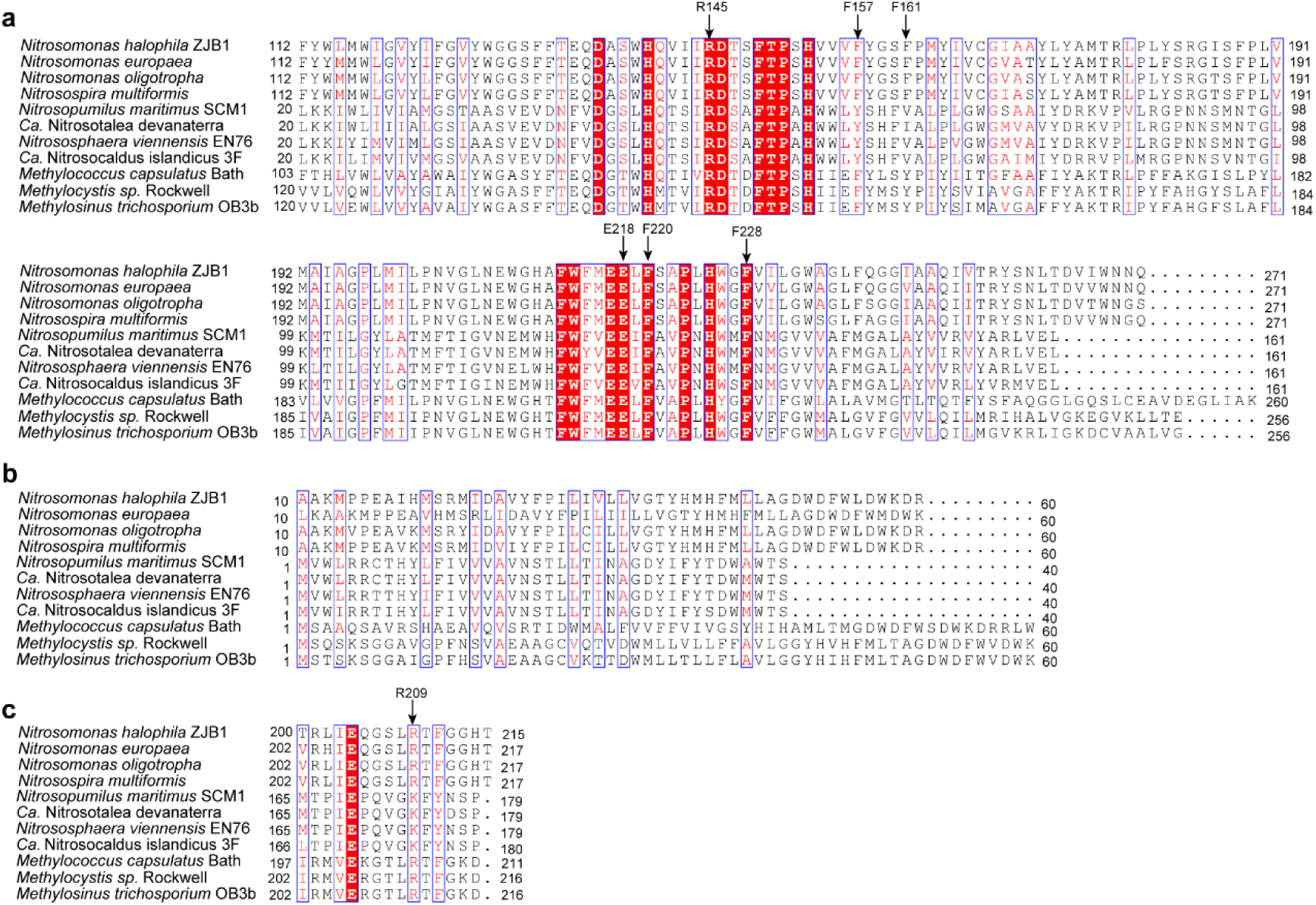
Conservation analysis of the hydrophobic channels and lid regions from ammonia and methane-oxidizing microorganisms. Sequence alignments of the hydrophobic channels and the lid regions in selected AOA, AOB, and methane-oxidizing bacteria. Alignments depict the latter four TMs of the hydrophobic channel in the AmoC/PmoC subunit (**a**), the first TM of the hydrophobic channel in the AmoA/PmoA subunit (**b**), and the adjacent lid regions in the AmoA/PmoA subunit (**c**). The analysis encompassed the following species: AOA—*Nitrosopumilus maritimus* SCM1, *Ca. Nitrosotalea devanaterra*, *Nitrososphaera viennensis* EN76, *Ca. Nitrosocaldus islandicus* 3F; AOB—*Nitrosomonas halophila* ZJB1, *Nitrosomonas europaea*, *Nitrosomonas oligotropha*, *Nitrosospira multiformis*; methane-oxidizing bacteria—*Methylococcus capsulatus* (Bath), *Methylocystis sp.* Rockwell and *Methylosinus trichosporium* OB3b. Numbers flanking the sequences denote the residue positions within the respective full-length protein.

**Extended Data Figure 39.**
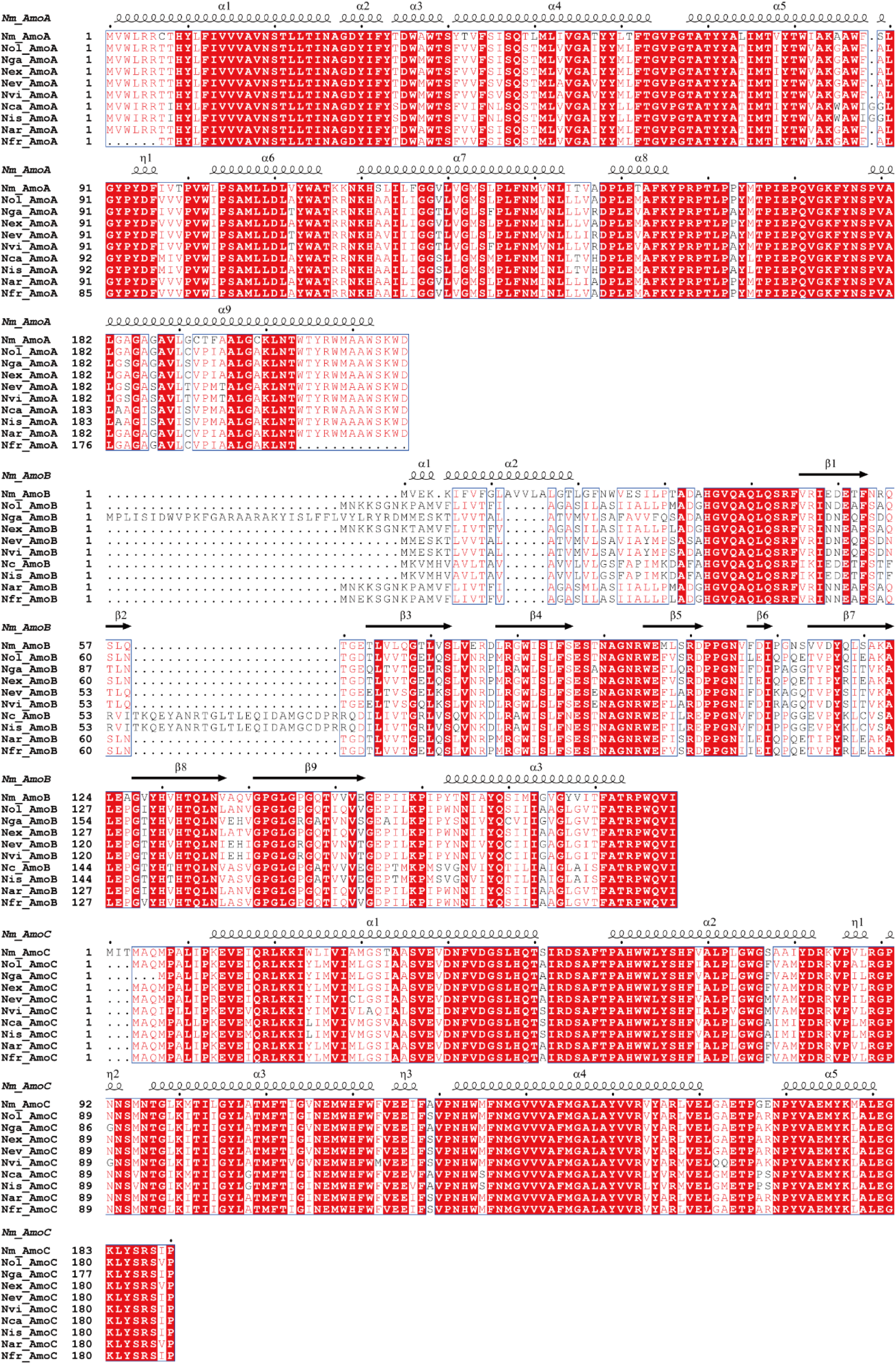
Sequence conservation of core AMO subunits across AOA. Multiple sequence alignments of the core archaeal AMO subunits AmoA, AmoB, and AmoC from representative AOA lineages. Secondary-structure elements are shown above the alignments. Conserved residues are highlighted, indicating strong conservation of the core AMO framework across marine and terrestrial AOA. Nm: *N. maritimus* SCM1, Nol: *Candidatus Nitrosocosmicus oleophilus* MY3, Nga: *Candidatus Nitrososphaera gargensis*, Nex: *Candidatus nitrosocosmicus exaquare*, Nev: *Candidatus Nitrososphaera evergladensis* SR1, Nvi: *Nitrososphaera viennensis* EN76, Nca: *Nitrosocaldus cavascurensis* SCU2, Nis: *Candidatus Nitrosocaldus islandicus* 3F, Nar: *Candidatus_Nitrosocosmicus_arcticus*, Nfr: *Candidatus Nitrosocosmicus franklandus* NFRAN1.

**Extended Data Figure 40.**
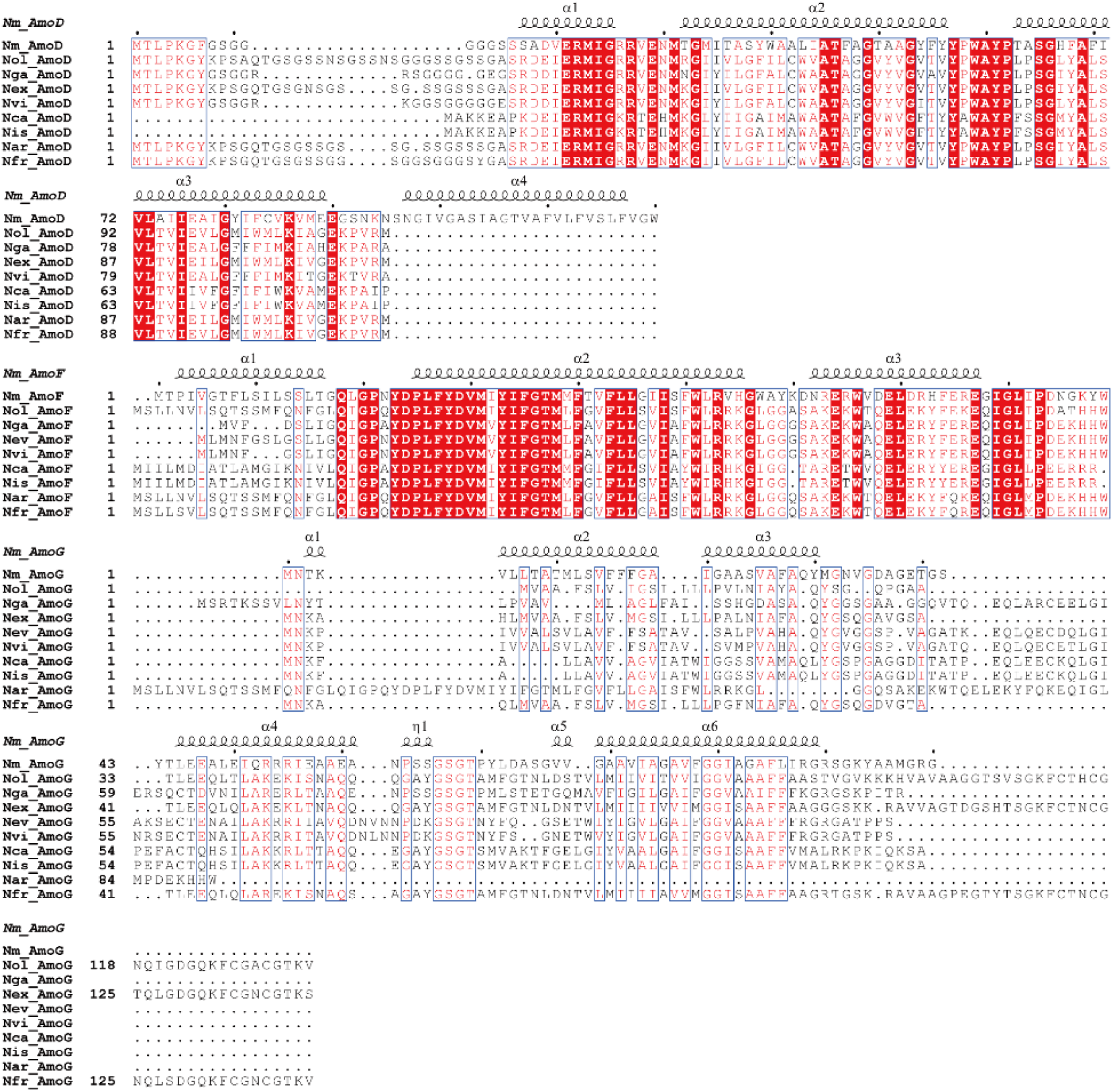
Sequence conservation of accessory AMO subunits across AOA. Multiple sequence alignments of the archaeal AMO accessory subunits AmoD, AmoF, and AmoG from representative AOA lineages. Conserved sequence regions are highlighted, supporting conservation of accessory subunit architecture among diverse terrestrial AOA.

**Extended Data Figure 41.**
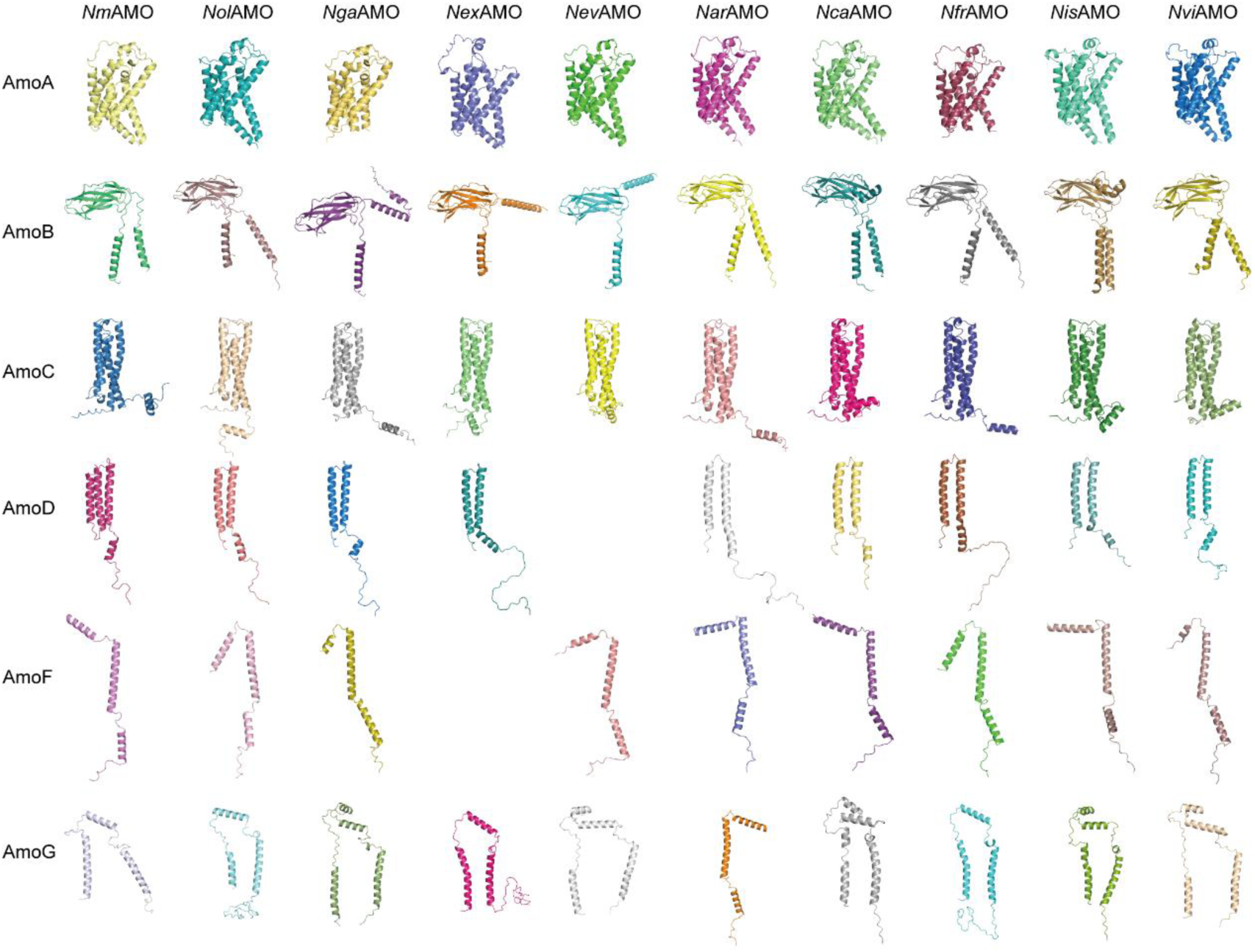
Structural comparison of archaeal AMO subunits. Comparison of experimentally determined *Nm*AMO subunits with AlphaFold-predicted homologous subunits from representative terrestrial AOA. The core and accessory subunits exhibit similar predicted folds across AOA lineages, supporting broad conservation of the archaeal AMO structural framework.

**Extended Data Figure 42.**
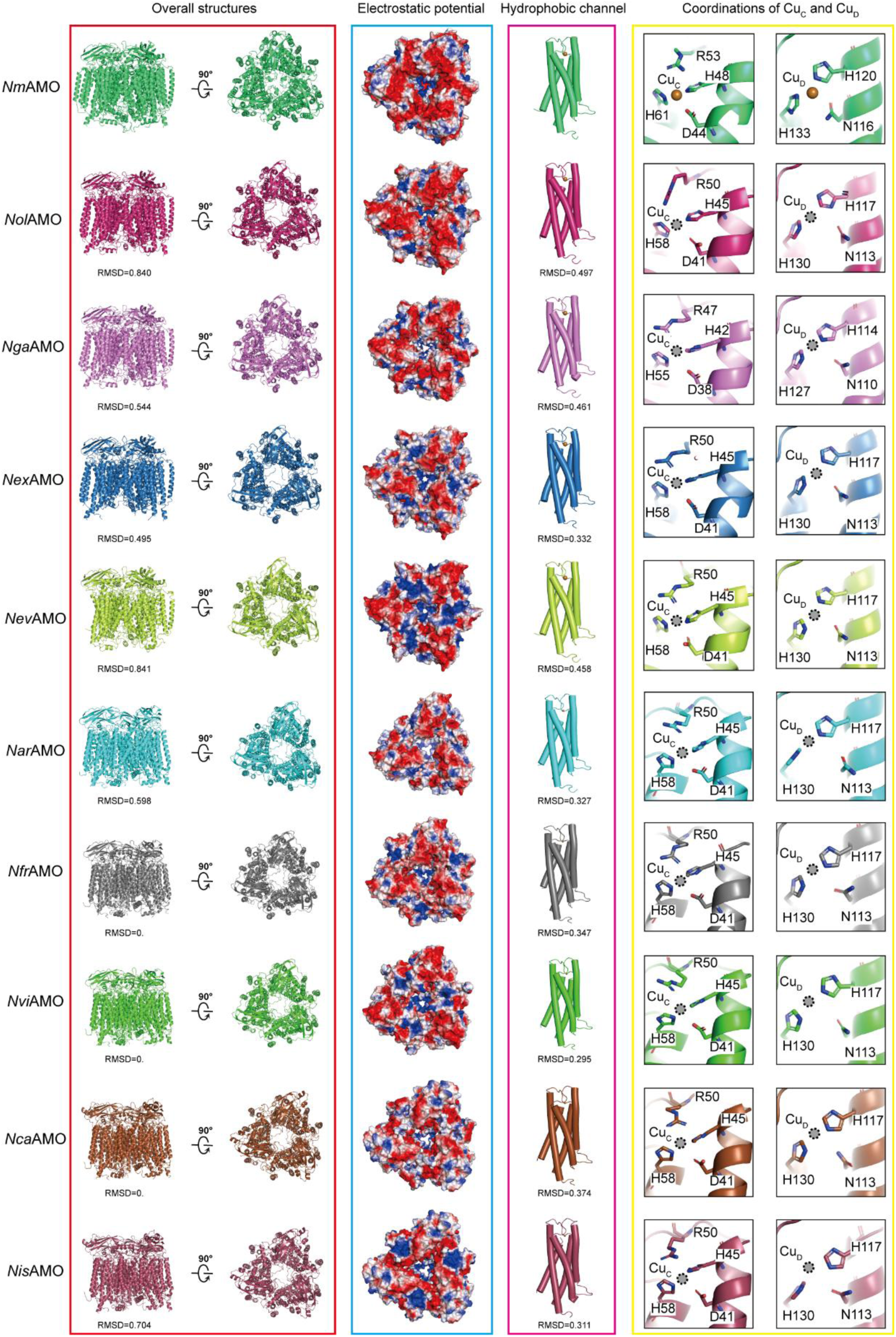
Conservation of overall architecture, surface electrostatics, hydrophobic channel, and copper coordination in archaeal AMOs. Comparison of *Nm*AMO with AlphaFold-predicted AMO models from representative AOA. Overall trimeric architectures, electrostatic surface potentials, hydrophobic channels, and Cu_C_/Cu_D_ coordination environments are shown. The conserved channel architecture and copper-coordinating residues support the use of *Nm*AMO as a structural framework for understanding archaeal AMOs across terrestrial AOA lineages.

**Extended Data Figure 43.**
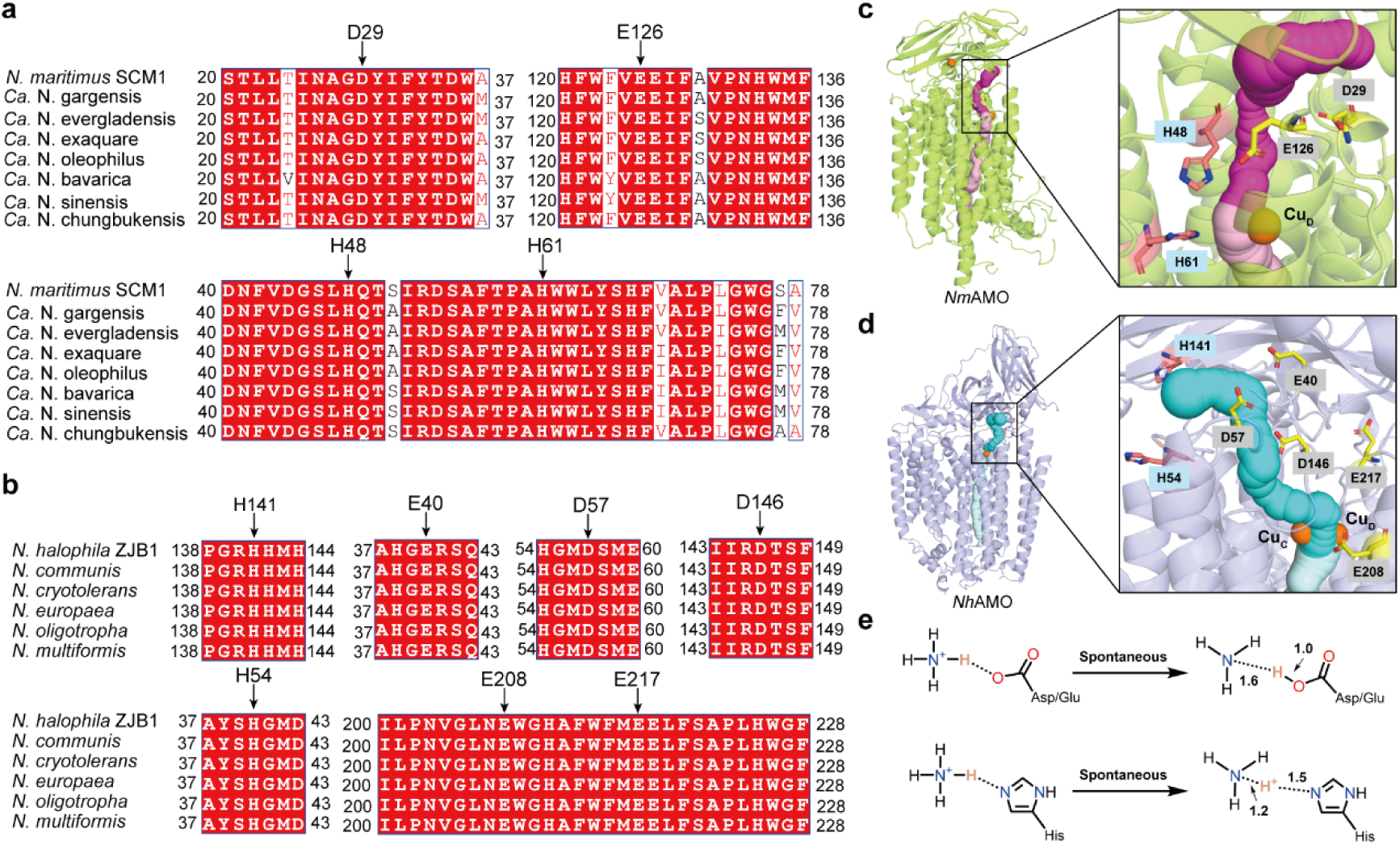
Potential proton acceptors during ammonium translocation in *Nm*AMO and *Nh*AMO. **a-b.** Sequence alignments showing conserved acidic and histidine residues near the putative substrate-translocation pathways in *Nm*AMO (**a**) and *Nh*AMO (**b**). **c-d.** Structural locations of these residues along the hydrophobic channels of *Nm*AMO (**c**) and *Nh*AMO (**d**). The channel is shown as a surface, and copper centers are shown as spheres. These conserved Asp, Glu, and His residues may provide local proton-accepting sites during NH_4_^+^ translocation and deprotonation before catalysis. **e.** QM simulations reveal residue-dependent modulation of NH_4_^+^ deprotonation. In the presence of nearby Asp/Glu residues, spontaneous proton transfers from NH_4_^+^ occurs, leading to NH_3_ formation. By contrast, nearby His residues promote proton delocalization and destabilize the NH_4_^+^ state.

**Extended Data Figure 44.**
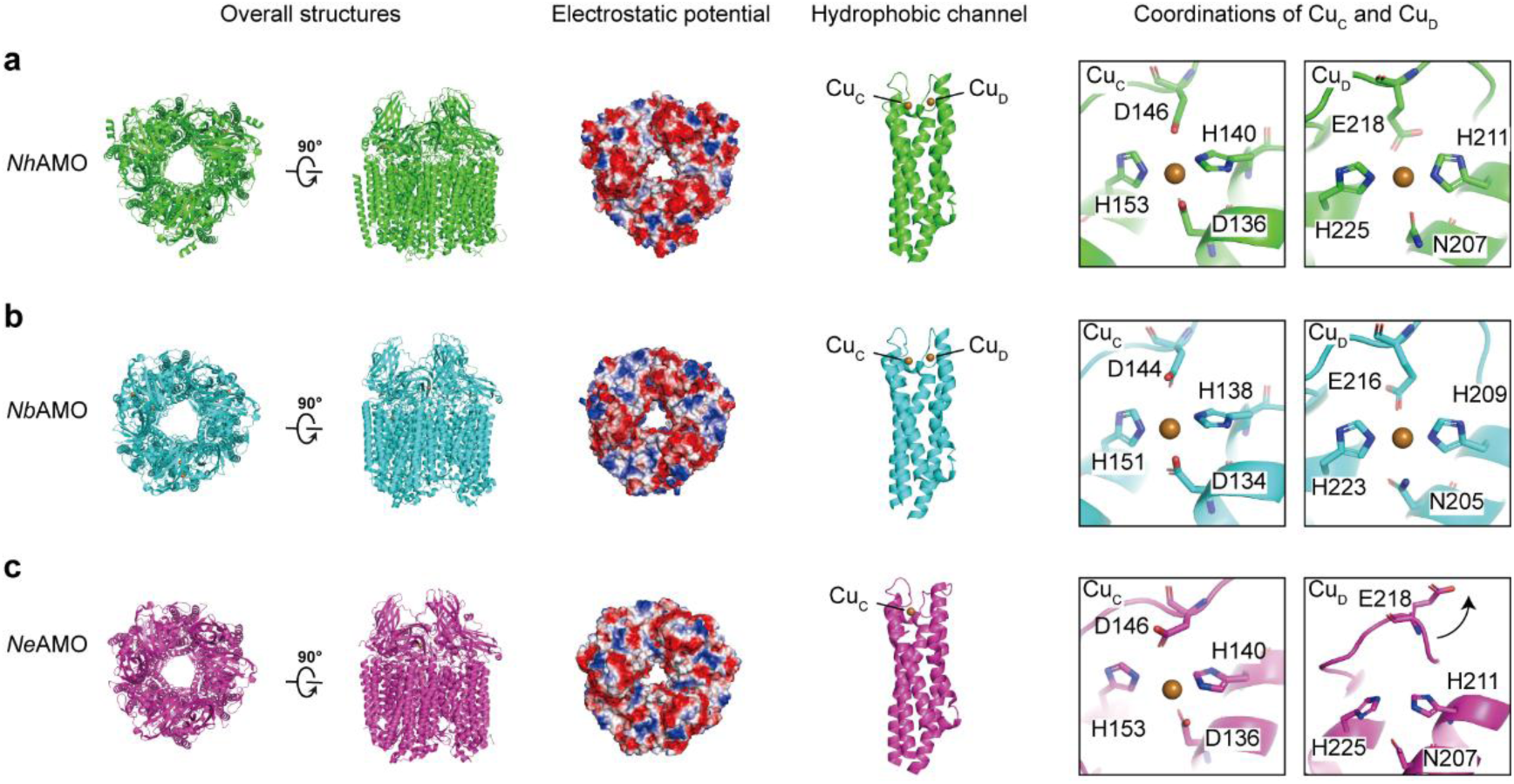
Structural comparison of bacterial AMOs from *N. halophila*, *N. briensis*, and *N. europaea*. Comparison of *Nh*AMO (**a**), *Nb*AMO (**b**), and *Ne*AMO (**c**), including overall structures, electrostatic surface potentials, hydrophobic channels, and Cu_C_/Cu_D_ coordination environments. *Nh*AMO and *Nb*AMO share conserved co-occupied Cu_C_–Cu_D_ dicopper centers, whereas *Ne*AMO shows a distinct Cu_D_-site configuration with Cu_C_ occupancy and altered E218 orientation.

**Extended Data Table 1.** Cryo-EM data collection, refinement and validation statistics.

|  | *Nh*AMO-DMP<br>PDB 9XK5<br>EMDB-66948 | *Nh*AMO-ATU<br>PDB 9XKB<br>EMDB-66954 | active *Nm*AMO<br>PDB 9XJ2<br>EMDB-66926 | inactivated<br>*Nm*AMO<br>PDB 9XGS<br>EMDB-66857 |
| --- | --- | --- | --- | --- |
| **Data collection and processing** |  |  |  |  |
| Magnification | 130,000 | 130,000 | 130,000 | 130,000 |
| Voltage (kV) | 300 | 300 | 300 | 300 |
| Camera | K3 | K3 | K3 | K3 |
| Electron exposure (e<sup>-</sup>/Å<sup>2</sup>) | 50 | 50 | 50 | 50 |
| Defocus range (μm) | -1.5 ~ -2.0 | -1.5 ~ -2.0 | -1.5 ~ -2.0 | -1.5 ~ -2.0 |
| Pixel size (Å) | 0.668 | 0.668 | 0.668 | 0.668 |
| Micrographs (no.) | 15,310 | 10,073 | 16,914 | 15,382 |
| Initial particle images (no.) | 1,251,206 | 899,677 | 391,760 | 1,234,473 |
| Final particle images (no.) | 251,041 | 110,072 | 35,413 | 43,134 |
| Symmetry imposed | C3 | C3 | C3 | C3 |
| Map resolution (Å) | 2.15 | 2.47 | 2.88 | 2.75 |
| Map sharpen B factor (Å<sup>2</sup>) | 72.5 | 80.1 | 92.2 | 90.8 |
| FSC threshold | 0.143 | 0.143 | 0.143 | 0.143 |
| **Refinement** |  |  |  |  |
| Initial model used | 9LEG | 9LEG | 9XGS | - |
| Model resolution (Å) | 2.15 | 2.41 | 2.88 | 2.75 |
| FSC threshold | 0.143 | 0.143 | 0.143 | 0.143 |
| **Model composition** |  |  |  |  |
| Non-hydrogen atoms | 28,820 | 29,868 | 22,305 | 24,327 |
| Protein residues | 2,865 | 2,922 | 2,634 | 2,634 |
| Ligands | 93 | 83 | 24 | 48 |
| Waters | 1023 | 618 | - | - |
| **B factors (Å<sup>2</sup>)** |  |  |  |  |
| Protein | 30.03 | 31.00 | 121.06 | 119.15 |
| Ligand | 20.41 | 20.41 | 122.66 | 140.00 |
| Water | 30.00 | 30.00 | - | - |
| **R.m.s. deviations** |  |  |  |  |
| Bond lengths (Å) | 0.024 | 0.010 | 0.003 | 0.005 |
| Bond angles (°) | 1.062 | 1.302 | 0.753 | 0.696 |
| **Validation** |  |  |  |  |
| MolProbity score | 1.67 | 2.29 | 2.46 | 2.32 |
| Clashscore | 4.76 | 8.66 | 13.19 | 9.56 |
| Rotamer outliers (%) | 1.49 | 4.02 | 4.72 | 4.96 |
| **Ramachandran plot** |  |  |  |  |
| Favored (%) | 95.78 | 94.64 | 95.48 | 95.78 |
| Allowed (%) | 4.01 | 4.43 | 4.41 | 4.10 |
| Disallowed (%) | 0.21 | 0.93 | 0.12 | 0.12 |

**Extended Data Table 2.**
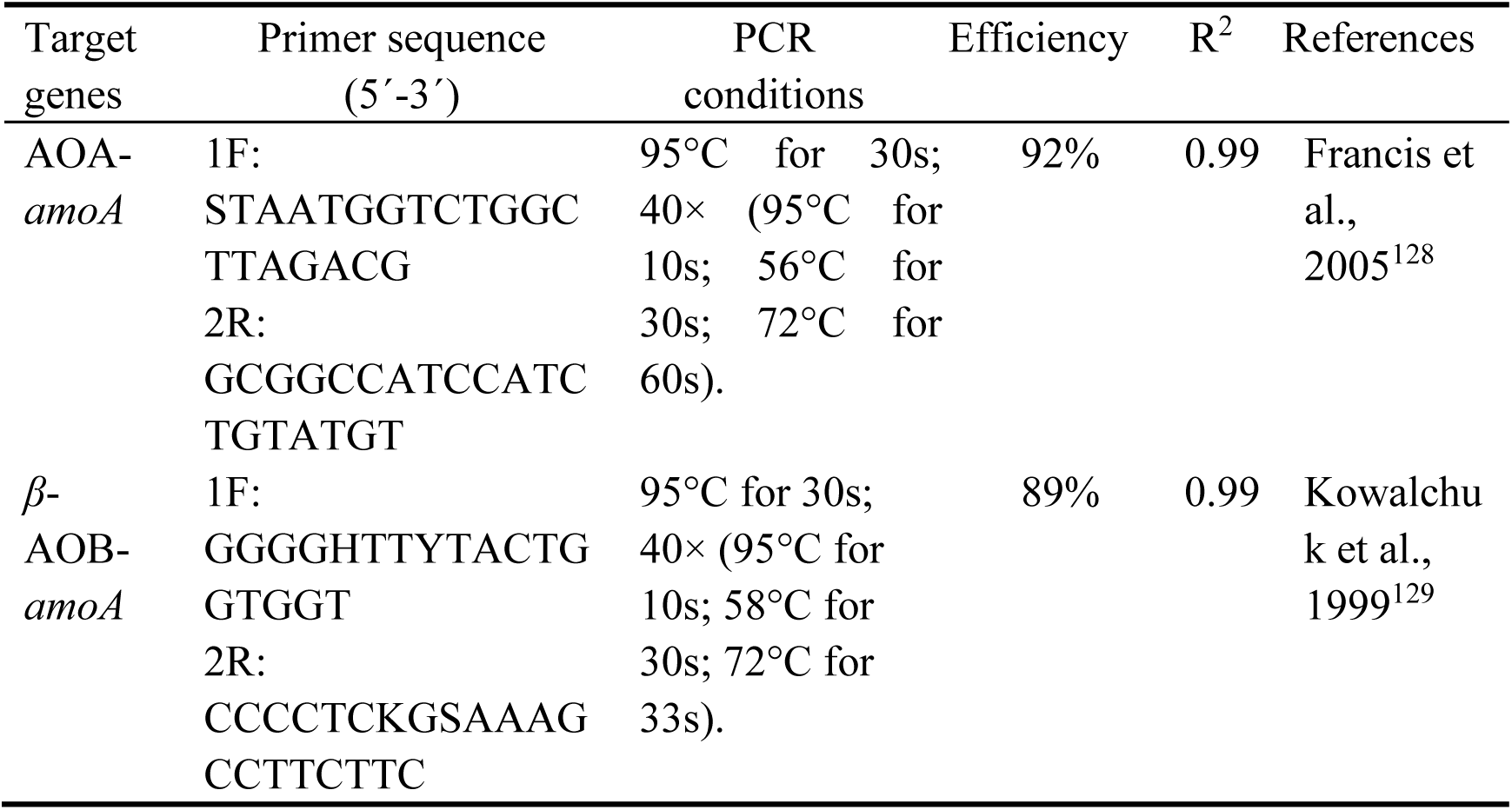
qPCR-based quantification of *amoA* gene abundance in AOA and AOB.

**Extended Data Table 3.** Copper content in active and inactivated *Nm*AMO samples measured by ICP-MS.

| Sample type | Sample name | Sample weight (g) | Final vol. (mL) | Element | Measured conc. (µg/L) | Cu (mg/kg)<sup>a</sup> |
| --- | --- | --- | --- | --- | --- | --- |
| Active *Nm*AMO | AMO_WT-1 | 0.040 | 10 | Cu | 211.52 | 59.14<sup>b</sup> |
|  | AMO_WT-2 | 0.040 | 10 | Cu | 211.65 | 57.18 |
|  | AMO_WT-3 | 0.040 | 10 | Cu | 218.82 | 57.98 |
|  | buffer-1 | 0.0078 | 10 | Cu | 1.32 | 1.69 |
|  | buffer-2 | 0.0078 | 10 | Cu | 1.35 | 1.73 |
|  | buffer-3 | 0.0078 | 10 | Cu | 1.32 | 1.70 |
| Inactivated *Nm*AMO | AMO_ATU-1 | 0.033 | 10 | Cu | 197.93 | 59.98 |
|  | AMO_ATU-2 | 0.033 | 10 | Cu | 204.05 | 61.83 |
|  | AMO_ATU-3 | 0.033 | 10 | Cu | 199.67 | 60.51 |
|  | buffer-1 | 0.0064 | 10 | Cu | 0.37 | 0.59 |
|  | buffer-2 | 0.0064 | 10 | Cu | 0.37 | 0.57 |
|  | buffer-3 | 0.0064 | 10 | Cu | 0.37 | 0.58 |

<sup>a</sup>the net copper mass in the sample was calculated as: $m(\text{Cu})_{\text{net}} = m(\text{Cu})_{\text{sample}} - m(\text{Cu})_{\text{buffer}}$ . Analogous to pMMO, AMO accounted for approximately 80% of the total membrane-bound protein abundance.
<sup>b</sup>the molecular weight of *Nm*AMO trimer is 360.3 kDa.

**Extended Data Table 4.** QM/MM energies (in Hartree, a.u.) of all species.

| UMN15/def2-TZVP//def2-SVP |  |  |  |  |
| --- | --- | --- | --- | --- |
| Species | QM | MM | QM/MM | ZPE |
| <sup>3</sup>IC1<sub>CuD</sub> | -3142.8305 | -56.3701 | -3199.2007 | 0.4495 |
| <sup>3</sup>TS2<sub>CuD</sub> | -3142.7978 | -56.3655 | -3199.1634 | 0.4388 |
| <sup>3</sup>IC2<sub>CuD</sub> | -3142.8207 | -56.3763 | -3199.1971 | 0.4495 |
| <sup>1</sup>IC1<sub>CuD</sub> | -3142.8252 | -56.3697 | -3199.1949 | 0.4899 |
| <sup>1</sup>TS2<sub>CuD</sub> | -3142.7953 | -56.3615 | -3199.1569 | 0.4874 |
| <sup>1</sup>IC2<sub>CuD</sub> | -3298.6746 | -53.7768 | -3352.4514 | 0.4900 |
| <sup>2</sup>IC3<sub>CuC/CuD</sub> | -4774.5514 | -138.6438 | -4913.1951 | 0.4140 |
| <sup>2</sup>TS4<sub>CuC</sub> | -4774.5349 | -138.6386 | -4913.1736 | 0.4101 |
| <sup>2</sup>IC4<sub>CuC</sub> | -4774.5619 | -138.6307 | -4913.1926 | 0.4127 |
| <sup>2</sup>TS5<sub>CuC</sub> | -4774.5354 | -138.6366 | -4913.1720 | 0.4017 |
| <sup>2</sup>IC5<sub>CuC</sub> | -4774.5729 | -138.6391 | -4913.2120 | 0.4081 |
| <sup>2</sup>TS4<sub>CuD</sub> | -4774.5185 | -138.6430 | -4913.1615 | 0.4066 |
| <sup>2</sup>IC4<sub>CuD</sub> | -4774.5637 | -138.6354 | -4913.1991 | 0.4079 |
| <sup>2</sup>TS5<sub>CuD</sub> | -4774.5309 | -138.6308 | -4913.1617 | 0.4070 |
| <sup>2</sup>IC5<sub>CuD</sub> | -4774.6262 | -138.6072 | -4913.2334 | 0.4072 |

**Extended Data Table 5.** UMN15/def2-TZVP//def2-SVP QM energies (in Hartree, a.u.) of the species.

| UMN15/def2-TZVP//def2-SVP |  |  |  |
| --- | --- | --- | --- |
| Species | EE | ZPE | EE+ZPE |
| <sup>1</sup>RC<sub>CuC</sub> | -2452.921 | 0.239 | -2452.682 |
| <sup>1</sup>TS1<sub>CuC</sub> | -2452.909 | 0.233 | -2452.676 |
| <sup>1</sup>IC1<sub>CuC</sub> | -2452.928 | 0.236 | -2452.692 |
| <sup>1</sup>TS2<sub>CuC</sub> | -2452.924 | 0.236 | -2452.688 |
| <sup>1</sup>IC2<sub>CuC</sub> | -2452.938 | 0.240 | -2452.698 |
| <sup>1</sup>TS3<sub>CuC</sub> | -2452.894 | 0.231 | -2452.663 |
| <sup>1</sup>PC<sub>CuC</sub> | -2452.967 | 0.242 | -2452.725 |
| <sup>3</sup>RC<sub>CuC</sub> | -2452.927 | 0.239 | -2452.688 |
| <sup>3</sup>TS1<sub>CuC</sub> | -2452.906 | 0.233 | -2452.673 |
| <sup>3</sup>IC1<sub>CuC</sub> | -2452.928 | 0.236 | -2452.692 |
| <sup>3</sup>TS2<sub>CuC</sub> | -2452.925 | 0.236 | -2452.689 |
| <sup>3</sup>IC2<sub>CuC</sub> | -2452.926 | 0.236 | -2452.690 |
| <sup>1</sup>RC<sub>CuD</sub> | -2433.489 | 0.264 | -2433.225 |
| <sup>1</sup>TS1<sub>CuD</sub> | -2433.471 | 0.258 | -2433.213 |
| <sup>1</sup>IC1<sub>CuD</sub> | -2433.506 | 0.262 | -2433.248 |
| <sup>1</sup>TS2<sub>CuD</sub> | -2433.496 | 0.258 | -2433.238 |
| <sup>1</sup>IC2<sub>CuD</sub> | -2433.509 | 0.263 | -2433.246 |
| <sup>1</sup>TS3<sub>CuD</sub> | -2433.487 | 0.263 | -2433.224 |
| <sup>1</sup>PC<sub>CuD</sub> | -2433.553 | 0.266 | -2433.287 |
| <sup>3</sup>RC<sub>CuD</sub> | -2433.497 | 0.264 | -2433.233 |
| <sup>3</sup>TS1<sub>CuD</sub> | -2433.473 | 0.257 | -2433.216 |
| <sup>3</sup>IC1<sub>CuD</sub> | -2433.507 | 0.262 | -2433.249 |
| <sup>3</sup>TS2<sub>CuD</sub> | -2433.496 | 0.258 | -2433.238 |
| <sup>3</sup>IC2<sub>CuD</sub> | -2433.507 | 0.262 | -2433.245 |
| <sup>2</sup>IC6<sub>CuC-CuD</sub> | -4830.626 | 0.479 | -4830.147 |
| <sup>2</sup>TS7<sub>CuC-CuD</sub> | -4830.605 | 0.473 | -4830.132 |
| <sup>2</sup>IC7<sub>CuC-CuD</sub> | -4830.627 | 0.476 | -4830.151 |
| <sup>2</sup>TS8<sub>CuC-CuD</sub> | -4830.625 | 0.475 | -4830.150 |
| <sup>2</sup>IC8<sub>CuC-CuD</sub> | -4830.626 | 0.476 | -4830.150 |
| <sup>2</sup>TS9<sub>CuC-CuD</sub> | -4830.611 | 0.477 | -4830.134 |
| <sup>2</sup>IC9<sub>CuC-CuD</sub> | -4830.675 | 0.479 | -4830.196 |
| <sup>4</sup>IC6<sub>CuC-CuD</sub> | -4830.627 | 0.479 | -4830.148 |
| <sup>4</sup>TS7<sub>CuC-CuD</sub> | -4830.606 | 0.473 | -4830.133 |
| <sup>4</sup>IC7<sub>CuC-CuD</sub> | -4830.627 | 0.476 | -4830.151 |
| <sup>4</sup>TS8<sub>CuC-CuD</sub> | -4830.626 | 0.475 | -4830.151 |
| <sup>4</sup>IC8<sub>CuC-CuD</sub> | -4830.627 | 0.476 | -4830.151 |
| <sup>4</sup>TS9<sub>CuC-CuD</sub> | -4830.588 | 0.477 | -4830.111 |
| <sup>4</sup>IC9<sub>CuC-CuD</sub> | -4830.549 | 0.482 | -4830.067 |

**Extended Data Table 6.**
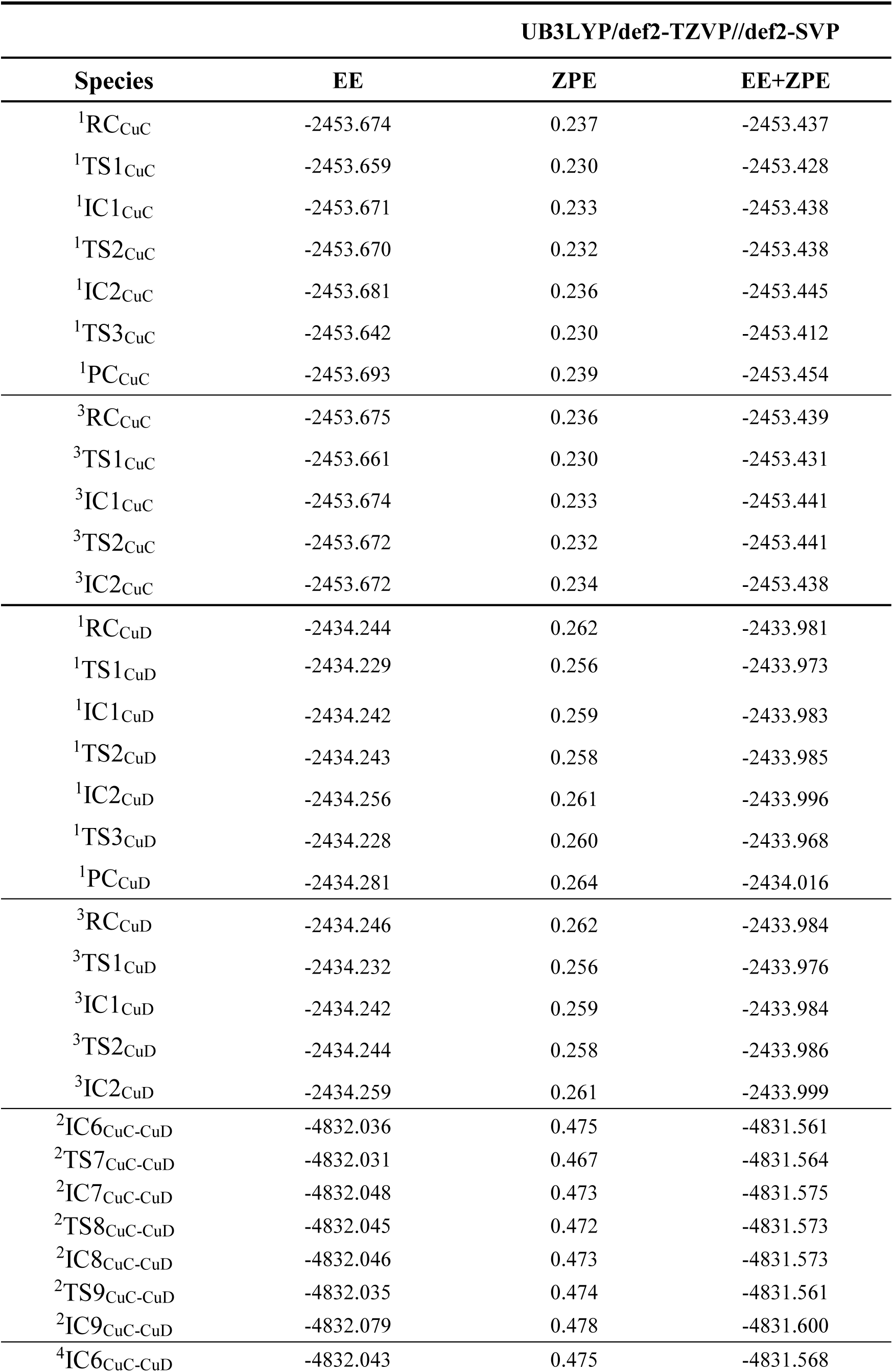
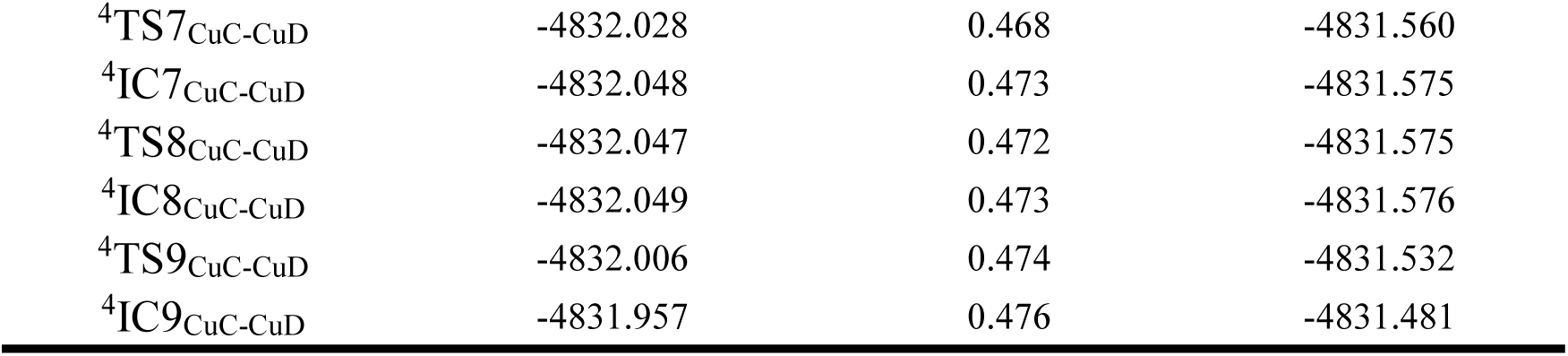
UB3LYP/def2-TZVP//def2-SVP QM energies (in Hartree, a.u.) of the species.

| UB3LYP/def2-TZVP//def2-SVP |  |  |  |
| --- | --- | --- | --- |
| Species | EE | ZPE | EE+ZPE |
| <sup>1</sup>RC<sub>CuC</sub> | -2453.674 | 0.237 | -2453.437 |
| <sup>1</sup>TS1<sub>CuC</sub> | -2453.659 | 0.230 | -2453.428 |
| <sup>1</sup>IC1<sub>CuC</sub> | -2453.671 | 0.233 | -2453.438 |
| <sup>1</sup>TS2<sub>CuC</sub> | -2453.670 | 0.232 | -2453.438 |
| <sup>1</sup>IC2<sub>CuC</sub> | -2453.681 | 0.236 | -2453.445 |
| <sup>1</sup>TS3<sub>CuC</sub> | -2453.642 | 0.230 | -2453.412 |
| <sup>1</sup>PC<sub>CuC</sub> | -2453.693 | 0.239 | -2453.454 |
| <sup>3</sup>RC<sub>CuC</sub> | -2453.675 | 0.236 | -2453.439 |
| <sup>3</sup>TS1<sub>CuC</sub> | -2453.661 | 0.230 | -2453.431 |
| <sup>3</sup>IC1<sub>CuC</sub> | -2453.674 | 0.233 | -2453.441 |
| <sup>3</sup>TS2<sub>CuC</sub> | -2453.672 | 0.232 | -2453.441 |
| <sup>3</sup>IC2<sub>CuC</sub> | -2453.672 | 0.234 | -2453.438 |
| <sup>1</sup>RC<sub>CuD</sub> | -2434.244 | 0.262 | -2433.981 |
| <sup>1</sup>TS1<sub>CuD</sub> | -2434.229 | 0.256 | -2433.973 |
| <sup>1</sup>IC1<sub>CuD</sub> | -2434.242 | 0.259 | -2433.983 |
| <sup>1</sup>TS2<sub>CuD</sub> | -2434.243 | 0.258 | -2433.985 |
| <sup>1</sup>IC2<sub>CuD</sub> | -2434.256 | 0.261 | -2433.996 |
| <sup>1</sup>TS3<sub>CuD</sub> | -2434.228 | 0.260 | -2433.968 |
| <sup>1</sup>PC<sub>CuD</sub> | -2434.281 | 0.264 | -2434.016 |
| <sup>3</sup>RC<sub>CuD</sub> | -2434.246 | 0.262 | -2433.984 |
| <sup>3</sup>TS1<sub>CuD</sub> | -2434.232 | 0.256 | -2433.976 |
| <sup>3</sup>IC1<sub>CuD</sub> | -2434.242 | 0.259 | -2433.984 |
| <sup>3</sup>TS2<sub>CuD</sub> | -2434.244 | 0.258 | -2433.986 |
| <sup>3</sup>IC2<sub>CuD</sub> | -2434.259 | 0.261 | -2433.999 |
| <sup>2</sup>IC6<sub>CuC-CuD</sub> | -4832.036 | 0.475 | -4831.561 |
| <sup>2</sup>TS7<sub>CuC-CuD</sub> | -4832.031 | 0.467 | -4831.564 |
| <sup>2</sup>IC7<sub>CuC-CuD</sub> | -4832.048 | 0.473 | -4831.575 |
| <sup>2</sup>TS8<sub>CuC-CuD</sub> | -4832.045 | 0.472 | -4831.573 |
| <sup>2</sup>IC8<sub>CuC-CuD</sub> | -4832.046 | 0.473 | -4831.573 |
| <sup>2</sup>TS9<sub>CuC-CuD</sub> | -4832.035 | 0.474 | -4831.561 |
| <sup>2</sup>IC9<sub>CuC-CuD</sub> | -4832.079 | 0.478 | -4831.600 |
| <sup>4</sup>IC6<sub>CuC-CuD</sub> | -4832.043 | 0.475 | -4831.568 |
| <sup>4</sup>TS7<sub>CuC-CuD</sub> | -4832.028 | 0.468 | -4831.560 |
| <sup>4</sup>IC7<sub>CuC-CuD</sub> | -4832.048 | 0.473 | -4831.575 |
| <sup>4</sup>TS8<sub>CuC-CuD</sub> | -4832.047 | 0.472 | -4831.575 |
| <sup>4</sup>IC8<sub>CuC-CuD</sub> | -4832.049 | 0.473 | -4831.576 |
| <sup>4</sup>TS9<sub>CuC-CuD</sub> | -4832.006 | 0.474 | -4831.532 |
| <sup>4</sup>IC9<sub>CuC-CuD</sub> | -4831.957 | 0.476 | -4831.481 |

**Extended Data Table 7.** Spin density population of key atoms for all species.

| Species |  | Spin density |  |  |  |
| --- | --- | --- | --- | --- | --- |
| QM/MM model | Cu<sub>D</sub> | Cu<sub>C</sub> | O1 | O2 | CoQ10 |
| <sup>3</sup>IC1<sub>CuD</sub> | 0.39 | - | 0.70 | 0.77 | 0.0 |
| <sup>3</sup>TS2<sub>CuD</sub> | 0.46 | - | 0.51 | 0.38 | 0.50 |
| <sup>3</sup>IC2<sub>CuD</sub> | 0.51 | - | 0.27 | 0.06 | 0.99 |
| <sup>1</sup>IC1<sub>CuD</sub> | -0.47 | - | 0.18 | 0.46 | 0.0 |
| <sup>1</sup>TS2<sub>CuD</sub> | -0.52 | - | 0.27 | 0.40 | 0.21 |
| <sup>1</sup>IC2<sub>CuD</sub> | -0.51 | - | -0.28 | -0.06 | 0.99 |
| <sup>2</sup>IC3<sub>CuC/CuD</sub> | 0.14 | 0.44 | 0.24 | 0.04 | - |
| <sup>2</sup>TS4<sub>CuC</sub> | 0.18 | 0.42 | 0.25 | 0.05 | - |
| <sup>2</sup>IC4<sub>CuC</sub> | 0.12 | 0.46 | 0.18 | 0.12 | - |
| <sup>2</sup>TS5<sub>CuC</sub> | 0.42 | 0.65 | -0.23 | -0.13 | - |
| <sup>2</sup>IC5<sub>CuC</sub> | 0.64 | 0.67 | -0.80 | 0.28 | - |
| <sup>2</sup>TS4<sub>CuD</sub> | 0.19 | 0.42 | 0.21 | 0.07 | - |
| <sup>2</sup>IC4<sub>CuD</sub> | -0.63 | 0.57 | 0.96 | -0.04 | - |
| <sup>2</sup>TS5<sub>CuD</sub> | -0.60 | 0.59 | 0.75 | 0.20 | - |
| <sup>2</sup>IC5<sub>CuD</sub> | 0.67 | -0.64 | 0.91 | 0.0 | - |
| QM model | Cu<sub>D</sub> | Cu<sub>C</sub> | O | Substrate |  |
| <sup>1</sup>RC<sub>CuC</sub> | - | -0.64 | 0.88 | -0.04 |  |
| <sup>1</sup>TS1<sub>CuC</sub> | - | -0.60 | 0.38 | 0.50 |  |
| <sup>1</sup>IC1<sub>CuC</sub> | - | -0.59 | -0.16 | 1.00 |  |
| <sup>1</sup>TS2<sub>CuC</sub> | -- | -0.56 | -0.16 | 0.95 |  |
| <sup>1</sup>IC2<sub>CuC</sub> | - | -0.59 | -0.19 | 1.10 |  |
| <sup>1</sup>TS3<sub>CuC</sub> | - | -0.28 | -0.08 | 0.30 |  |
| <sup>1</sup>PC<sub>CuC</sub> | - | 0.0 | 0.0 | 0.0 |  |
| <sup>3</sup>RC<sub>CuC</sub> | - | 0.54 | 1.19 | 0.0 |  |
| <sup>3</sup>TS1<sub>CuC</sub> | - | 0.56 | 0.69 | 0.56 |  |
| <sup>3</sup>IC1<sub>CuC</sub> | - | 0.59 | 0.20 | 1.00 |  |
| <sup>3</sup>TS2<sub>CuC</sub> | - | 0.59 | 0.20 | 0.95 |  |
| <sup>3</sup>IC2<sub>CuC</sub> | - | 0.59 | 0.19 | 1.10 |  |
| <sup>1</sup>RC<sub>CuD</sub> | -0.62 | - | 0.83 | 0.0 |  |
| <sup>1</sup>TS1<sub>CuD</sub> | -0.58 | - | 0.32 | 0.52 |  |
| <sup>1</sup>IC1<sub>CuD</sub> | -0.59 | - | -0.06 | 0.90 |  |
| <sup>1</sup>TS2<sub>CuD</sub> | -0.57 | - | -0.24 | 0.97 |  |
| <sup>1</sup>IC2<sub>CuD</sub> | -0.52 | - | 0.08 | 0.57 |  |
| <sup>1</sup>TS3<sub>CuD</sub> | -0.28 | - | 0.05 | 0.27 |  |
| <sup>1</sup>PC<sub>CuD</sub> | 0.0 | - | 0.0 | 0.0 |  |
| <sup>3</sup>RC<sub>CuD</sub> | 0.50 | - | 1.24 | 0.0 |  |
| <sup>3</sup>TS1<sub>CuD</sub> | 0.54 | - | 0.73 | 0.56 |  |
| <sup>3</sup>IC1<sub>CuD</sub> | 0.59 | - | 0.31 | 0.91 |  |
| <sup>3</sup>TS2<sub>CuD</sub> | 0.59 | - | 0.24 | 0.97 |  |

| <sup>3</sup>IC2<sub>CuD</sub> | 0.59 | - | 0.30 |  | 0.85 |
| --- | --- | --- | --- | --- | --- |
| **QM model** | **Cu<sub>D</sub>** | **Cu<sub>C</sub>** | **O1** | **O2** | **Substrate** |
| <sup>2</sup>IC6<sub>CuC-CuD</sub> | -0.58 | 0.65 | 1.18 | 0.01 | 0.0 |
| <sup>2</sup>TS7<sub>CuC-CuD</sub> | -0.58 | 0.62 | 0.52 | 0.12 | 0.52 |
| <sup>2</sup>IC7<sub>CuC-CuD</sub> | -0.64 | 0.64 | 0.21 | 0.01 | 0.97 |
| <sup>2</sup>TS8<sub>CuC-CuD</sub> | -0.66 | 0.64 | 0.03 | 0.11 | 0.97 |
| <sup>2</sup>IC8<sub>CuC-CuD</sub> | -0.64 | 0.62 | 0.22 | 0.11 | 0.94 |
| <sup>2</sup>TS9<sub>CuC-CuD</sub> | -0.38 | 0.64 | 0.20 | 0.11 | 0.67 |
| <sup>2</sup>IC9<sub>CuC-CuD</sub> | 0.0 | 0.62 | 0.0 | 0.11 | 0.0 |
| <sup>4</sup>IC6<sub>CuC-CuD</sub> | 0.58 | 0.65 | 1.18 | 0.18 | 0.0 |
| <sup>4</sup>TS7<sub>CuC-CuD</sub> | 0.59 | 0.65 | 0.59 | 0.18 | 0.52 |
| <sup>4</sup>IC7<sub>CuC-CuD</sub> | 0.66 | 0.65 | 0.23 | 0.01 | 1.00 |
| <sup>4</sup>TS8<sub>CuC-CuD</sub> | 0.65 | 0.66 | 0.23 | 0.11 | 0.96 |
| <sup>4</sup>IC8<sub>CuC-CuD</sub> | 0.65 | 0.67 | 0.22 | 0.11 | 0.93 |
| <sup>4</sup>TS9<sub>CuC-CuD</sub> | 0.67 | 0.64 | 0.29 | 0.13 | 0.89 |
| <sup>4</sup>IC9<sub>CuC-CuD</sub> | 0.64 | 0.64 | 0.0 | 0.19 | 0.0 |

**Extended Data Table 8.**
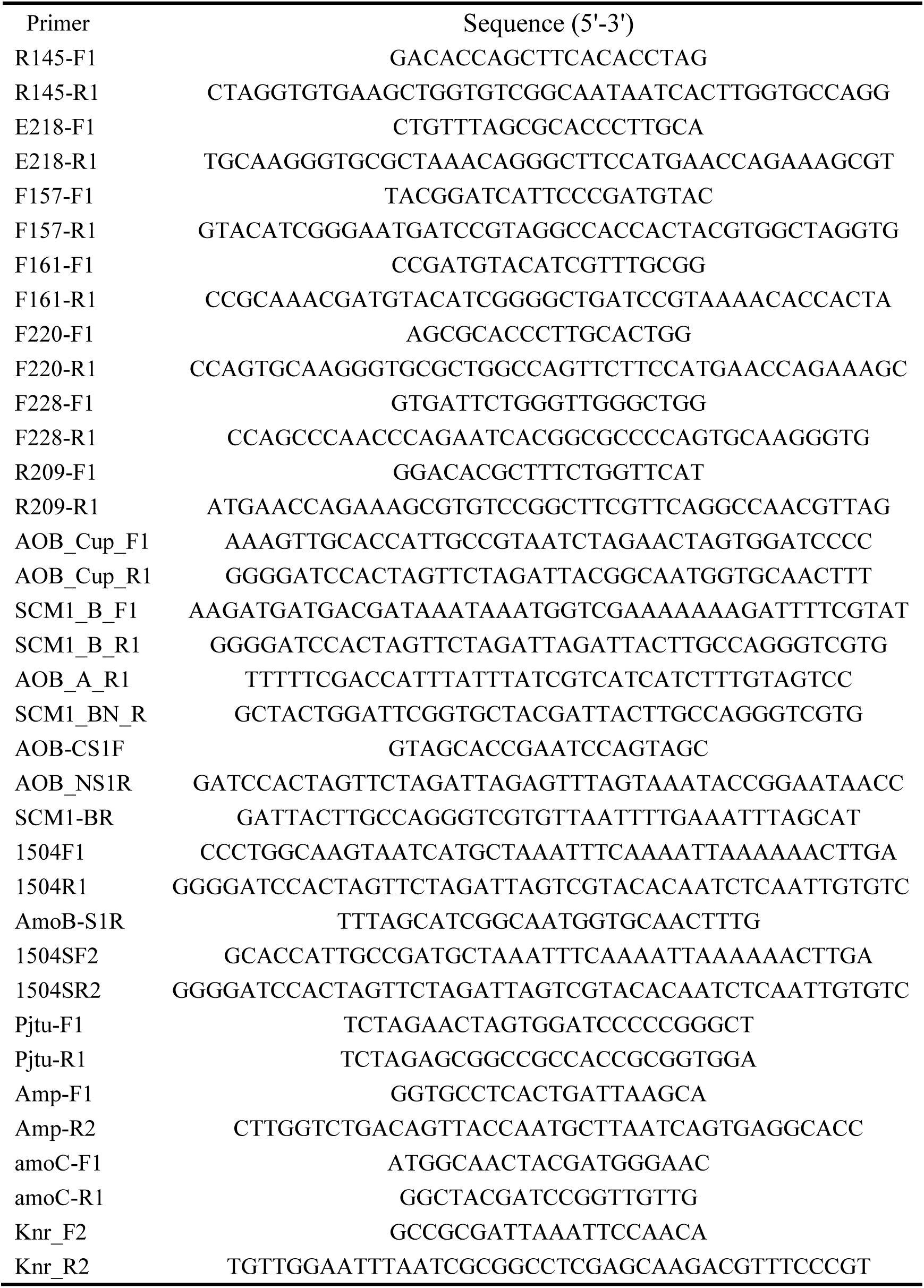
Oligonucleotide primers used in this study.

